# Loss-of-Function RUP-Variants Influence Ubiquitin-Proteasome System and Enhance Carotenoid and Folate Levels in Tomato

**DOI:** 10.1101/2025.04.17.649262

**Authors:** Chaitanya Charakana, Suresh Kumar Gupta, Anusha Sunkum, Satyavathi Valluri, Jayram Bagri, Kamal Tyagi, Rakesh Kumar, Hymavathi Salava, Himabindu Vasuki Kilambi, Venkataramana Sivapuram, Rameshwar Sharma, Yellamaraju Sreelakshmi

**Author notes:** **Corresponding authors:** (R. Sharma); (Y. Sreelakshmi). **Author’s E-mails:** (CC); (SKG); (AS); (SV); (JB); (KT); (RK); (HS); (HVK); (VS). Equal contribution 2^nd^ Author (Suresh Kumar Gupta, Anusha Sunkum). Equal contribution 5^th^ Author (Kamal Tyagi, Rakesh Kumar, Hymavathi Salava). **Permanent address:**. Department of Life Science, School of Life Sciences, Central University of Karnataka, Kalaburagi, Karnataka, 585367 (Rakesh Kumar). School of Biological Sciences, National Institute of Science Education and Research, PO Bhimpur-Padanpur, 752050, Odisha, India.

## Abstract

REPRESSOR OF UV-B PHOTOMORPHOGENESIS (RUP) negatively regulates UV- B signalling, yet its broader physiological roles in crop plants remain largely unexplored. We compared two tomato accessions bearing truncated RUP proteins—*rup-1* and *rup-2* (*rup-* variants)—with the cultivar Arka Vikas (AV), which harbours the native RUP protein. Seedlings of *rup*-variants exhibited enhanced tolerance to supplemental UV-B light. Red ripe (RR) fruits of *rup*-variants showed significantly elevated carotenoid and folate levels compared to AV. Introgression of *rup*-variants into AV confirmed that the increased carotenoid accumulation is genetically linked to RUP truncation. Metabolomic profiling of *rup*-variants revealed a substantial shift in primary metabolic homeostasis, particularly at the breaker stage, marked by a pronounced reduction in sugars and amino acids. Proteomic analyses of *rup*- variants across ripening stages identified that a significant proportion of differentially expressed proteins belonged to chaperones and ubiquitin-proteasome system (UPS). Upregulation of four key enzymes in the carotenoid biosynthesis pathway likely contributed to increased lycopene content in *rup*-variants. Elevated folate levels in *rup*-variants were associated with the upregulation of folate biosynthesis and C1 metabolism enzymes. Despite widespread metabolic reprogramming in *rup*-variants, hormonal regulatory pathways remained largely unaltered. Our results suggest that RUP modulates metabolic pathways during fruit ripening, and its loss triggers metabolic reprogramming associated with elevated folate/carotenoid levels.

## Introduction

Ripening of fruits involves a major cellular and metabolic transition, transforming fruits from unpalatable to palatable with aroma. In tomato (*Solanum lycopersicum*), fruit ripening is primarily governed by the genetic makeup but also modulated by environmental factors. Light is one such environmental factor that influences several facets of tomato ripening. During early fruit development, light provides nearly 20% of the carbohydrate requirement through photosynthesis within the fruits (**Tanaka et al., 1974; Hetherington et al., 1998**). The role of light is not limited to photosynthesis during tomato fruit growth; at the onset and during ripening, several metabolic processes are modulated by light. Initial studies indicated that different spectral regions of light regulate different aspects of tomato ripening. Illumination of tomato fruits with different lights revealed that red light was most effective in promoting fruit softening during early ripening, while blue light promoted the red hue development in later ripening stages (**Jen, 1974**).

Extensive studies on light-regulated morphogenesis in higher plants have demonstrated that plants perceive the spectral quality, duration, direction, and intensity of light through multiple photoreceptors (**Paik and Huq, 2019; Kharshiing et al., 2019**). These photoreceptors sense different spectral regions of light; phytochromes primarily sense red and far-red light (600–750 nm); cryptochromes, phototropins, and F-box flavin-binding proteins (FKF1, LKP2, and Zeitlupe) detect blue/UV-A light (320–500 nm); and UV RESISTANCE LOCUS 8 (UVR8) is specialized for UV-B perception (280–320 nm). The studies involving mutants established that in addition to regulating plant growth and morphogenesis, phytochromes, phototropin1, and cryptochrome2 modulate tomato fruit ripening, such as accumulation of lycopene (**Gupta et al., 2014**; **Kilambi et al., 2021; Fantini et al., 2019**). Similarly, transgenic manipulation of the expression of phytochromes, cryptochromes, and UVR8 affects fruit ripening, including lycopene levels (**Li et al., 2018**; **Fantini et al., 2019; Bianchetti et al., 2020).**

Molecular genetic analyses of Arabidopsis light-signalling mutants have established CONSTITUTIVELY PHOTOMORPHOGENIC 1 (COP1) as a central regulator of light-mediated development, functioning through interactions with multiple photoreceptors to modulate plant growth responses (**Ponnu and Hoecker, 2021**). In the darkness, COP1 forms an E3 ubiquitin ligase complex with SUPPRESSOR OF PHYA-105 (SPA) proteins, targeting photomorphogenesis-promoting transcription factors such as HY5 for proteasomal degradation. Upon red light exposure, activated phytochromes interact with SPA proteins, leading to the destabilization of the COP1-SPA complex, allowing the accumulation of transcription factors promoting photomorphogenesis **(Sheerin et al., 2015).** In contrast to phytochromes, blue light-activated cryptochromes directly bind to COP1, inhibiting its E3 ligase activity and thereby stabilizing positive regulators of light signalling **(Wang et al., 2000; Liu et al., 2011).**

Similar to cryptochromes, UVR8 directly interacts with COP1 to mediate light signalling. In darkness, UVR8 exists as a homodimer; upon UV-B exposure, it undergoes monomerization and subsequently interacts with COP1 (**Liu and Jenkins, 2025**). This interaction involves the β-propeller domain of UVR8, and the WD40 repeat domain of COP1, with the C-terminal C27 domain of UVR8 playing a regulatory role in modulating COP1 activity (**Yin et al., 2015**). Unlike its function in visible light signalling, where COP1 acts as a repressor, COP1 serves as a positive regulator of UV-B–induced photomorphogenesis (**Oravecz et al., 2006**). The UVR8-COP1 complex promotes the expression of UV-B-responsive genes by dissociating the COP1– SPA complex from the CUL4-DDB1 E3 apparatus, leading to photomorphogenic responses and stress acclimation. The inactivation of the UVR8 is facilitated by its dimerization by WD40-repeat proteins RUP1 (*REPRESSOR OF UV PHOTOMORPHOGENESIS)* and RUP2, which interact with the C27 domain of UVR8, effectively turning off the UV-B signalling (**Heijde and Ulm, 2013).**

While RUPs serve as repressors to modulate the UVR8-mediated signalling, these can also function independently of the UV-B signalling pathway. During Arabidopsis seed germination, RUP1 and RUP2 physically interact with ABA INSENSITIVE 5 and suppress ABA-mediated inhibition of germination (**Singh et al., 2024**). RUP1 and RUP2 are also components of CUL4-DDB1-based E3 ubiquitin ligase, which targets HY5, a key transcription factor in photomorphogenesis for degradation. COP1 interacts directly with RUP1 and RUP2 via its WD40 domain, promoting their degradation to facilitate HY5 expression (**Ren et al., 2019**). Moreover, RUP1/RUP2 also regulate HY5 stability independently of UVR8 (**Podolec et al., 2021**). RUP2 also physically interacts with CONSTANS (CO), a major floral inducer, and represses its binding to the promoter of *FT*, the gene encoding florigen. Seemingly, RUP1 and RUP2 have multifunctional roles in plant development. Their multifunctional role is also supported by the enhancement of *RUP1* and *RUP2* expression by cryptochrome and phytochromes A in Arabidopsis seedlings (**Tissot and Ulm, 2020**).

In contrast to Arabidopsis, the tomato possesses a single copy of the *RUP* gene. Silencing or overexpression of *UVR8*, the main target of *RUP*, elicited the opposite fruit phenotype; *UVR8* RNAi (*UVR8^Ri^*) lines produce pale-green fruits, whereas UVR8 overexpression (*UVR8^OE^*) leads to dark green fruits with elevated lycopene content compared to the wild type (**Li et al., 2018**). Similar to Arabidopsis, tomato RUP inhibits the nuclear retention of UVR8, as nuclear levels of UVR8 were much higher in a *rup*-mutant than in the WT (**Fang et al., 2022**). Likewise, RUP also promotes the re-dimerization of tomato UVR8 monomers (**Zhang et al., 2021**).

While the role of RUP protein in UVR8 signalling in tomato is broadly similar to Arabidopsis (**Liu et al., 2020**), its specific contribution to tomato fruit metabolism remains largely unexplored. In this study, we examined two tomato accessions harbouring single nucleotide polymorphisms (SNPs) in the *RUP* gene that result in truncated RUP proteins. Fruits from plants carrying the *rup*-variant protein exhibited significantly elevated levels of carotenoids and folates. Genetic analysis established a strong linkage between the *rup-*variant alleles and increased carotenoid accumulation. Proteomic profiling across fruit ripening stages further indicated that alterations in protein protection and degradation pathways underlie the upregulation of carotenoid and folate biosynthesis in *rup*-variant fruits.

## Results

### Onset of ripening elevates *RUP* expression in tomato fruit

In tomato, photoreceptors and their downstream signalling partners play a significant role in fruit ripening (**Kilambi et al., 2013, 2021; Gupta et al., 2014**). To identify new signalling partners, we analysed public transcriptome databases for light-signalling-associated genes that are upregulated during tomato fruit development and ripening. Our analysis identified the gene *REPRESSOR OF UV PHOTOMORPHOGENESIS* (*SlRUP;* Solyc11g005190), which is upregulated from the immature green (IMG) stage to the late breaker (BR) stage of ripening (**Figure 1**). Unlike Arabidopsis, the tomato has a single RUP protein that shares 55.5% and 56.1% homology with AtRUP1 and AtRUP2, respectively (**Zhang et al., 2021**) and is phylogenetically closer to Arabidopsis RUP1 (**Figure S1; Yan et al., 2023**).

**Figure 1.**
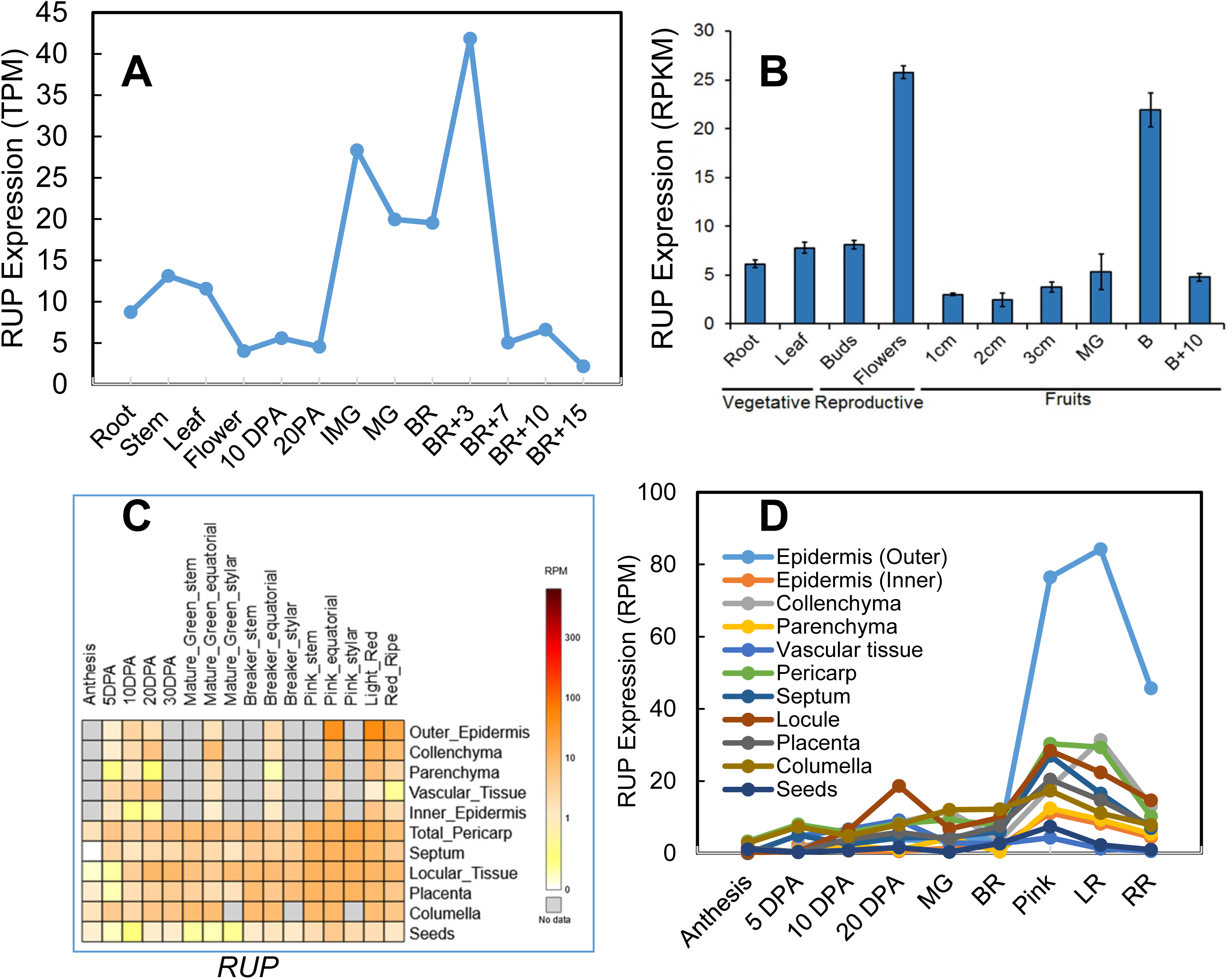
*RUP* (*Solyc11g005190*) gene expression in different organs and ripening fruits of tomato. **A.** Data Source-Table S1: Yan C, Yang T, Wang B, Yang H, Wang J, Yu Q. (2023) Genome-Wide Identification of the WD40 Gene Family in Tomato (*Solanum lycopersicum L*.). Genes. **14:** 1273. **B.** Data source-Tomato Genome Consortium (2012). The tomato genome sequence provides insights into fleshy fruit evolution. *Nature*, **485:** 635. **C-D.** Data source-https://tea.solgenomics.net. Abbreviations: DPA-Days post-anthesis; MG-Mature green; BR-Breaker; LR-Light red; RR-Red ripe; BR+#-Days from breaker.

The tomato *RUP* gene is intronless, spans 1,083 base pairs, and encodes a 360-amino-acid protein belonging to the WD-repeat protein family with seven WD40 repeats. Spatiotemporal analysis of *RUP* expression in fruit tissues revealed high expression levels in the pericarp and adjacent tissues, with a marked increase in expression in the outer epidermis during ripening (**Fernandez-Pozo et al., 2017; Shinozaki et al., 2018**) (**Figure 1**). Interestingly, *RUP* expression does not correlate with *UVR8* (Solyc05g018620), which expresses nearly at a similar level throughout the fruit development (**Figure S2**).

### Loss of RUP protein elevates carotenoid levels in tomato fruits

An earlier study showed that reducing *RUP* (*LeCOP1like*, AW625993) expression via RNA interference resulted in tomato plants with exaggerated photomorphogenesis and increased carotenoid levels in the fruit (**Liu et al., 2004**). Given RUP’s potential role in modulating carotenoid levels, we screened 391 tomato accessions for single nucleotide polymorphisms (SNPs) in *RUP* using EcoTILLING (**Mohan et al., 2016; Sarma et al., 2018**). Our analysis identified 20 SNPs distributed across four accessions. Based on SNP diversity, we designated these accessions as *rup-1* (EC398697), *rup-2* (EC529085), *rup-3* (EC2790), and *rup-4* (EC398716), collectively referred to as *rup*-variants (**Table S1**). We selected the Arka Vikas (AV) as a control cultivar to compare with *rup*-variants based on the absence of SSR markers polymorphism with *rup-1* and *rup-2* (**Figure S3**).

In *rup-1* and *rup-2*, insertions caused frameshift mutations, resulting in protein truncation. In *rup-1*, a C489CC insertion (D164R) led to premature termination of the protein at R164*. Similarly, the T456TT insertion in *rup-2* altered the protein sequence after C152 and terminated it at R164* (**Figure S4**). While *rup-1* contained five SNPs and *rup-2* had an additional two insertions and two SNPs, these were downstream of the truncation site and thus considered redundant. The *rup-3* and *rup-4* contained seven (four synonymous, three nonsynonymous) and two (one synonymous, one nonsynonymous) SNPs, respectively (**Figure S4**). According to DDMut and SIFT analyses, the nonsynonymous SNPs in *rup-3* are predicted to be tolerated and unlikely to affect RUP function. For *rup-4,* results for nonsynonymous SNP were mixed: DDMut predicted a destabilizing effect, while SIFT classified the variant as tolerated (**Table S1**).

Preliminary spectrophotometric analysis of red ripe (RR) fruits showed elevated total carotenoid levels in all *rup* variants relative to AV: *rup-1* (2.15-fold), *rup-2* (2.3-fold), *rup-3* (2.17-fold), and *rup-4* (1.74-fold). This feature was comparable to *cop1like* RNAi lines [*cop1like* RNAi-A (1.34-fold) and RNAi-B (1.6-fold)] (**Figure S5**). Given that *rup-1* and *rup-2* have high carotenoid levels and bear protein truncation, we selected *rup-1* and *rup-2* for detailed analysis. Detailed UHPLC analysis of the complete carotenoid profile of RR fruits of *rup-1* and *rup-2* revealed that *rup-1* (1.67-fold) and *rup-2* (1.52-fold) had higher carotenoid levels compared to AV (**Table 1**). The above increase was primarily due to elevated levels of lycopene (1.76-fold *rup-1*; 1.69-fold *rup-2*), phytoene (1.83-fold *rup-1*; 1.40-fold *rup-2*), and also β-carotene, and lutein, particularly during the transition from the breaker (BR) to the RR stage. In addition to carotenoids, *rup-1* and *rup-2* fruits also showed significantly higher levels of folate vitamers compared to AV (**Figure S6**). Although total folate levels were elevated, the proportional distribution of individual vitamers remained similar to that in AV.

**Table 1.**
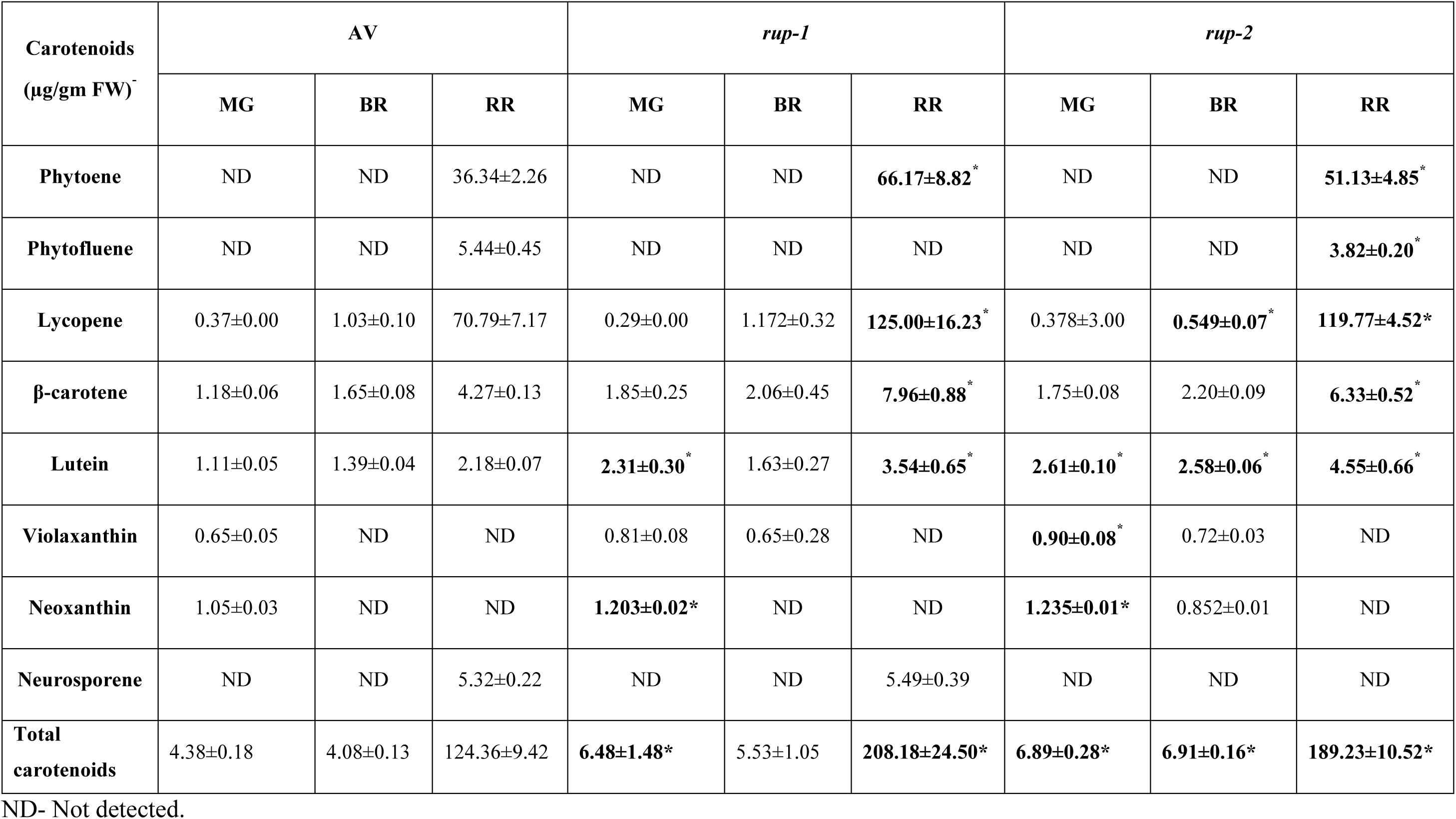
Carotenoid levels in fruits of Arka Vikas (AV) and *rup*-variants at different stages of ripening. The significantly changed carotenoids at the respective stages are highlighted with bold letters. The carotenoid data are mean ± SE (n ≥ 3). Statistical significance is represented by * for P value ≤ 0.05.

### Influence of *rup* alleles on the phenotype of tomato plants

Seedlings of CRISPR-knockout *RUP* mutants of tomato grown under white light exhibit photomorphogenic development similar to wild type (WT) but display increased resistance to supplemental UV-B light compared to WT (**See Supplementary Figure 6 of Zhang et al., 2021**). In agreement with this, *rup-1* and *rup-2* seedlings also showed higher resistance to supplemental UV-B compared to AV (**Figure 2A, Figure S7**). Comparative analysis of dark-and white-light-grown seedlings revealed that while AV and *rup-2* appeared phenotypically similar, *rup-1* seedlings were distinctly different. Specifically, *rup-1* showed shorter hypocotyls under both dark and light conditions and reduced root length in the dark(**Figure 2B-E, Figure S8**). In adult plants, *rup-1* also displayed higher chlorophyll A levels in the leaves compared to *rup-2* and AV. Although the overall vegetative morphology of *rup*-variants and AV was comparable, the fruits of *rup-1* and *rup-2* were smaller than those of AV **(Figure 2F**, **Figure S9-S10**).

**Figure 2.**
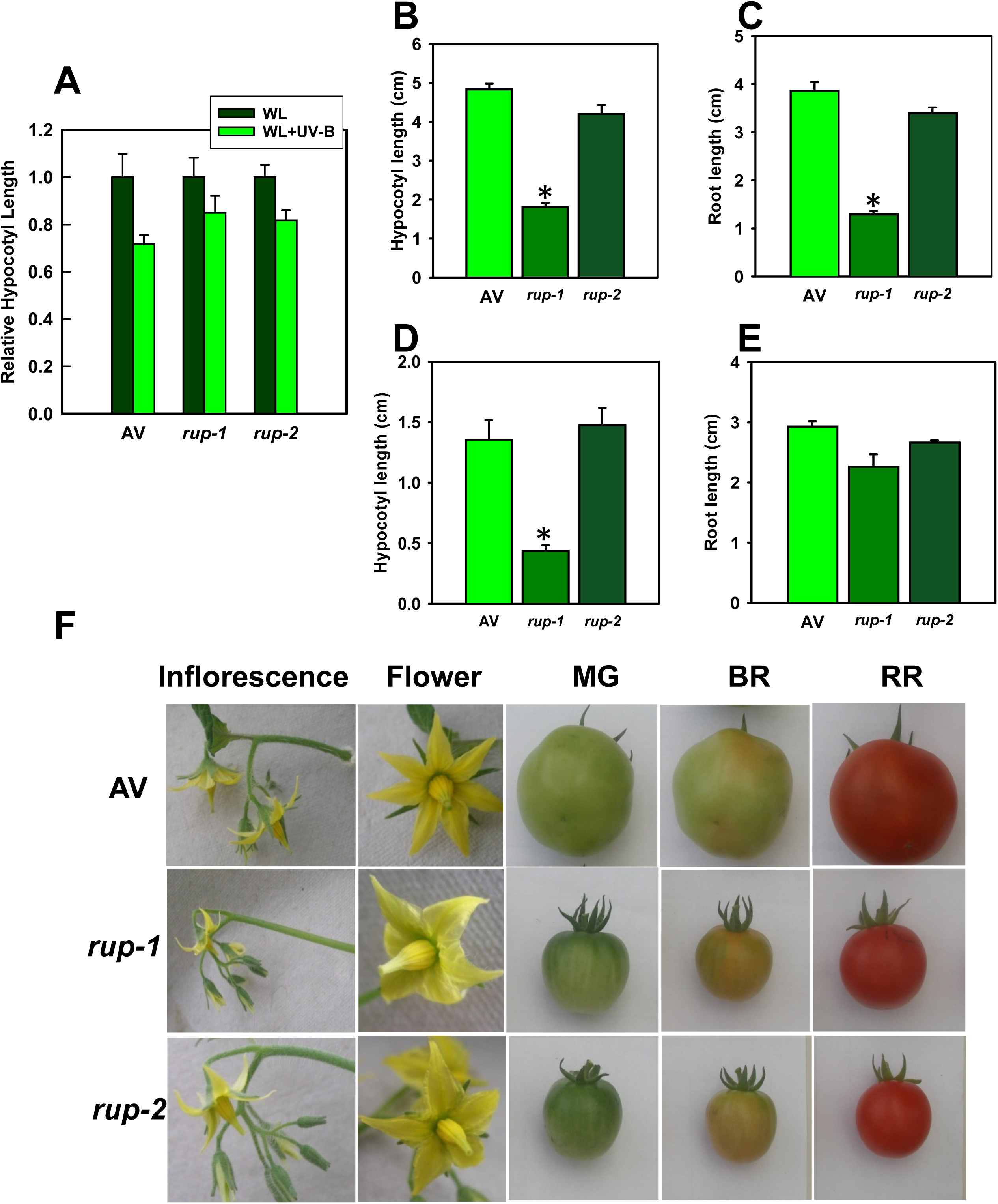
Vegetative and reproductive phenotypes of AV and *rup-*variants. **A.** Hypocotyl length of seedlings (7-day-old) grown under white light (WL). A set of seedlings was exposed daily to UV-B for 4-h. **B-E.** Hypocotyl and root lengths of 3-day-old dark-grown (**B-C**) and light-grown seedlings (**D-E**) of AV and *rup*-variants. **F.** Flower and fruit morphology of *rup-1* and *rup-2* compared to AV. Fruits of the *rup*-variants are significantly smaller than AV fruits at all developmental stages: Mature green (**MG**), Breaker (**BR**), and Red Ripe (**RR**). Each histogram is a representation of the mean value ±SE (n ≥ 3). Asterisks indicate significant differences with P < 0.05 (*).

### Introgression of *rup*-variants elevates carotenoid levels in Arka Vikas fruits

To confirm that the elevated carotenoid level in RR fruits of *rup*-variants was genetically linked to RUP truncation, *rup-1* and *rup-2* were crossed with AV. The F_2_ seedlings segregated for the *rup-1* gene in a Mendelian ratio, and plants were grown to maturity (**Figure S11**). Analysis of RR fruits of F_2_ plants showed a genetic linkage between carotenoid levels and RUP truncation, with homozygous *rup-1/rup-1* and *rup-2/rup-2* plants displaying higher carotenoid levels than AV (*RUP/RUP*) (**Figure S12**). Homozygous BC_1_F_2_ (*rup-1/rup-1* and *rup-2/rup-2*) plants were advanced to BC_4_F_2_ generation in the AV background. The RR fruits of *rup-1/rup-1* and *rup-2/rup-2* BC_4_F_2_ plants revealed higher carotenoid levels like *rup*-variants parental lines, confirming that introgression of *rup* alleles increased carotenoid accumulation in AV (**Figure S12-S13, Dataset S1**). Similar to the respective *rup* parent lines, elevation in carotenoids in *rup/rup* BC_4_F_2_ fruits was primarily due to higher levels of lycopene, phytoene, β-carotene, and lutein compared to AV (**Figure S13**).

### Reduced *lycopene β-cyclase* expression may contribute to high carotenoid levels in *rup* fruits

Although *rup*-variants displayed higher carotenoid levels in fruits compared to AV, the expression of *DXS*, a key gene involved in the synthesis of the carotenoid precursor geranyl-geranyl diphosphate (GGPP), was nearly similar to AV. Additionally, no significant variation was observed in the expression of key carotenoid biosynthesis genes *—PSY1*, *PDS*, *ZISO*, *ZDS*, and *CRITSO*—relative to AV. However, the expression of *CYCB*, which is involved in the conversion of lycopene to β-carotene, was lower in both *rup*-variants. Notably, the expression of the *CHRC,* a gene contributing to carotenoid sequestration, was significantly upregulated in *rup*-variants, whereas the expression of *PAP3* remained similar to AV (**Figure S14**). Furthermore, the *rup*-variants did not alter the expression of light-signalling genes such as *COP1* and *HY5*. The only exception was *SPA1*, which was downregulated during ripening (**Figure S15**).

### The metabolic shift in *rup*-variants is confined mainly to the BR stage

The increased folate and carotenoid levels in RR fruits indicated that *rup-*variants influence the overall metabolic homeostasis during fruit ripening. Profiling of primary metabolites during fruit ripening identified 82 metabolites in *rup-1*, *rup-2*, and AV. Partial Least-Squares Discriminant Analysis (PLS-DA) revealed that metabolically, *rup-1*, *rup-2*, and AV were closely aligned at the mature green (MG) and red-ripe (RR) stages. However, a distinct separation was observed at the breaker (BR) stage, where the PLS-DA profile of the *rup*-variants diverged significantly from AV across all three component axes (**Figure 3A**).

**Figure 3.**
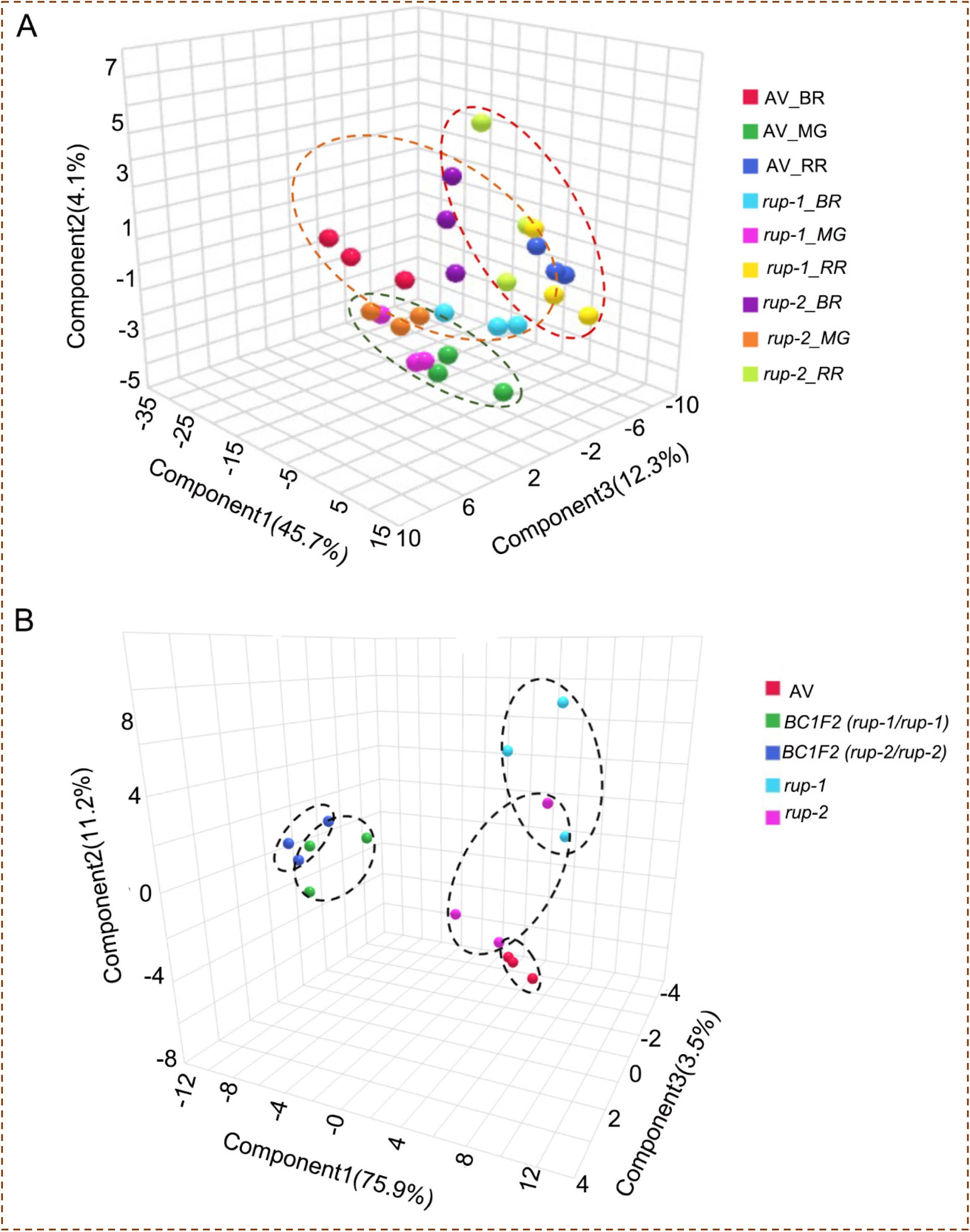
Partial Least-Squares Discriminant Analysis (PLS-DA) of metabolites of AV and *rup-*variants fruits. (A) PLS-DA plots of AV, *rup-1,* and *rup-2* along with AV at different ripening stages of fruits. Each ripening stage is encircled by a dashed line oval. MG, mature green (Green Oval); BR, breaker (Orange Oval); RR, red ripe (Red Oval). (B). PLS-DA plot of RR fruits of AV and respective *rup*-variants, along with BC_1_F_2_ *(rup-1/rup-1)* and BC_1_F_2_ *(rup-2/rup-2)* plants, where the *rup* gene is present in homozygous condition. The PLS-DA was constructed using MetaboAnalyst 5.0. The variance of the components C1, C2, and C3 are given within parentheses. See **Dataset S2-S3** for details.

Analysis of individual metabolite profiles revealed that the majority were downregulated (↓), with relatively few upregulated (↑), in the *rup*-variants compared to AV. In *rup-1*, 40 out of 82 metabolites showed significant changes (MG: 3↓, 5↑; BR: 26↓, 0↑; RR: 9↓, 6↑), while in *rup-2*, 35 metabolites significantly changed (MG: 8↓, 3↑; BR: 14↓, 0↑; RR: 9↓, 5↑) relative to AV (**Figure 4A, Dataset S2**). Consistent with the PLS-DA results, the most pronounced shift occurred at the BR stage, where all significantly altered metabolites were downregulated in *rup-1* (26↓) and *rup-2* (14↓). A notable feature at the BR stage was a substantial reduction in sugar levels. Of the 19 sugars detected, 14 in *rup-1* and 8 in *rup-2* were lower than in AV. Several metabolites exhibited shared regulation patterns in *rup-1* and *rup-2* during ripening (MG: 3↓, 2↑; BR: 12↓; RR: 6↓, 2↑). A key metabolite of the TCA cycle, citrate, was consistently downregulated across all ripening stages in *rup-1,* and during the MG and BR stages of *rup-2* (**Figure S16**). Nevertheless, some metabolites showed variant-specific regulation, such as increased fatty acid levels at the MG and higher methyl maleate and malate levels at the RR stage in *rup-2*.

**Figure 4.**
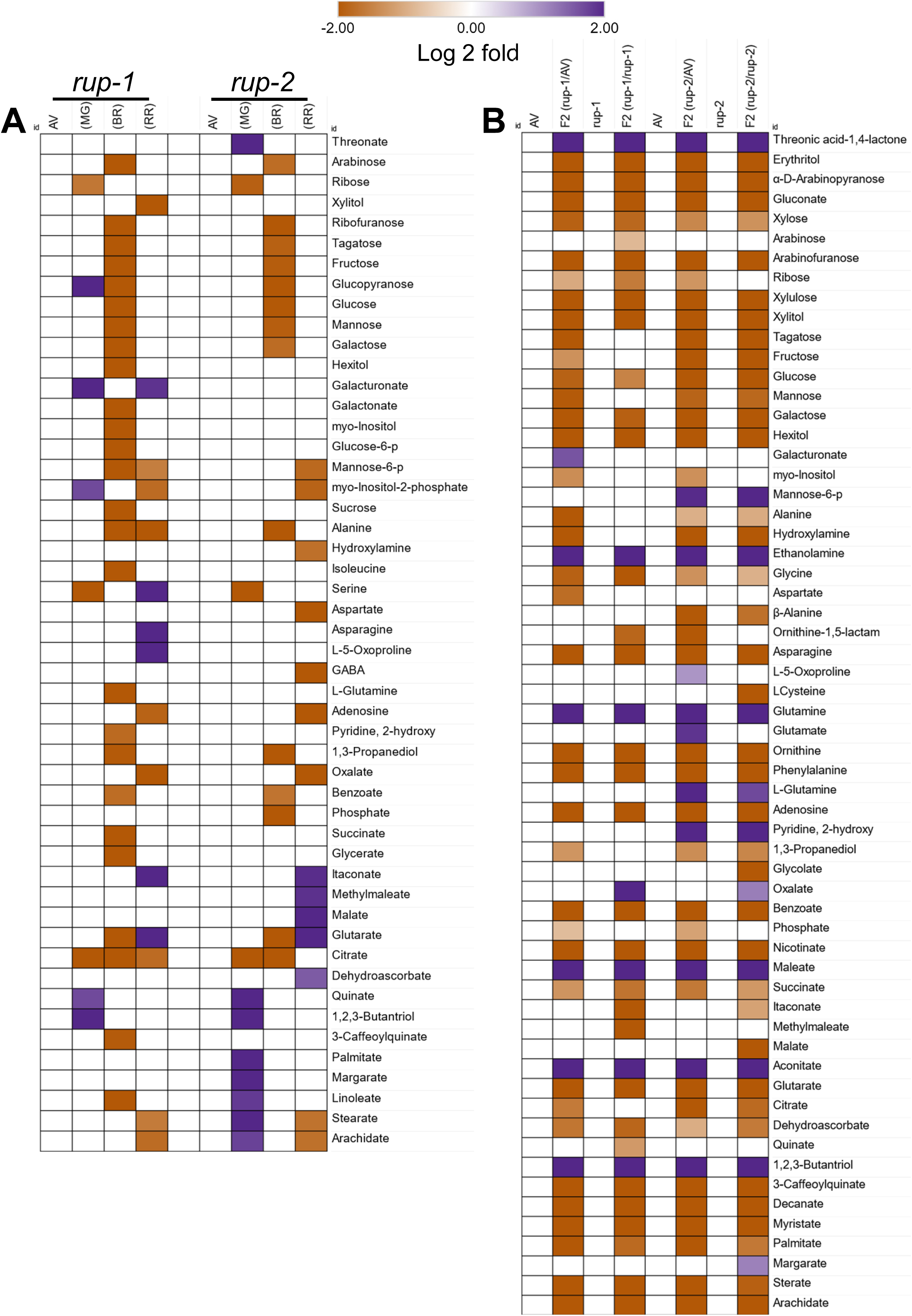
Relative metabolite profiles of *rup-*variant fruits. **A.** The relative metabolite profiles of *rup-1* and *rup-2* fruits compared to AV fruits at MG, BR, and RR stages of ripening. **B.** The relative metabolite profiles of RR fruits of BC_1_F_2_ (*rup-1/rup-1*) and BC_1_F_2_ (*rup-2/rup-2*) plants, where the *rup* gene is present in homozygous condition. The relative changes were determined with respect to AV and *rup* parents. Only significantly changed metabolites, with reference to respective control, are depicted (Log2 fold ≥ ± 0.584, p-value ≤ 0.05, n ≥ 3). See **Dataset S2-S3 for** metabolite levels and significance.

### Introgression of *rup*-variants lowers metabolite levels in the progeny fruits

To identify metabolites regulated by *rup-*variants, RR fruits of AV (*RUP*/*RUP*), *rup-1* (*rup-1*/*rup-1*), *rup-2* (*rup-2/rup-2*), *rup-1* homozygous BC_1_F_2_ (*rup-1/rup-1*), and *rup-2* homozygous BC_1_F_2_ (*rup-2/rup-2*) were analysed. PLS-DA analysis revealed distinct metabolic profiles for BC_1_F_2_ *rup-1/rup-1* and *rup-2/rup-2*, distinguishing them from parental lines (**Figure 3B**). Out of 68 metabolites detected in RR fruits of AV and *rup-*variants, only 51 were present in BC_1_F_2_ *rup-1/rup-1* and BC_1_F_2_ *rup-2/rup-2* fruits. Additionally, six new stress-related metabolites, including ethanolamine, glutamine, aconitate, and maleate, were identified in the BC_1_F_2_ fruits of *rup-1/rup-1* and *rup-2/rup-2*. The absence of 17 metabolites in BC_1_F_2_ *rup-1/rup-1* and *rup-2/rup-2* fruits, compared to parental lines, indicates that these metabolites were suppressed in the mixed genetic backgrounds of AV and *rup*-variants. Given the similarity in the metabolite profiles of BC_1_F_2_ *rup-1/rup-1* and *rup-2/rup-2* fruits, it appears that both *rup-1* and *rup-2* influence RR fruit metabolites in a comparable manner.

Consistent with PLS-DA, the metabolome of BC_1_F_2_ *rup-1/rup-1* and *rup-2/rup-2* fruits exhibited extensive downregulation compared to their respective parents. In BC_1_F_2_ *rup-1/rup-1* fruits, 33 metabolites showed similar changes relative to AV and *rup-1* (6**↑**, 27**↓**) (**Figure 4B, Dataset S3**). Additionally, other metabolites in BC_1_F_2_ *rup-1/rup-1* were uniquely downregulated with respect to AV (10**↓**, 1**↑**) or *rup-1* (5**↓**, 1**↑**). The metabolome of BC_1_F_2_ *rup-2/rup-2* fruits displayed even more pronounced downregulation, with 43 metabolites commonly altered compared to AV and *rup-2* (9**↑**, 34**↓**). In addition, the BC_1_F_2_ *rup-2/rup-2* fruits showed further metabolomic shifts with respect to only AV (4**↓**, 2**↑**) and *rup-2* (2**↓**, 2**↑**). Among the metabolite classes, most sugars, including galactose, glucose, and xylose, were downregulated in *rup-1/rup-1* and *rup-2/rup-2* progeny. Similarly, amino acids such as glycine and ornithine were reduced. TCA cycle intermediates like succinate and dehydroascorbate and fatty acids such as palmitate and stearate were also lowered. Overall, the metabolomes of BC_1_F_2_ *rup-1/rup-1* and *rup-2/rup-2* fruits exhibited extensive metabolic downregulation relative to their progenitor parents.

### *rup*-variants have higher amounts of carotenoid biosynthesis proteins

To decipher the molecular basis of elevated carotenoids and folate levels in *rup*-variant fruits, we compared the proteomes of *rup*-variants with that of AV (**Dataset S4**). Out of ≥ 2500 detected proteins (**Figure S17A**), nearly ≥ 20% of proteins were differentially regulated. The differentially expressed proteins consisted of either uniquely regulated in *rup-1* and *rup-2* [*rup-1* (MG: 121**↑**, 153**↓**; BR: 125**↑**, 195**↓**; RR: 183**↑**, 90**↓**); *rup-2* (MG: 131**↑**, 100**↓**; BR: 148**↑**, 152; RR: 189**↑**, 109**↓**)] or overlapped between *rup-1* and *rup-2* (MG; 189; BR: 269; RR: 352) (**Figure S17B**). A substantial proportion of these overlapping proteins showed shared up-or downregulation between *rup-1* and *rup-2* (MG-79**↑**, 78**↓**; BR-94**↑**, 128**↓**; RR-175**↑**, 154**↓**) (**Figure S17C)**. A six-set Venn analysis comparing unique and overlapping proteins across all ripening stages revealed that several proteins were common to two or more ripening stages (**Figure S18, Dataset S5**). The common proteins in all three ripening stages in *rup-1* and *rup-2* (14**↑**, 7**↓**) were largely related to protein homeostasis. GO enrichment analysis revealed that the differentially regulated proteins affected a broad spectrum of cellular functions (**Figure S19**). The most impacted categories included protein biosynthesis and protein homeostasis, particularly the ubiquitin-proteasome system. Additional enriched categories included carbohydrate metabolism and cellular respiration. Markedly, proteins related to redox homeostasis, amino acid biosynthesis, and enzyme classification exhibited greater upregulation than downregulation.

The mapping of differentially regulated proteins onto the overall metabolic network revealed that *rup-1* and *rup-2* affected several metabolic pathways in the ripening fruits relative to AV (**Figure 5**). The impact of the *rup* variants on metabolism was dynamic and varied with the progression of ripening. Consistent with lower amino acid levels, most enzymes involved in amino acid biosynthesis were downregulated, while those associated with amino acid degradation were upregulated in *rup-1* and *rup-2* (**Figure S20)**. Likewise, carbohydrate metabolism was broadly affected, which may have led to reduced sugar levels observed in fruits of *rup-1* and *rup-2* (**Figure S21-22)**. Similarly, the elevated C1 metabolism in *rup-1* and *rup-2* appears to be associated with increased folate levels in RR fruits.

**Figure 5.**
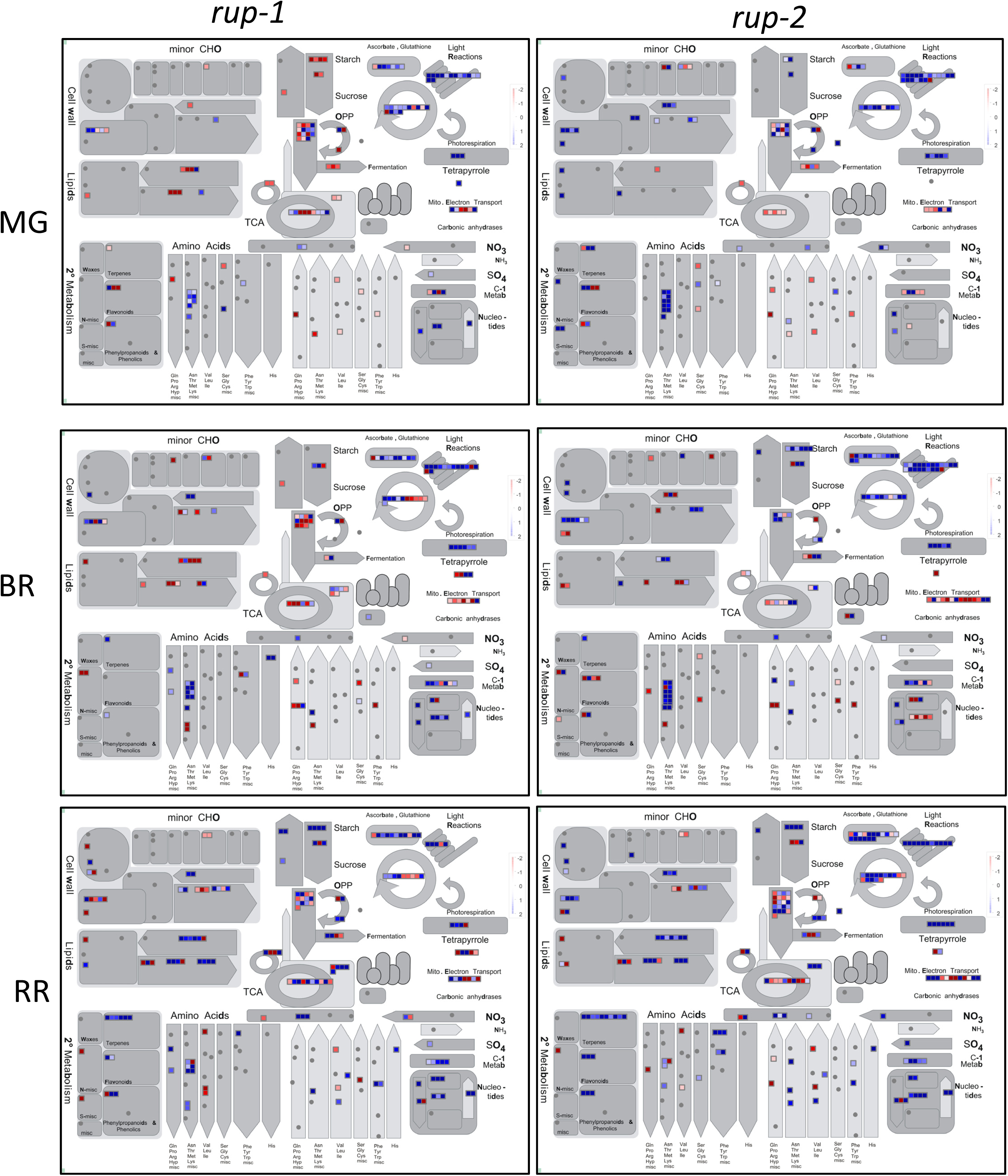
Mapman depiction of proteome changes in *rup-*variants fruits at different stages of ripening. Each square represents one protein. The red square indicates the upregulation of proteins, and the blue square indicates the downregulation of proteins with respect to AV. Only significantly different /proteins (log2 fold ≥±0.58, P ≤0.05) are depicted on the map. Data are mean ± SE (n = 3), P ≤ 0.05. (For details, see **Dataset S4**). MG, Mature green; BR, Breaker; RR, Red ripe.

A detailed examination of C1 metabolism revealed that two key enzymes involved in folate biosynthesis, formyl tetrahydrofolate ligase and methylene tetrahydrofolate reductase, were upregulated at the RR stage in *rup-1* and *rup-*2 **(Figure 6).** The higher folate levels were also supported by the increased abundance of serine hydroxymethyl transferase, which facilitates the transfer of a hydroxymethyl group from serine to tetrahydrofolate. Likewise, elevated levels of S-adenosyl-L-homocysteine hydrolase and methyl-tetrahydrofolate-dependent methionine synthase contribute to maintaining the methylation cycle by balancing the levels of S-adenosyl-methionine and S-adenosyl-L-homocysteine.

**Figure 6.**
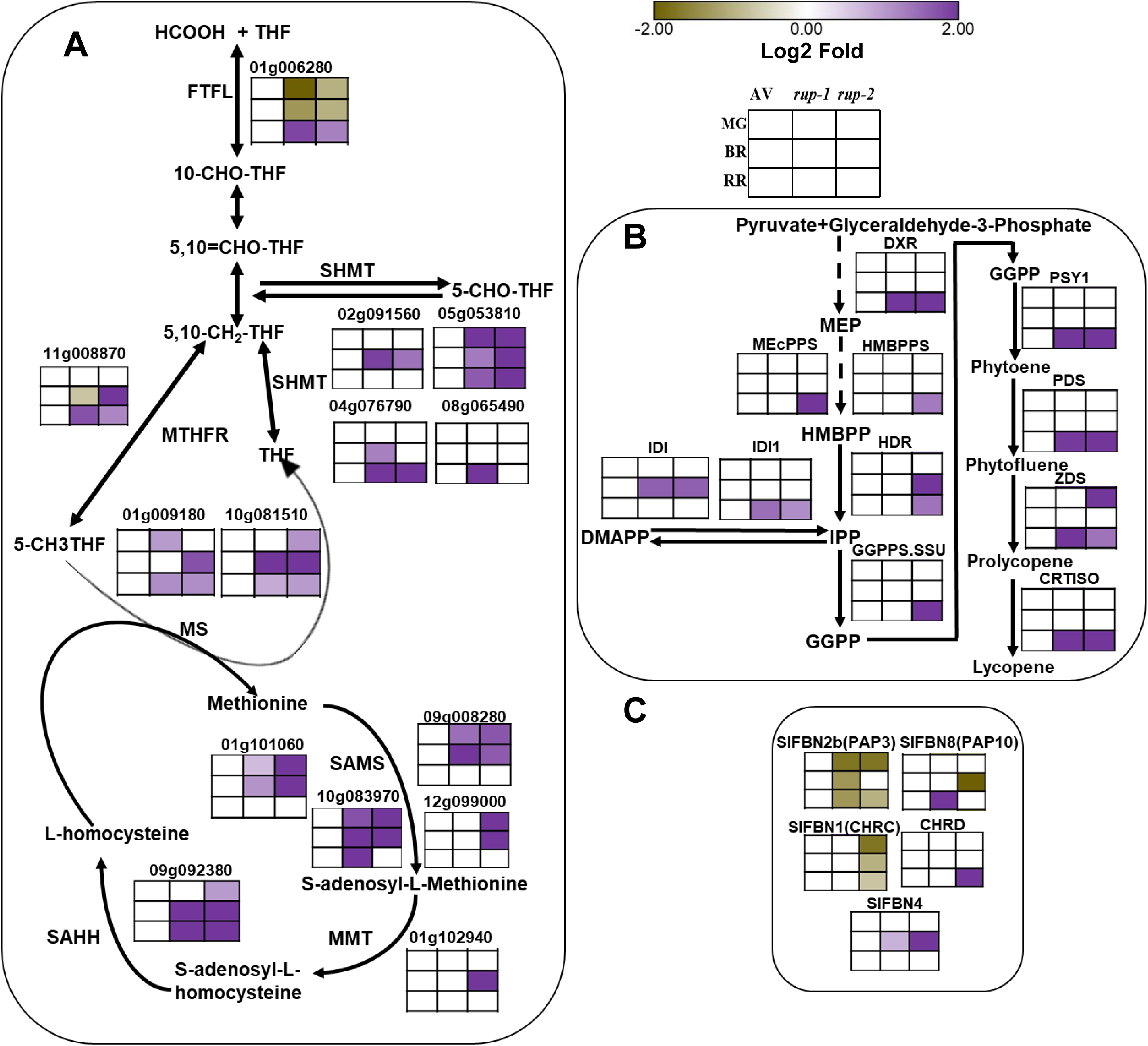
Regulation of C-1 metabolism and carotenoids biosynthesis in *rup-1* and *rup-2* relative to AV. **A.** Changes in the abundance of proteins of C-1 metabolism. **B.** Changes in the level of proteins regulating methylerythritol and carotenoid biosynthesis pathway. **C.** Changes in levels of carotenoid storage and sequestration-related proteins. The grid on top of the figure depicts the ripening stages used for AV, *rup-1,* and *rup-2* fruits. The SOLYC prefix before each protein ID is removed for convenience. Only significantly different proteins (Log2 fold > ± 0.584, p < 0.05, n = 3) relative to AV are shown on heatmaps. See **Dataset S4** and **S6** for detailed proteome data. **Abbreviations:** THF-Tetrahydrofolate; 10-CHO-THF-10-formyl-THF; 5,10=CH-THF-5,10 methenyl-THF; 5,10-CH2-THF-5,10 Methylene-THF; 5-CHO-THF-5-Formyl-THF; CHRD-Constitutive plastid-lipid-associated protein; CRTISO-Carotenoid isomerase; DMAPP-Dimethylallyl diphosphate; DXR-1-Deoxy-D-xylulose-5-phosphate reductoisomerase; FTFL-Formyl tetrahydrofolate ligase. DXS-1-deoxy-D-xylulose-5-Phosphate synthase; GGPPS.SSU-Geranylgeranyl diphosphate synthase small subunit subfamily type II; HCOOH-Formate; HMBPPS, 4-Hydroxy-3-methylbut-2-en-1-yl diphosphate synthase; IDI-Isopentenyl diphosphate Δ-isomerase; IPP-Isopentenyl pyrophosphate; MEcPPS-2-C-methyl-D-erythritol 2,4-cyclodiphosphate synthase; MMT-Methionine S-methyltransferase; MS-Methionine synthase; MTHFR-Methylenetetrahydrofolate reductase; PDS-Phytoene desaturase; PSY1-Phytoene synthase; SAHH-S-adenosyl-L-homocysteine hydrolase; SAM-S-adenosylmethionine; SAMS-S-Adenosylmethionine synthetase; SHMT-Serine hydroxymethyl transferase; SlFBN-Tomato fibrillin; THF-Tetrahydrofolate; ZDS-ζ-carotene desaturase.

The increased levels of lycopene appear to be associated with the upregulation of the methylerythritol phosphate (MEP) pathway and lycopene biosynthesis pathways, particularly at the RR stage. Two key enzymes of the MEP pathway, 1-deoxy-D-xylulose-5-phosphate reductoisomerase, and isopentenyl-diphosphate delta-isomerase1, were upregulated in *rup-1* and *rup-2*. In addition, several other enzymes in the pathway were upregulated in RR fruits in *rup-2.* Post-GGPP formation, four enzymes involved in lycopene synthesis—phytoene synthase 1, phytoene desaturase, ζ-carotene desaturase, and carotene isomerase—were upregulated in *rup-1* and *rup*-2. Elevated levels of fibrillins, (FBN8, FBN4) and the constitutive plastid-lipid-associated protein (CHRD) may further contribute to increased lycopene accumulation by facilitating its sequestration in chromoplasts.

Compared to AV, the cellular metabolic homeostasis of *rup-1* and *rup-2* showed broad alterations. The primary affected pathways included cell wall modification, glycolysis, the TCA cycle, and carbohydrate metabolism (**Figure S21–S24**). A notable feature was the enhanced protection of proteins, mediated by the modulation of chaperonins and the ubiquitin-proteasome system (**Figure S25–S27**). This increased protein protection may represent a compensatory response to the reduced UV-B protection observed in the *rup*-variants due to the loss of RUP function. Interestingly, despite significant changes in overall metabolism, the hormonal regulatory pathways were minimally affected, with variations limited to a few proteins primarily involved in jasmonic acid biosynthesis (**Figure S28**).

## Discussion

Tomato fruit ripening begins with a climacteric surge in ethylene, which activates metabolic pathways responsible for fruit softening and the conversion of chloroplasts into lycopene-rich chromoplasts. Although light is not obligatorily required for ripening, light plays a modulatory role. Both ethylene and light signalling pathways modulating ripening converge on the regulation of proteolysis for signal transduction (**Yang et al., 2010; Deng et al., 2018; Wang et al., 2019**). Photoreceptors, including phytochromes, cryptochromes, and UVR8, associate with the ubiquitin-proteasome system (UPS) to mediate photomorphogenic responses (**Eckardt et al., 2024**). Given that tomato UVR8 can complement UV-B signalling in Arabidopsis *uvr8* mutants and interacts with COP1 (**Dong et al., 2021**), UV-B signalling in tomatoes likely involves UPS modulation, similar to the pathway in Arabidopsis.

### *rup*-variants exhibit higher accumulation of carotenoids relative to AV

Tomato RUP exhibits strong homology to AtRUP1 and AtRUP2 (**Zhang et al., 2021**). Similar to the *Atrup1rup2* mutant (**Gruber et al., 2010**), *Slrup* mutants display reduced hypocotyl elongation under UV-B compared to the wild type (WT) (**Zhang et al., 2021**). Since the *rup*-variants used by us were natural accessions, they had no wild-type control. Therefore, we compared them with the Arka Vikas (**AV**) cultivar, which carries the native *RUP* gene. Under UV-B, the *rup*-variants showed a phenotype more similar to acclimated *Slrup* mutants (**Zhang et al., 2021**), as the reduction of hypocotyl elongation was milder than in AV. This phenotypic variation may stem from differences in their mutated gene sequences: *Slrup^CR8^* encodes a truncated protein lacking 37 amino acids (E20 to A56), while *Slrup^CR22^* produces a jumbled peptide after R18* (**Zhang et al., 2021**). These structural differences could affect RUP-UVR8 interactions, particularly since residues D122, E138, and D140 in Arabidopsis RUP2 are critical for UVR8 re-dimerization *in planta* and *in vitro* (**Wang et al., 2023**). Although the *rup*-variants encode a truncated 164AA protein, they retain conserved residues at D121, E137, and D139 (**Table S2**), suggesting potential UVR8 binding capacity.

Adult *rup*-variant plants were phenotypically similar to Arka Vikas (AV), except for *rup-1*, which exhibited slightly higher leaf chlorophyll levels. A key difference between AV and the *rup*-variants was the significantly higher carotenoid content in *rup*-variant fruits, even higher than that of *RUP*-RNAi-silenced lines (**Liu et al., 2004**). Considering RUP terminates UV-B signalling by facilitating UVR8 re-dimerization, it is plausible that impaired UVR8 re-dimerization in *rup*-variants leads to increased UVR8 monomer accumulation, which promotes carotenoid biosynthesis. Consistent with the above notion, *UVR8* overexpression enhances carotenoid levels (1.25-fold) in tomato fruits (**Li et al., 2018**). Since total carotenoid levels in *rup-1* (1.67-fold) and *rup-2* (1.52-fold) were much higher than the *UVR8^OE^* line, it is likely that the loss of *rup* activity and increased UVR8 monomer levels combinedly contribute to increased carotenoid levels.

Since we compared the carotenoid levels in the *rup*-variant with another tomato cultivar AV, it remains possible that higher carotenoid levels were intrinsic to *rup*-variants but not due to the truncation of RUP protein. The introgression of *rup*-variants in Arka Vikas’s background negates this possibility. The fruits bearing homozygous *rup*-variants alleles in the BC_4_F_2_ generation had higher carotenoid levels like *rup-1* and *rup-2.* Since the expected recovery by BC_4_F_2_ generation is ≥ 99% of the recurrent parental genome (**Hospital, 2003**), the higher carotenoid levels in BC_4_F_2_ fruits can be attributed to the introgression of *rup*-variant genes in Arka Vikas background.

### Metabolic pathways are attenuated in BR *rup*-variant fruits

Although the *rup*-variant fruits were smaller than AV, only a few metabolites differed at the MG stage. However, at the BR stage, *rup*-variants unusually exhibited lower levels of key metabolites—particularly sugars and metabolites derived from sugars. In contrast, another light-signalling mutant-*Nps1*, shows higher levels of sugars in fruits (**Kilambi et al., 2021**). Though *rup*-variants displayed an overall downregulation of the primary metabolome, the fruits had higher levels of carotenoids and folate, similar to a *trifoliate* mutant (**Tyagi et al., 2022**). While it is not possible to distinguish whether the lower metabolite levels result from increased consumption or reduced biosynthesis, their lowering indicates that the cellular homeostasis of *rup*-variant fruits at the BR stage differs from AV. Notwithstanding the diverse genetic backgrounds, the metabolic lowering of sugar was common between *rup*-*1* and *rup*-*2*. This entails that *rup*-variants influence the metabolome in a similar fashion as their encoded protein sequences are nearly identical.

In both *rup*-variants, at all ripening stages, the reduced levels of key metabolism regulatory proteins, such as 6-phosphogluconolactonase (key enzyme for oxidative pentose phosphate pathway), aconitase (TCA cycle) and acid β-fructofuranosidase (sucrose hydrolysis) signifying lowering of the metabolic processes. The concurrent reduction in sugars and upregulation of carotenoids/folate suggests a shift in resource allocation, diverting carbon from primary metabolism to secondary metabolites (**Zhu et al., 2022**). Taken together, it entails that *rup-1* and *rup-2* influence cellular homeostasis using a common signalling pathway.

The prominent effect of RUP on metabolite profiling at the BR stage also correlates with its increased expression after the onset of fruit ripening. The metabolite profiles of RR fruits of BC_1_F_2_ homozygous *rup-1* and *rup-2* plants are nearly similar and differ from AV and respective *rup*-parental lines. The above similarity in metabolic profiles supports that the lowering of metabolite levels emanates from the influence of truncated *rup-1* and *rup-2* on the cellular metabolome.

The wide-ranging influence of *rup*-*1* and *rup*-*2* on cellular metabolome—including elevated folate and carotenoid levels—may stem from the persistence of monomeric UVR8 form, leading to exaggerated signal transduction. Considering that Arabidopsis RUP is also a constituent of CUL4-DDB1-based E3 ubiquitin ligase (**Ren et al., 2019**), it is plausible that tomato RUP similarly regulates the ubiquitin-proteasome system. The above view is supported by the conservation of UV-B signalling between tomato and Arabidopsis, as tomato UVR8 can functionally complement UV-B responses in the *Atuvr8* mutant (**Dong et al., 2021**). Considering that mutation in DDB1 (*hp1*)—another key constituent of CUL4-DDB1-based E3 ubiquitin ligase—also increases carotenoid levels in tomato fruits (**Wang et al., 2019),** the truncation of RUP may similarly affect the E3 ubiquitin ligase activity.

### Carotenoid biosynthesis proteins are upregulated in *rup*-variants

Proteomic analysis of *rup*-variant fruits revealed significant differences in protein abundance compared to AV across multiple ripening stages. Mainly, protein homeostasis and protein biosynthesis were the largest altered functional classes at all ripening stages, hinting at the modulation of the ubiquitin-proteasome system. Consistent with observed reductions in sugar levels, proteins involved in carbohydrate metabolism and cellular respiration were also prominently affected. The striking similarity in protein modulation patterns between *rup-1* and *rup-2* during fruit ripening further supports the view that both variants influence cellular homeostasis through a shared mechanism.

A key feature of *rup*-variants was elevated levels of nutraceuticals, mainly carotenoids and folate, in red-ripe fruits. The increased levels of the above nutraceuticals seem to be related to the enhanced biosynthesis of these compounds. Consistent with this notion, most proteins involved in folate vitamer biosynthesis and transformation were upregulated in RR *rup*-variant fruits (**de la Garza et al., 2007; Tyagi et al., 2022, 2023)**. Notably, the elevated levels of enzymes such as formyl tetrahydrofolate ligase, methylene tetrahydrofolate reductase, serine hydroxymethyl transferase, and methionine synthase likely contribute to higher folate content. Parallel to the above, the proteins associated with methionine recycling also show elevated levels at one or more stages of ripening. Similar to *rup*-variants, higher folate levels in the *trifoliate* tomato mutant correlate with increased levels of proteins related to folate transformation and C1 metabolism (**Tyagi et al., 2022, 2023**).

In tomato fruits, carotenoid accumulation predominantly occurs during the transition from BR to RR stages. Unlike gibberellins, which are synthesized via the mevalonate pathway, the carotenoid precursor geranylgeranyl diphosphate (GGPP) is derived from the methylerythritol phosphate (MEP) pathway (**Gupta and Hirschberg, 2022**). Several MEP pathway proteins in *rup*-variants displayed higher levels at one or more ripening stages. Specifically, 1-deoxy-D-xylulose-5-phosphate reductoisomerase (DXR) and isopentenyl diphosphate Δ-isomerase (IDI) showed higher abundance at the RR stage in both *rup*-variants, suggesting an enhanced supply of GGPP for carotenoid biosynthesis.

Most importantly, the levels of key carotenoid biosynthesis proteins-phytoene synthase 1 (PSY1), phytoene desaturase (PDS), ζ-carotene desaturase (ZDS), and carotenoid isomerase (CRTISO) in red-ripe fruits are high in both *rup*-variants (**Isaacson et al., 2002; Fray et al., 1995; McQuinn et al. 2023)**. The upregulation of these proteins correlates with increased levels of intermediary molecules like phytoene and phytofluene, leading to enhanced carotenoid accumulation in *rup*-variants. Similarly, in the *MYB117* mutant, elevated levels of ζ-carotene isomerase and CRTISO contribute to higher carotenoid content (**Tyagi et al., 2022**). Given that transgenic manipulation of PSY, PDS, ZDS, and CRTISO modulates carotenoid levels in tomato fruits (**Fantini et al., 2013; Fraser et al., 2002**; **McQuinn et al., 2018**), it is plausible that increased carotenoid accumulation in *rup*-variants results from the higher abundance of these four key enzymes.

### Protein protection enhances carotenoid biosynthesis in *rup*-variants

Given that *rup*-variants show elevated carotenoid levels, similar to other light-signalling mutants like *hp1* (DDB1) and *hp2* (DET1) (**Liu et al., 2004; Wang et al., 2008**), it is plausible that RUP proteins functionally cooperate with these proteins. The above assumption is consistent with recent reports that RUP proteins physically interact with the CUL4-DDB1 E3 ubiquitin ligase complex to modulate UPS-mediated protein degradation (**Huang et al., 2014; Podolec et al., 2021**). In Arabidopsis, RUP proteins interact with DDB1 to regulate the turnover of HY5, a key transcription factor regulating light signalling (**Ren et al, 2019**). In like manner, tomato RUP may modulate the protein degradation activity of UPS by associating with DDB1. Arabidopsis RUPs also directly interact with both COP1 and HY5 (**Ren et al., 2019**), and disruption of these interactions in *rup*-variants may additionally influence the cellular metabolome.

In *rup*-variants, the truncation of RUP proteins may impair DDB1 function by disrupting RUP-DDB1 interactions, potentially leading to HY5 stabilization (**Ren et al., 2019**). Consistent with this, tomato DDB1 (*hp1*) and (DET1) *hp2* mutants show increased levels of

HY5, which reportedly binds to the promoters of key carotenogenesis genes, facilitating elevated carotenoid levels **(Liu et al., 2015; Wang et al., 2021).** Furthermore, the substantial reduction of *CYCB* expression in *rup*-variant fruits is consistent with the observation that HY5 negatively regulates *CYCB* expression, as *HY5-*silenced tomato lines show higher *CYCB* expression (**Wang et al., 2021).**

Except for *CYCB*, expression levels of carotenogenesis and light signalling-related genes in *rup*-variants showed no significant differences from AV. Ostensibly, the enhanced carotenoid accumulation in *rup*-variants may occur through an alternate pathway. Given that silencing of *DDB1* (*hp1*) in tomato fruits affects ribosomal protein accumulation and impairs translational capacity (**Wang et al., 2021**), it brings forth the possibility that *DDB1* can regulate carotenogenesis through post-transcriptional control of protein levels. The observed increase in key carotenoid biosynthetic enzymes - PSY1, PDS, ZDS, and CRTISO – supports the possibility that modulation of protein protection mechanisms is likely responsible for their elevated levels. The elevated levels of folate metabolism-related proteins are also consistent with a protein protection mechanism operating in *rup*-variants.

### HSP70-2 suppression elevates carotenoid levels in *rup*-variants

The protein protection mechanism functions through multiple pathways, primarily involving reduced protein proteolysis and chaperone-mediated protein stabilization (**Kim et al., 2013**). In *rup*-variants, elevated levels of ribosomal components, such as RPL3 (critical for ribosome assembly, translation, and peptidyl transferase activity), RPP0 (essential for recruitment of elongation factors), and the LSU of the ribosome (containing the peptidyl transferase centre) across all ripening stages suggest sustained protein protection (**Database S5**). Given that protein homeostasis and protein biosynthesis were the largest altered functional classes in *rup*-variants, the better protection of proteins may effectively mitigate any potential deleterious effects caused by the persistent presence of UVR8.

Proteomic analysis revealed significant downregulation of multiple ubiquitin-proteasome system (UPS) components in *rup*-variants compared to AV, including E1 ubiquitin-activating enzymes, E2 ubiquitin-conjugating enzymes, RING-type E3 ligases, deubiquitinating enzymes (DUBs), and 19S regulatory particles. As these proteins are essential for UPS-mediated protein degradation, their reduced abundance likely attenuates proteasomal processing, resulting in protein accumulation (**Eckardt et al., 2024**). The increased levels of folate and carotenoid biosynthesis pathway proteins are consistent with reduced protein degradation in *rup*-variants, leading to higher levels of respective nutraceuticals.

In tomato, the transcript levels of *Hsp70*.*2* (Solyc01g106210) increase during fruit ripening. Silencing of *Hsp70*.2 revealed that it negatively regulates carotenoid biosynthesis without altering chromoplast ultrastructure (**D’Andrea and Rodriguez-Concepcion, 2019**). Since in RR fruits of *rup*-variants, the level of Hsp70.2 protein is strongly downregulated, being a negative regulator, it may assist carotenoid accumulation by increasing the PSY1 protein level. Though the ClpC1 (**Solyc12g042060)** level, which targets PSY for degradation, is upregulated in *rup*-variants, the reduced Hsp70.2 levels would mitigate the interaction between PSY1 and ClpC1. Consistent with this, in Arabidopsis, a mutation in the *Hsp70.2* gene leads to the upregulation of the PSY protein in chloroplasts (**Welsch et al., 2018**).

In tomato, transgenic constitutive overexpression of *Hsp21* promotes lycopene accumulation during fruit ripening (**Neta-Sharir et al., 2005**). While the higher accumulation of HSP21 stimulates carotenoid levels in tomato *Nps1* mutant (**Kilambi et al., 2021**), the carotenoid accumulation in *rup*-variants is not linked with Hsp21, as it is downregulated in ripening fruits. Moreover, unlike *rup*-variants, *Nps1* fruits exhibit higher levels of Hsp70.2 protein. Although both *Nps1* and *rup*-variants modulate light signalling, they regulate carotenogenesis through different mechanisms. The *rup*-variants are more similar to the *Hsp70*.*2*-silenced tomato lines, showing no significant changes in the transcript levels of carotenogenesis-related genes. It is proposed that Hsp70.2 targets carotenoid pathway enzymes for degradation via Clp proteases, and its reduced levels allow other chaperones to reactivate inactive enzymes (**D’Andrea and Rodriguez-Concepcion, 2019**). Furthermore, reduced levels of E1, E2, and RING-E3 ligases in *rup*-variants may impair UPS-mediated degradation of carotenogenic enzymes. The elevated carotenoid levels in *rup*-variant fruits, along with higher levels of carotenogenesis pathway proteins and lower levels of Hsp70.2, broadly conform to the above proposal.

Molecular-genetic analyses of tomato mutants, such as *hp1* (DDB1) and *hp2* (DET1), have highlighted the critical role of CRL4C3D E3 ubiquitin ligase complexes (comprising COP10-DDB1-DET1-DDA1) in regulating carotenogenesis. Emerging evidence indicates that *hp1* may enhance carotenogenesis at dual levels, protecting HY5, a transcriptional activator, and modulating protein translation machinery (**Wang et al., 2021**). Similarly, *rup*-variants likely stimulate carotenogenesis in fruits by protecting HY5 and carotenogenesis enzymes. The proteomic and metabolomic data support the view that alteration of protein homeostasis in *rup*- variants is causally related to increased levels of carotenoid biosynthesis enzymes. The role of the Hsp70.2 chaperone in increasing carotenoid levels needs to be further investigated. To sum up, our study provides RUP as a novel regulator for modulating carotenoids and folate levels in tomato.

## EXPERIMENTAL PROCEDURES

### Screening for SNPs in *RUP* gene

The population used for the *RUP* screening was the same as described earlier in **Mohan et al. (2016)**, **Sarma et al. (2017),** and **Upadhyaya et al. (2016) (Dataset S7).** Tomato cultivar Arka Vikas (https://iihr.res.in/variety/arka-vikas) was used as a control for population screening and *rup*-variant comparisons. Genomic DNA was isolated from tomato accessions following the procedure of **Sreelakshmi et al. (2010).** *RUP* (Solyc11g005190) sequence was obtained from the Solanaceae Genomics Network (https://solgenomics.net/). The SNP detection was carried out as described by **Mohan et al. (2016),** followed by Sanger sequencing (**Table S3**). The CEL-I assay was used to ascertain the homozygosity of *rup*-variants in backcrossed progeny. SIFT (Sorting intolerant from tolerant; **Sim et al., 2012**) was used to predict the deleterious variations for protein functions. The modeling of the RUP protein of AV, *rup-3*, and *rup-4* was carried out using DDmut software **(Zhou et al., 2023**; https://biosig.lab.uq.edu.au/ddmut).

For phenotype analysis, seedlings were grown in plastic boxes on an agar medium (0.8% w/v agar) in total darkness or white light (100 µmol m^-2^s^-1^). The light treatment consisted of 16 h light and 8 h dark periods. For UV-B supplementation, plants were exposed to 4 hours of UV-B light (4 µmol m^-2^s^-1^) daily at the same time at mid-noon.

### Biochemical characterizations

The ripening stages of fruits were monitored as described by **Gupta et al. (2014).** Plants were grown in the greenhouse under the natural photoperiod (12-14 h day, 28±1°C; 10-12 h night, 14-18°C). The °Brix, fruit diameter, and firmness were determined as described by **Gupta et al. (2014)**. Spectrophotometric determination of carotenoids was carried out as described by **Mohan et al. (2016).** The total chlorophyll level in leaves was determined using the method of **Porra et al. (1989**). The carotenoid profiling was carried out following the procedure of **Gupta et al. (2015)**. Folate extraction and analysis were performed as described by Tyagi et al. (2015)

### Metabolic profiling

The primary metabolite analysis from seedlings and fruits using Leco Pegasus GC-MS was carried out by a protocol described in **Bodanapu et al. (2016).** Only metabolites with a ≥1.5-fold change (log2 ± 0.584) and p-value ≤0.05 were considered to be significantly different. PLS-DA was performed using Metaboanalyst 5.0 (http://www.metaboanalyst.ca/). Heat maps were generated using Morpheus (https://software.broadinstitute.org/morpheus/).

For metabolites detected only in *rup*-variants or AV, the missing values were replaced with 1/10 of the lowest value in the dataset.

### Gene Expression Analysis

The pericarp from fruit tissue was frozen in liquid N_2_ and homogenized to a fine powder. About 100 mg of homogenized powder was used for RNA extraction with the RNeasy kit (Qiagen). The extraction was carried out following the manufacturer’s instructions. Total RNA was reverse transcribed into cDNA using the first strand cDNA synthesis kit (Invitrogen, India) following the manufacturer’s instructions. The real-time quantitative PCR assays were performed as described in **Kilambi et al. (2013).** The primer sequences are given in **Table S3. Proteome profiling** Proteins (70 µg) were separated, identified, and quantified as described in **(Kilambi et al., 2016; 2021).** *Solanum lycopersicum* iTAG4.1 proteome sequence (ftp://ftp.solgenomics.net/tomato_genome/annotation/ITAG4.1_release/ITAG4.1_proteins.fas <u>ta</u>) was used as the database against which the searches were done. The functional annotation and classification of the ITAG4.1 proteins was carried out using the Mercator 4 tool (https://www.plabipd.de/portal/mercator4). The proteins with ≥ 1.5-fold change (Log2 ± 0.58) and p-value ≤ 0.05 were considered as differentially expressed. For proteins detected only in *rup-*variants or AV, the missing values were replaced with 1/10 of the lowest value in the dataset. The significantly expressed proteins were mapped on the overall metabolic pathway using Mapman 3.6 ORC1 software (https://mapman.gabipd.org/home).

### Statistical analysis

A minimum of three independent biological replicates was used for all experiments. Statistical Analysis On Microsoft Excel (https://prime.psc.riken.jp/MetabolomicsSoftware/StatisticalAnalysisOn-MicrosoftExcel) was used to obtain significant differences between data points. A Student’s *t*-test was also performed to determine significant differences (* for P ≤0.05). Heat maps and PLS-DA plots were generated using Morpheus (https://software.broadinstitute.org/morpheus/) and MetaboAnalyst 4.0 (https://www.metaboanalyst.ca/), respectively.

## ACKNOWLEDGMENTS

This work was supported by the Department of Biotechnology (DBT), India grants, BT/PR11671/PBD/16/828/2008, BT/PR/7002/PBD/16/1009/2012, and BT/COE/34/SP15209/2015 to RS and YS and BT/INF/22/SP44787/2021 to YS and RS. We also acknowledge the funds from DBT-BUILDER (BT/INF/22/SP41176/2020), MHRD-IOE (RC5-22-019) to the laboratory; CSIR fellowship to CC. We thank Dr. J. Giovannoni, Cornell University, Ithaca, USA, for providing seeds of *cop1-like* RNAi lines. Repository of Tomato Genomics Resources is a DBT-SAHAJ National facility.

## Conflict of Interest

The authors declare no conflict of interest.

## Author contributions—

CC did most experiments, SKG did backcrossing, carotenoids profiling of BC_1_F_2_ lines and metabolome analysis, AS did carotenoid profiling of BC_4_F_2_ lines and proteome analyses, JB did carotenoid and proteome profiling, SV did backcrossing, and UV-B phenotyping. KT did folate estimation, RK and HS did backcrossing, HVK measured fruit size, °Brix, and acidity, VS did the carotenoid estimation, and YS and RS designed the research and wrote the manuscript.

## Data Availability

The primary data of metabolite and proteome profiling is included as supplemental material. The mass spectrometry proteomics data are available via ProteomeXchange (**Vizcaíno et al., 2014**) with the dataset identifier PXD039219.

## Supplemental Data

**Figure S1.**
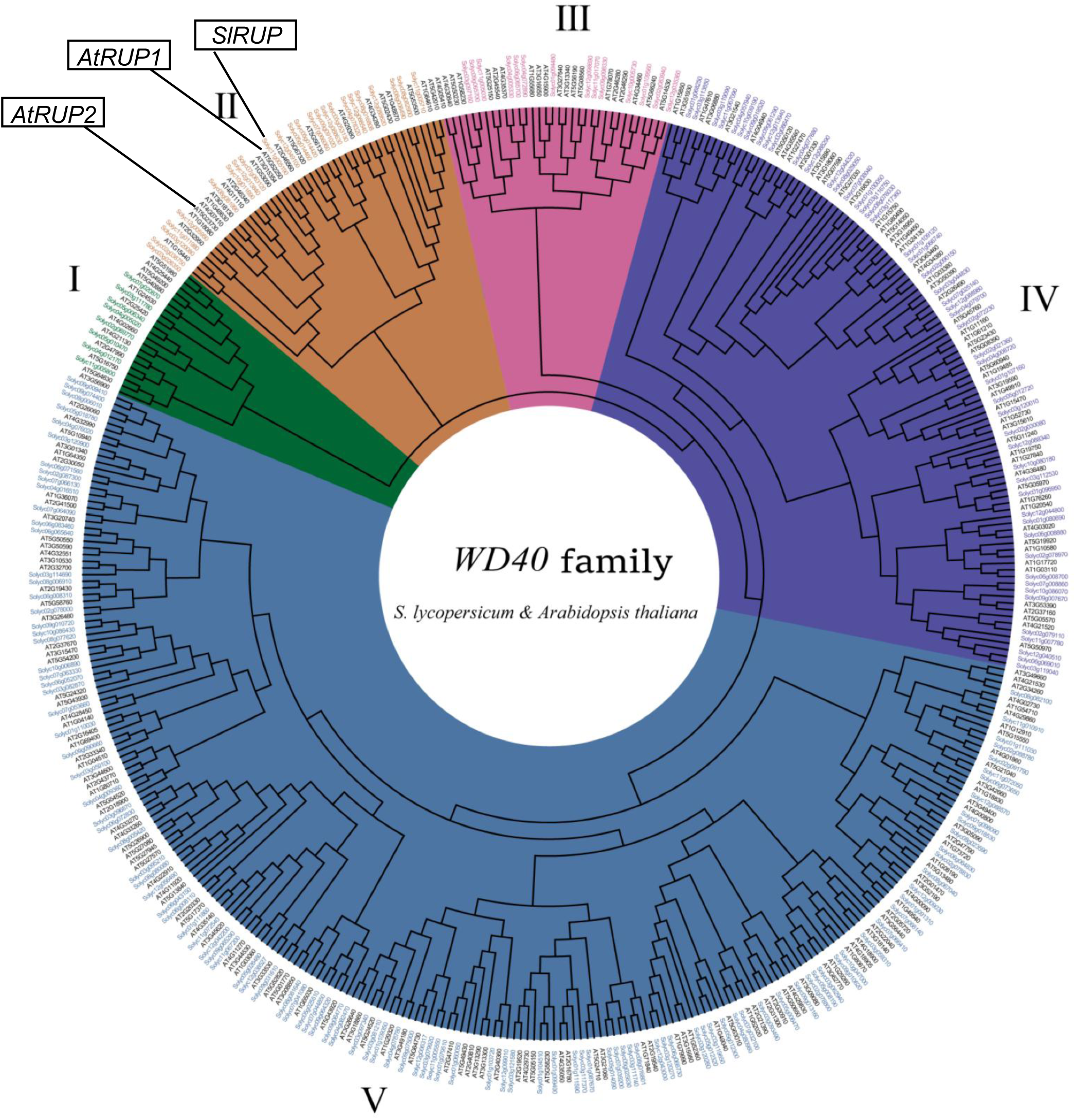
The dendrogram shows phylogenetic relationships between tomato and Arabidopsis WD40 proteins. The phylogenetic relation of tomato RUP (Solyc11g005190) protein with reference to Arabidopsis RUP1 and RUP2 proteins is depicted on the dendrogram. The dendrogram is from Figure 5 of **Yan C, Yang T, Wang B, Yang H, Wang J, Yu Q**. (2023) Genome-Wide Identification of the WD40 Gene Family in Tomato (Solanum lycopersicum L.). *Genes*.**14**: 1273. https://doi.org/10.3390/genes14061273

**Figure S2.**
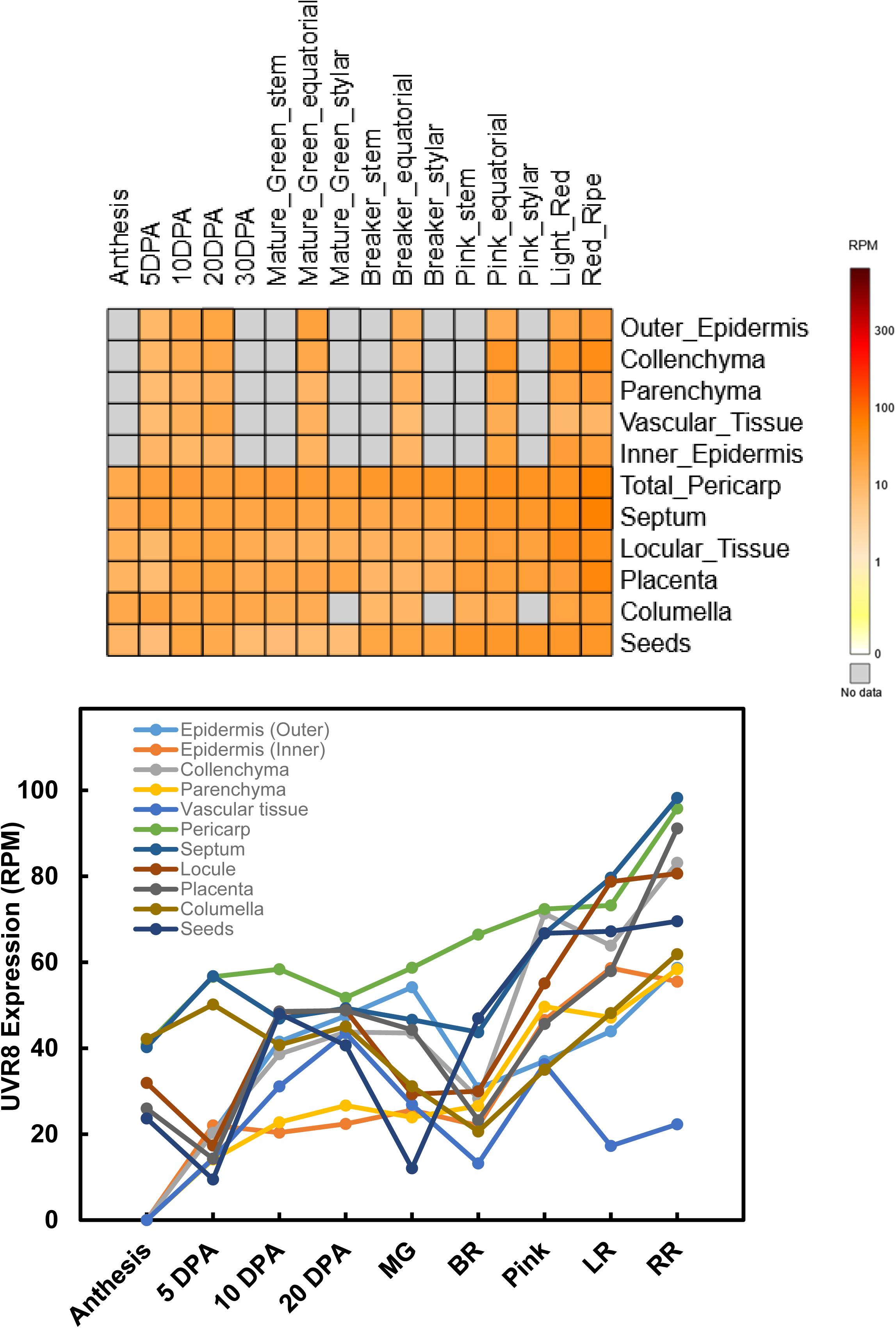
*UVR8* (Solyc05g018620) gene expression in tomato fruit. Data source-https://tea.solgenomics.net. Abbreviations: DPA-Days post-anthesis.

**Figure S3.**
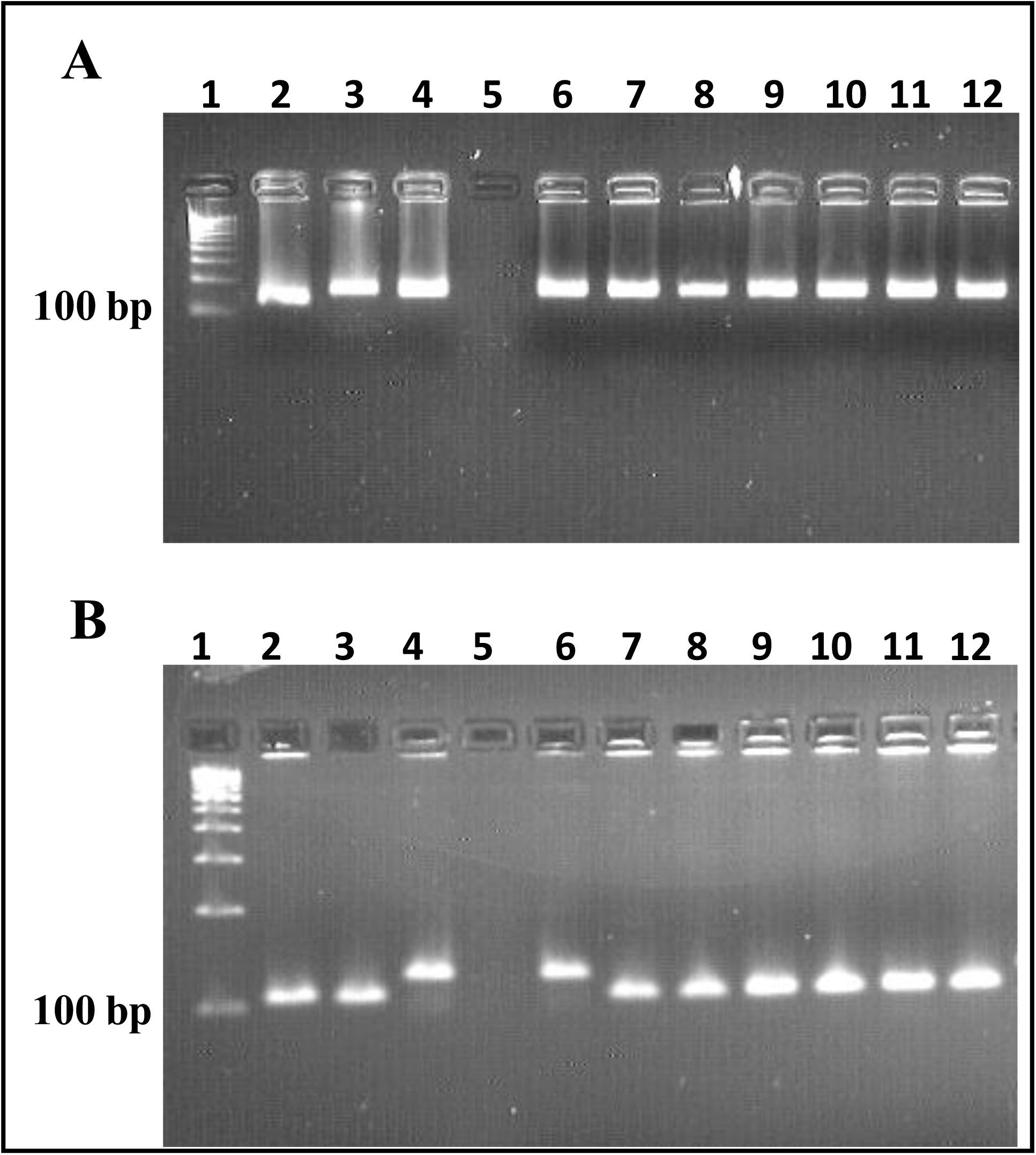
DNA profiling of Arka Vikas, Ailsa Craig, *S. pimpinellifolium, rup-1*, and *rup-2* using SSR markers. **A**-Marker SL100042i is polymorphic between Arka Vikas and Ailsa Craig but not with *S. pimpinellifolium*, *rup-1*, and *rup-2*. **B**-Marker U21085 is polymorphic between Arka Vikas and *S. pimpinellifolium* and not with others. Based on the above polymorphism, it was inferred that the *rup-1* and *rup-2* are closer to Arka Vikas. Lane 1-DNA Ladder, Lane 2-Ailsa Craig, Lane 3-Arka Vikas, Lane 4-*S*. *pimpinellifolium*, Lane 5-empty, Lane 6-*S*. *pimpinellifolium,* Lane 7,8,9-*rup-1*, and Lane 10,11&12-*rup-2.* See **Table S3** for the SSR primers sequence.

**Figure S4.**
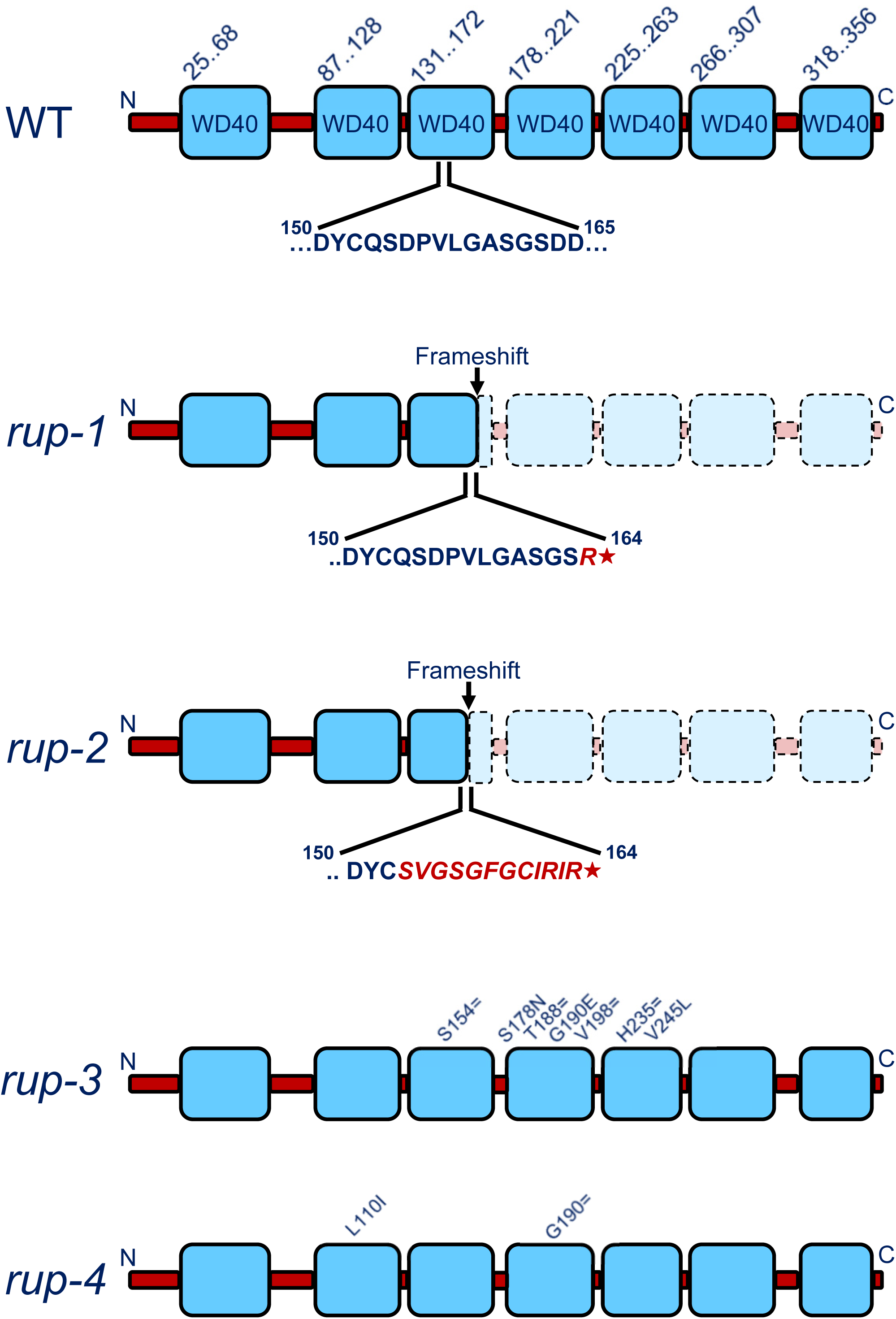
Functional domains of tomato RUP protein. The cartoon shows that in *rup-1* and *rup-2,* frameshift mutation leads to protein truncation at R164*. The mutations in *rup-3* and *rup-4* do not affect the formation of full-length protein. According to DDMut and SIFT analyses, the nonsynonymous SNPs in *rup-3* are tolerated and are unlikely to affect RUP function. For *rup-4*, results for nonsynonymous SNP were mixed: DDMut predicted a destabilizing effect, while SIFT indicated that the SNP is tolerated (**Table S1**).

**Figure S5.**
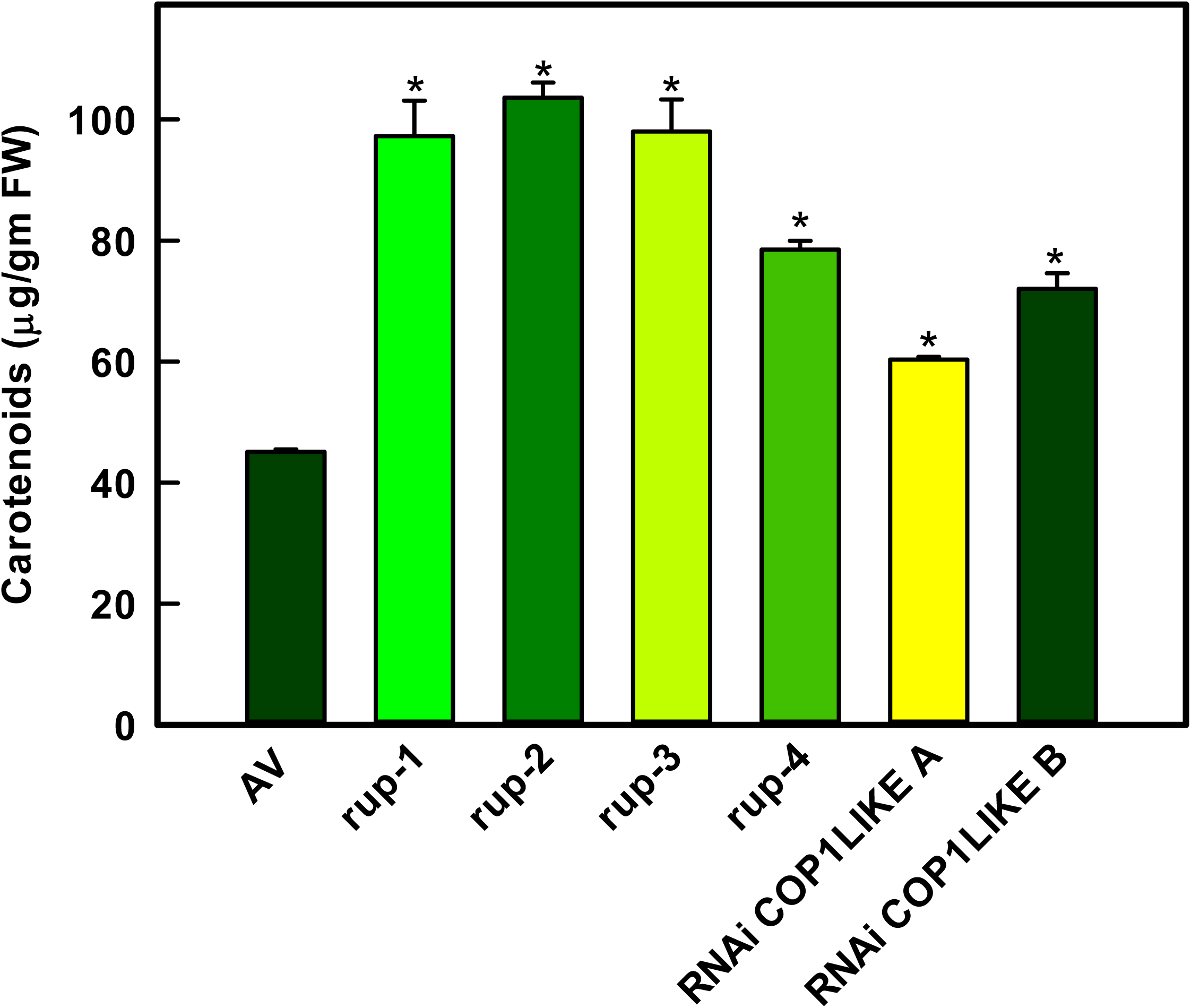
Carotenoid levels in red-ripe fruits of Arka Vikas (AV), *rup-*variants, and *RNAi-COP1-LIKE*. A spectrophotometric assay was used to determine the carotenoids. The carotenoid data are mean ± SE (n ≥3). Asterisks indicate significant differences with P < 0.05 (*) relative to AV.

**Figure S6.**
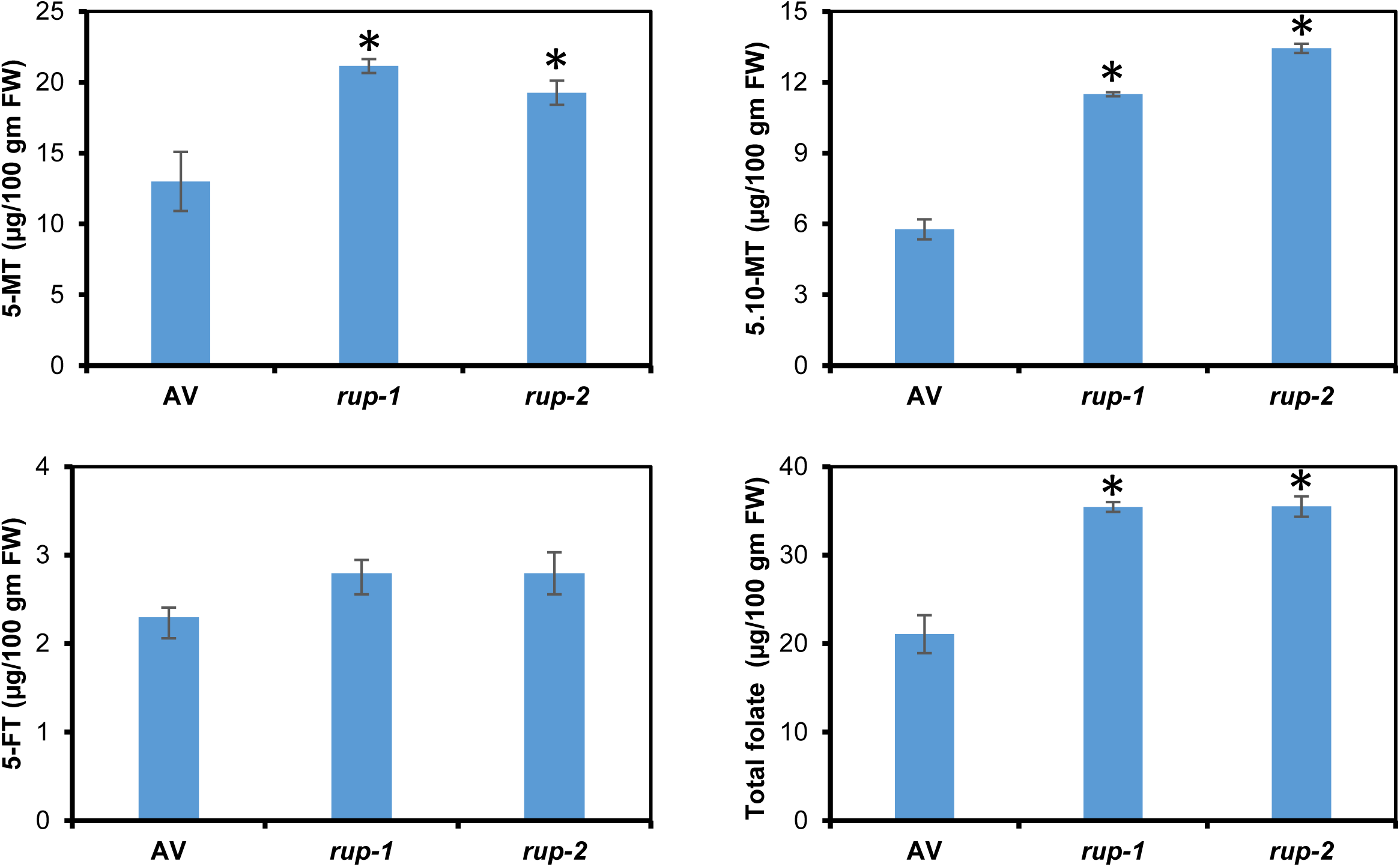
Total folate and its individual vitamers in red-ripe fruit of Arka Vikas (AV) and *rup*-variants. **Abbreviations:** 5,10-MT-5,10-methenyl tetrahydrofolate; 5-FT-5-Formyl-tetrahydrofolate; 5-MT-5-Methyl-tetrahydrofolate; T-Tetrahydrofolate. The folate data are mean ± SE (n ≥3). Asterisks indicate significant differences with P < 0.05 (*).

**Figure S7.**
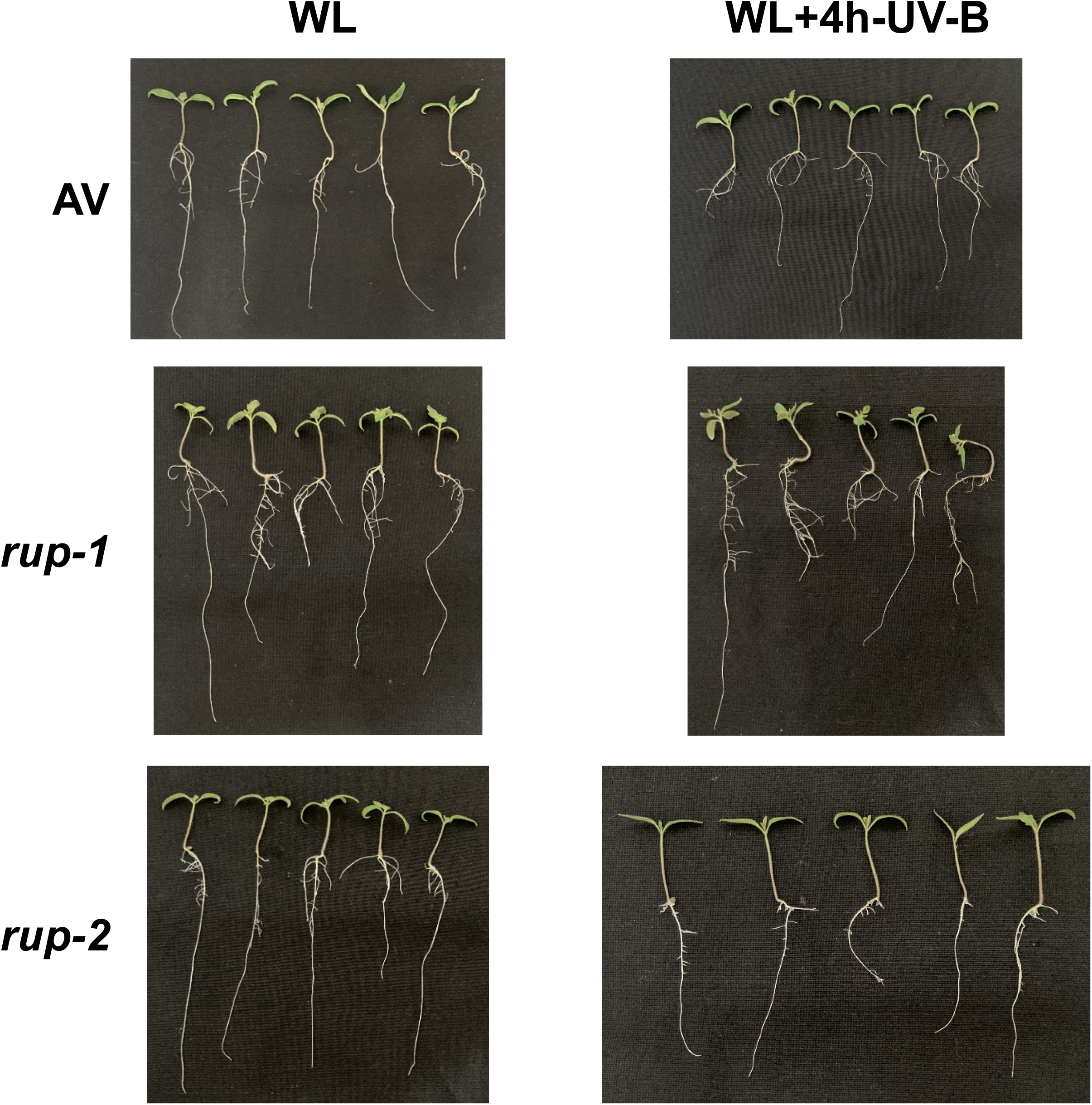
Seedling phenotype under white light (WL) or WL supplemented with 4 hours of UV-B exposure. Seeds germinated in darkness on agar and transferred to WL after radicle emergence. Seedlings were then exposed to 4 hours of UV-B light daily at the same time. Photographs were taken 7 days after radicle emergence.

**Figure S8.**
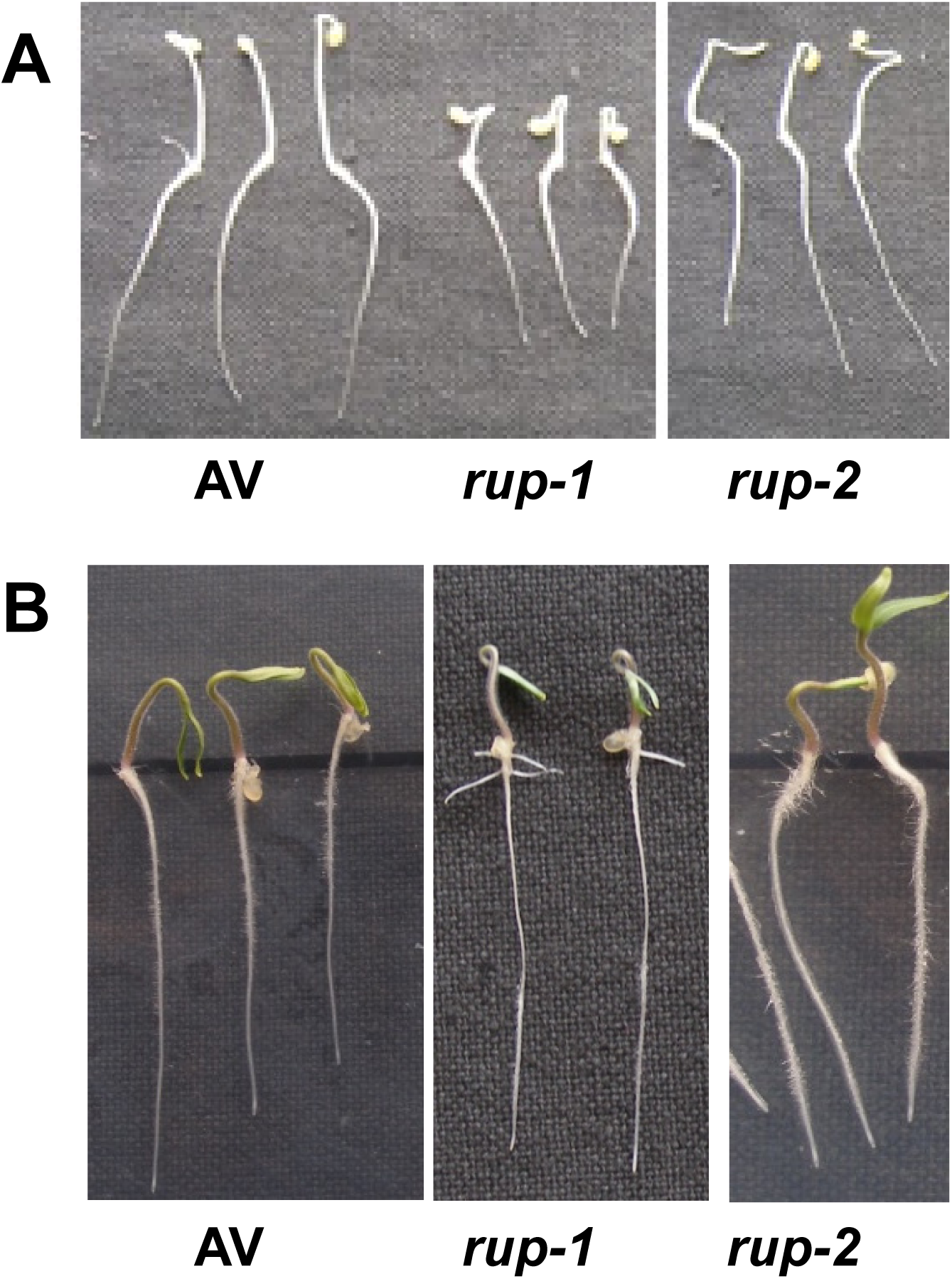
Phenotype of AV, *rup-1,* and *rup-2* seedlings grown under different growth conditions. **A.** Dark-grown seedlings (3-day-old) of AV and *rup*-variants. **B.** Light-grown seedlings (3-day-old) of AV and *rup-*variants.

**Figure S9:**
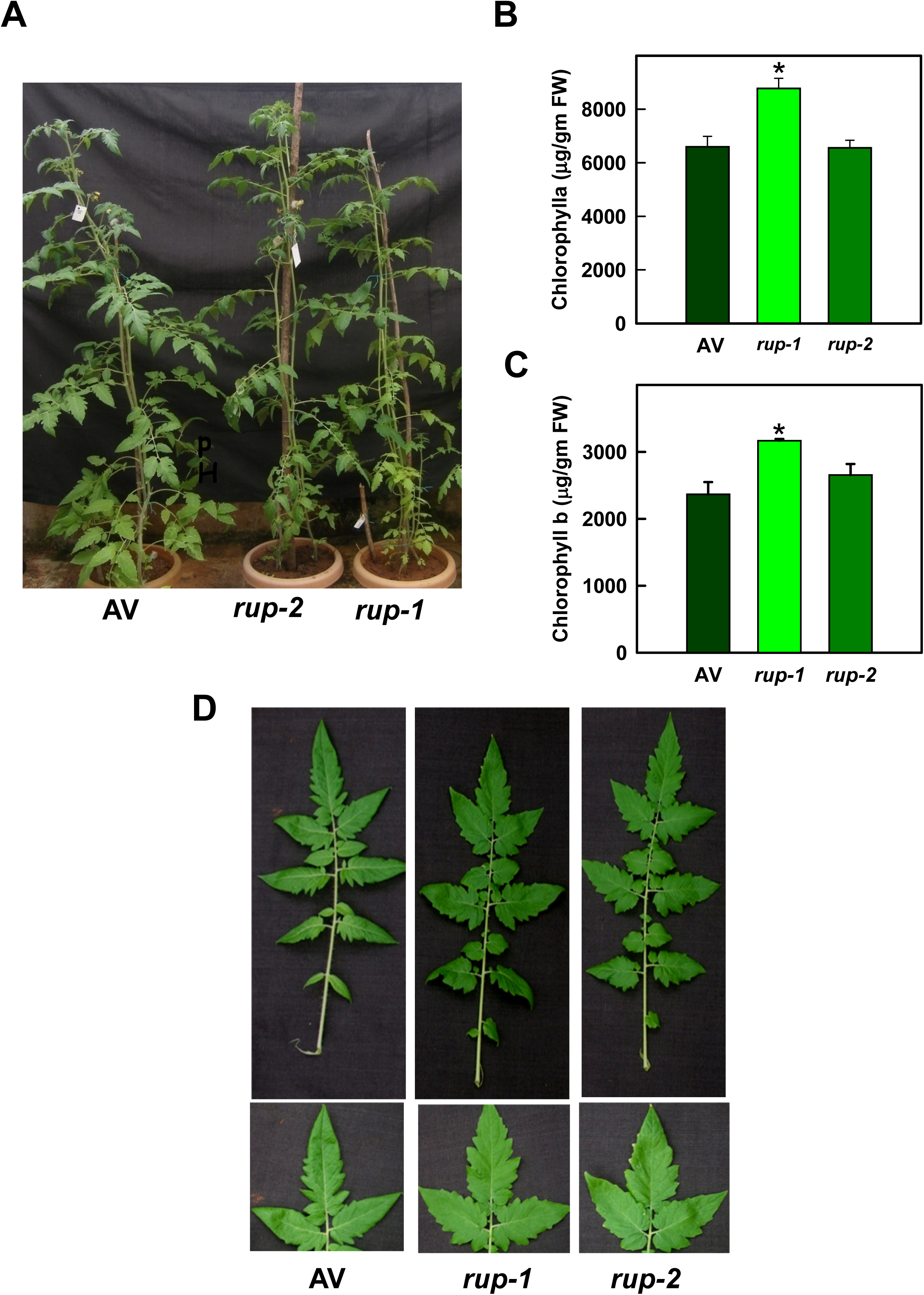
Vegetative phenotypes of AV and *rup-variants*. **A.** Two-month-old adult plant phenotypes of *rup-1* and *rup-2* compared to AV. **B-C.** Chlorophyll content (chlorophyll a and b) in adult plant leaves, with *rup-1* showing slightly higher levels of chlorophyll a and b compared to AV. **D.** Leaf morphology of *rup-1* and *rup-2* compared to AV. Each histogram represents the mean value ± SE (n ≥ 3). Asterisks indicate significant differences with P < 0.05 (*).

**Figure S10:**
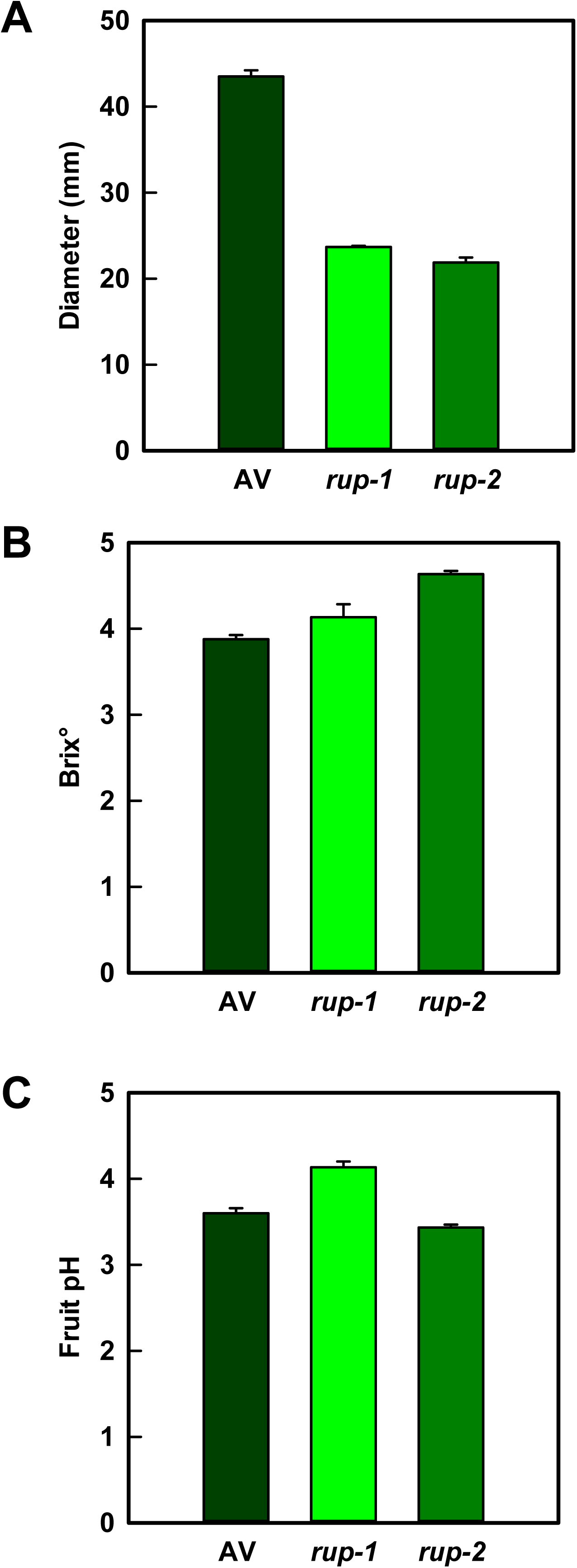
Comparison of chemical and physical parameters of red-ripe fruits of AV and *rup-*variants. **A.** Diameter of fruits. **B.** Total soluble solids content of the fruits**. C.** Acidity of the fruits.

**Figure S11.**
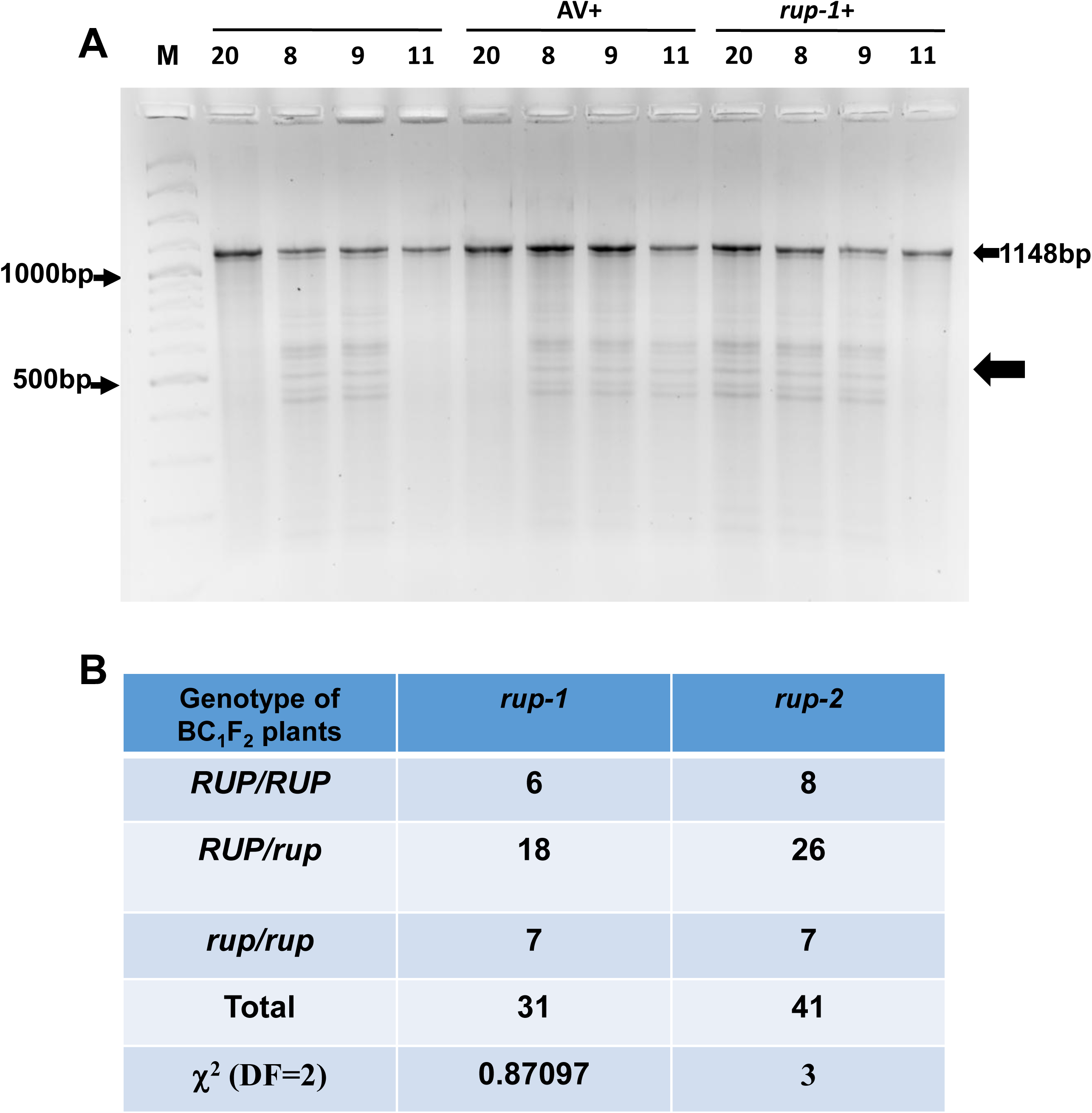
Genotype detection of AV (*RUP/RUP*) x *rup-1* (*rup-1/rup-1*) F_2_ plants using CEL I endonuclease assay. **A.** Representative image showing the use of CEL I assay to identify the genotype of BC_1_F_2_ plants. Genomic DNA from Arka Vikas (AV) and *rup-1* plants were PCR-amplified for the *RUP* gene, denatured, and renatured to form heteroduplex DNA, followed by CEL I digestion and gel electrophoresis. Cleavage products (Thick arrow **←**) generated by CEL I are observed only in heterozygous (*RUP/rup-1*) plants (# 8 and #9), while no cleavage is detected in homozygous plants (#20 and #11, left lanes). PCR and CEL I assays were performed with F_2_ plant DNA combined with either AV DNA (middle lanes, AV+) or *rup-1* DNA (right lanes, *rup-1+*), confirming that plant #20 is homozygous *RUP/RUP* and plant #11 is homozygous *rup-1/rup-1,* as indicated by the absence of cleavage products in the corresponding lanes. **B.** The number of BC_1_F_2_ plants analyzed using the CEL I assay and *RUP* and *rup-*variant genotype distribution. The BC_1_F_2_ plants segregated in a near-Mendelian (1:2:1) ratio for *rup-*variant, with a chi-square value indicating a single-trait genetic segregation. For the CEL I assay protocol. See Sreelakshmi et al. (2010) NEATTILL: A simplified procedure for nucleic acid extraction from arrayed tissue for TILLING and other high-throughput reverse genetic applications. *Plant Methods* 6: 3. **M**-DNA ladder.

**Figure S12.**
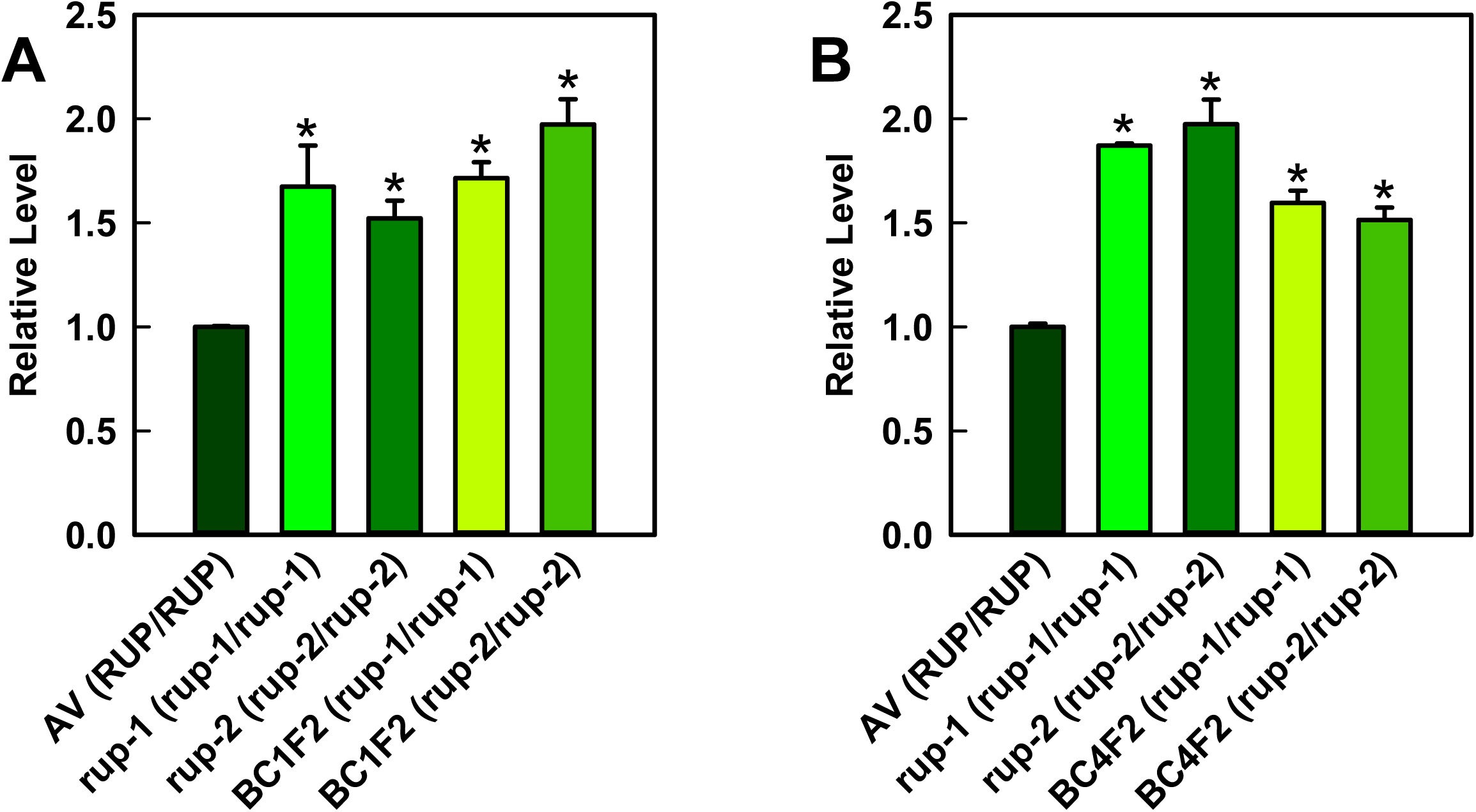
Relative levels of total carotenoids in red-ripe fruits of *rup*-variants and introgressed lines homozygous for the *rup* gene, compared to AV. **A.** Carotenoid levels in AV, *rup-*variants, and BC_1_F_2_ plants homozygous for *rup-1* (*rup-1/rup-1)* and *rup-2* (*rup-2/rup-2)*. **B.** Carotenoid levels in AV, *rup-*variants, and BC_4_F_2_ plants homozygous for *rup-1* (*rup-1/rup-1)* and *rup-2* (*rup-2/rup-2)*. Carotenoid data are presented as mean ± SE (n ≥ 3). Asterisks indicate significant differences with P < 0.05 (*).

**Figure S13.**
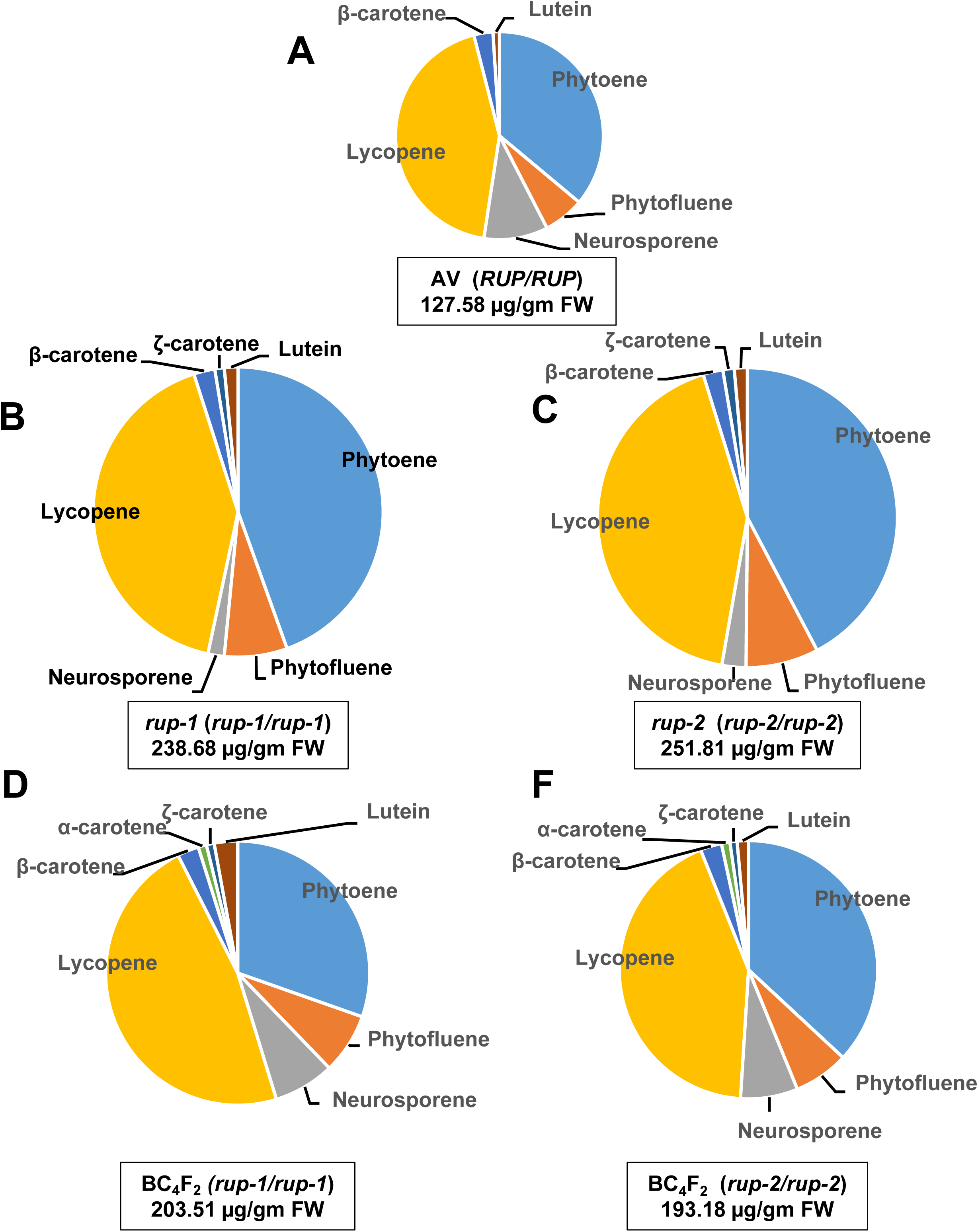
Distribution of carotenoids in red-ripe fruits of *rup*-variants, BC_4_F_2_ homozygous mutants, and Arka Vikas (AV). **A.** AV. **B.** *rup-1* (*rup-1/rup-1*) **C**. *rup-2* (*rup-2/rup-2*). **D.** BC_4_F_2_ (*rup-1/rup-1*). **E**. BC_4_F_2_(*rup-2/rup-2).* The proportional amounts of different carotenoids present in RR fruits are depicted in the pie diagram. Note that the total area of the pie is proportional to the total carotenoid amount. The values below the respective pie indicate the total carotenoid content (μg/gm FW). See **Dataset S1** for individual carotenoid levels and statistical significance.

**Figure S14.**
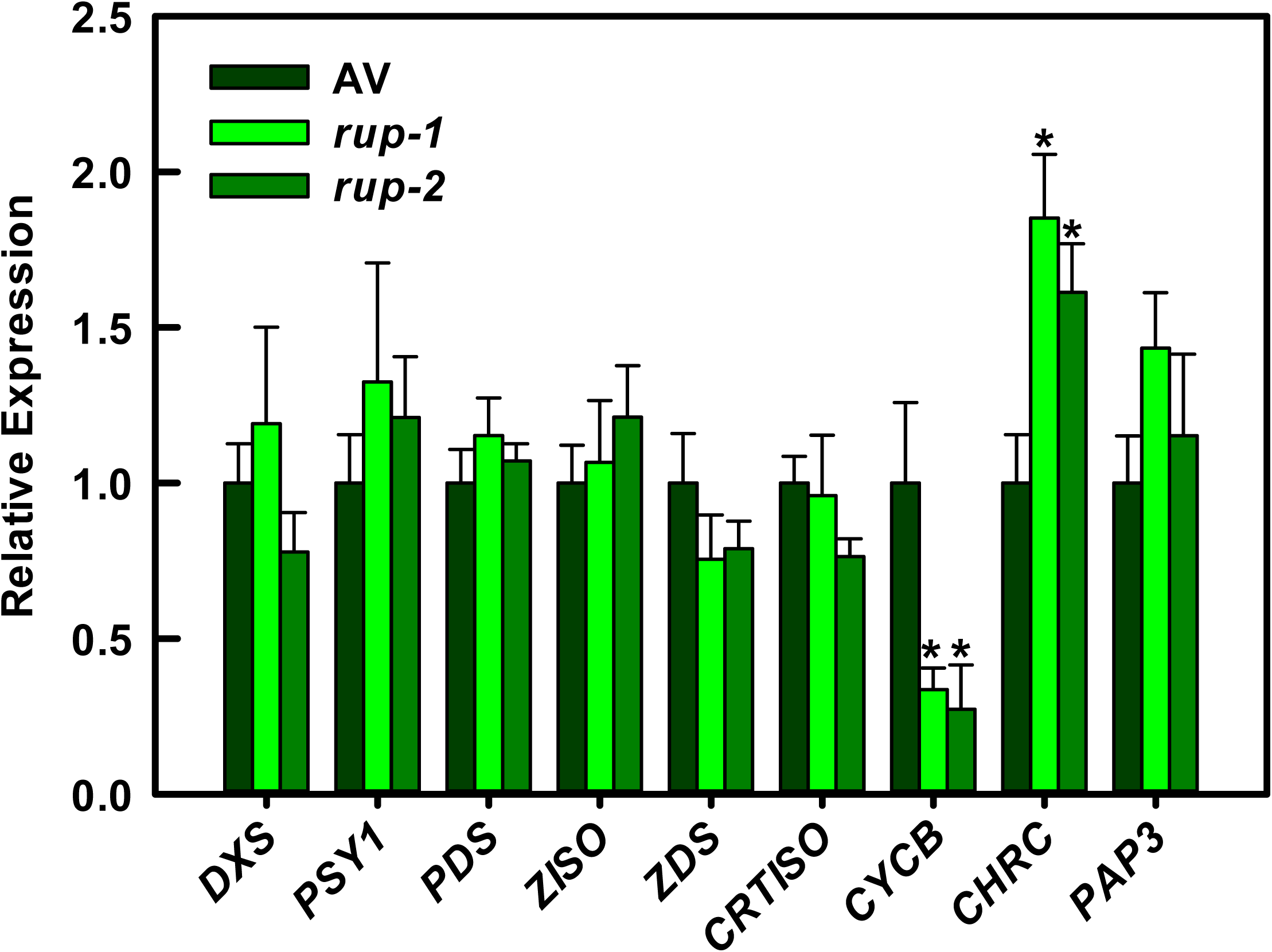
Transcript levels of carotenoid pathway genes in red-ripe fruits of *rup*-variants relative to AV. Data are presented as the ratio of expression in rup-variants relative to AV. **Abbreviation:** *CHRC-* Carotenoid-associated chromoplast protein; *CRTISO*-Carotenoid isomerase; *CYCB-* Lycopene β-cyclase; *DXS-* Deoxyxylulose 5-phosphate synthase; *PAP-* Plastid associated protein 3; *PDS-* Phytoene desaturase; *PSY-* Phytoene synthase; *ZDS-* ζ-carotene desaturase; *ZISO*-ζ-carotene isomerase. Each histogram represents the mean value ± SE (n ≥ 3). Asterisks indicate significant differences with P < 0.05 (*).

**Figure S15:**
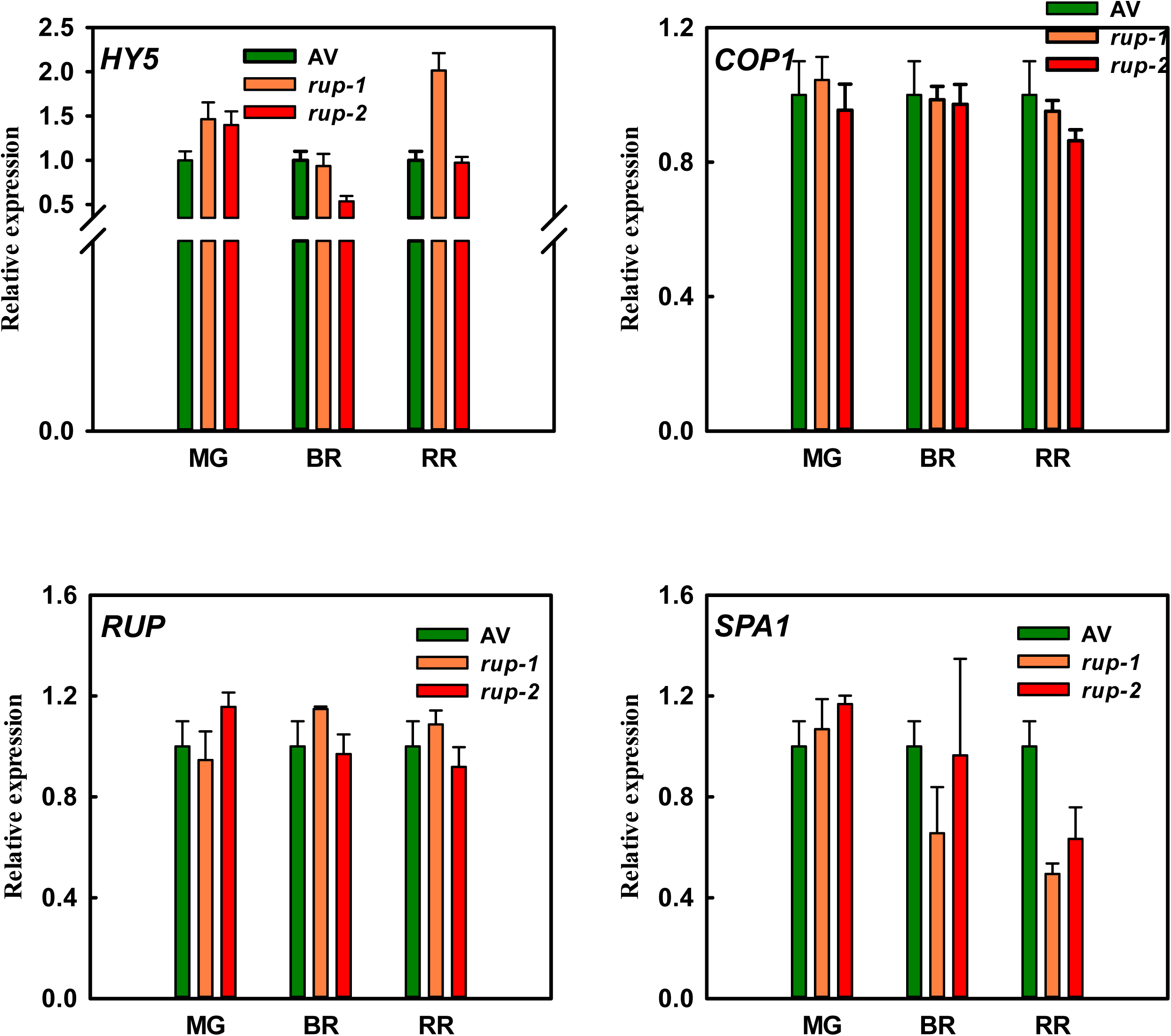
Expression of light-signaling genes during fruit ripening in *rup-*variants relative to AV. Data are presented as the ratio of expression in *rup*-variants relative to AV. **Abbreviation:** *COP1-* Constitutive Photomorphogenic 1*, HY5-* Elongated Hypocotyl 5*; RUP-* Repressor of UV Photomorphogenesis*, SPA1-* Suppressor of Phytochrome A. MG-Mature green, BR-Breaker, RR-Red ripe. Each histogram represents the mean value ± SE (n ≥ 3).

**Figure S16.**
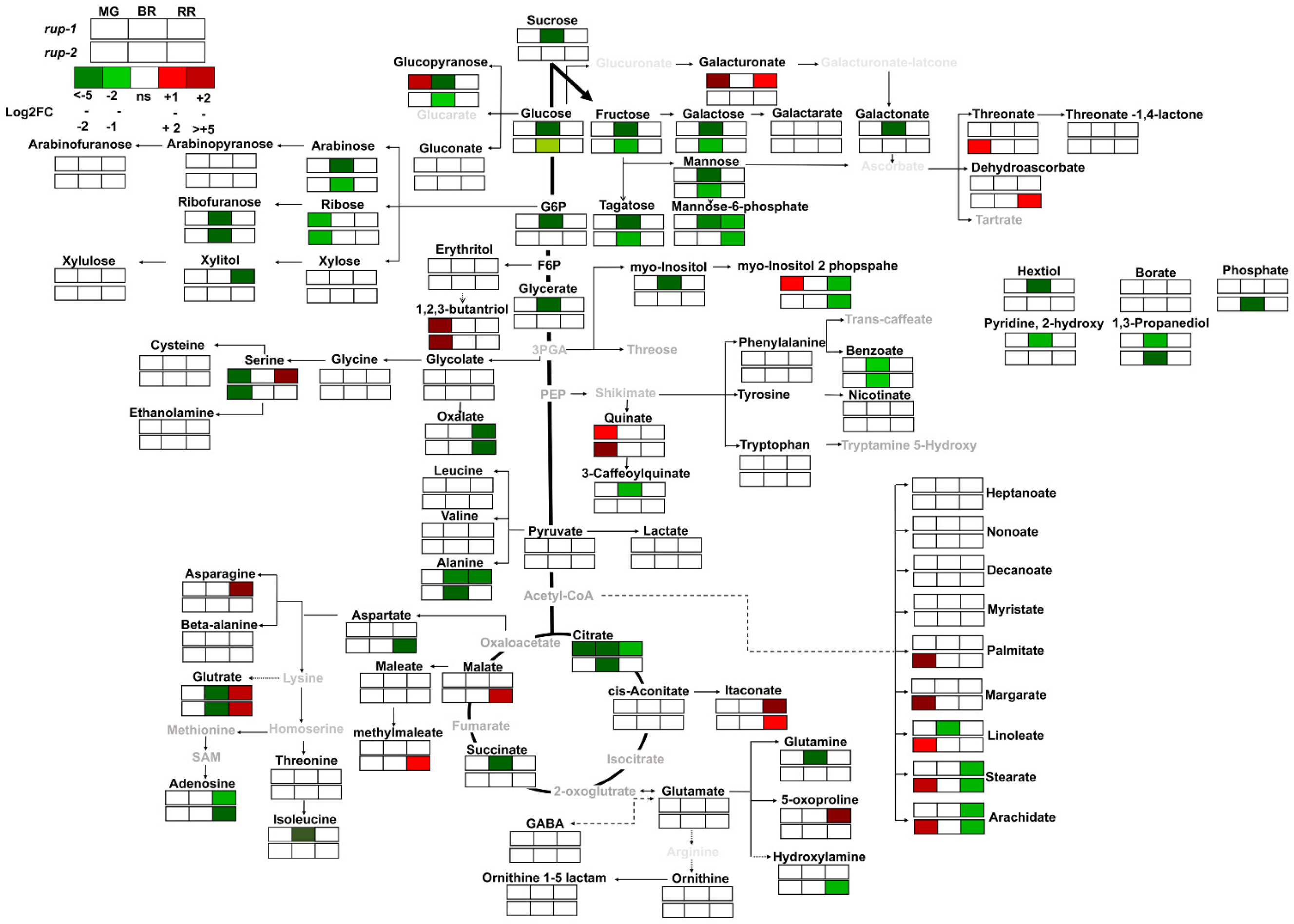
The levels of metabolites in *rup-*variant fruits at different ripening stages relative to AV. The relative changes in metabolite levels were calculated as the *rup*-variant/AV ratio for each ripening stage. The grid at the top left corner of the figure represents the ripening stages and the *rup-*variants used. Only metabolites with significant changes (log2 fold change ≥ ± 0.584; P-value ≤ 0.05; n ≥ 3) are displayed on the metabolic pathway. Identified metabolites are shown in bold black text, while metabolites below the detection limit are indicated in grey. White boxes represent no significant changes in metabolite levels. For detailed metabolite data, refer to **Dataset S2. MG**-Mature green, **BR**-Breaker, **RR**-Red ripe.

**Figure S17:**
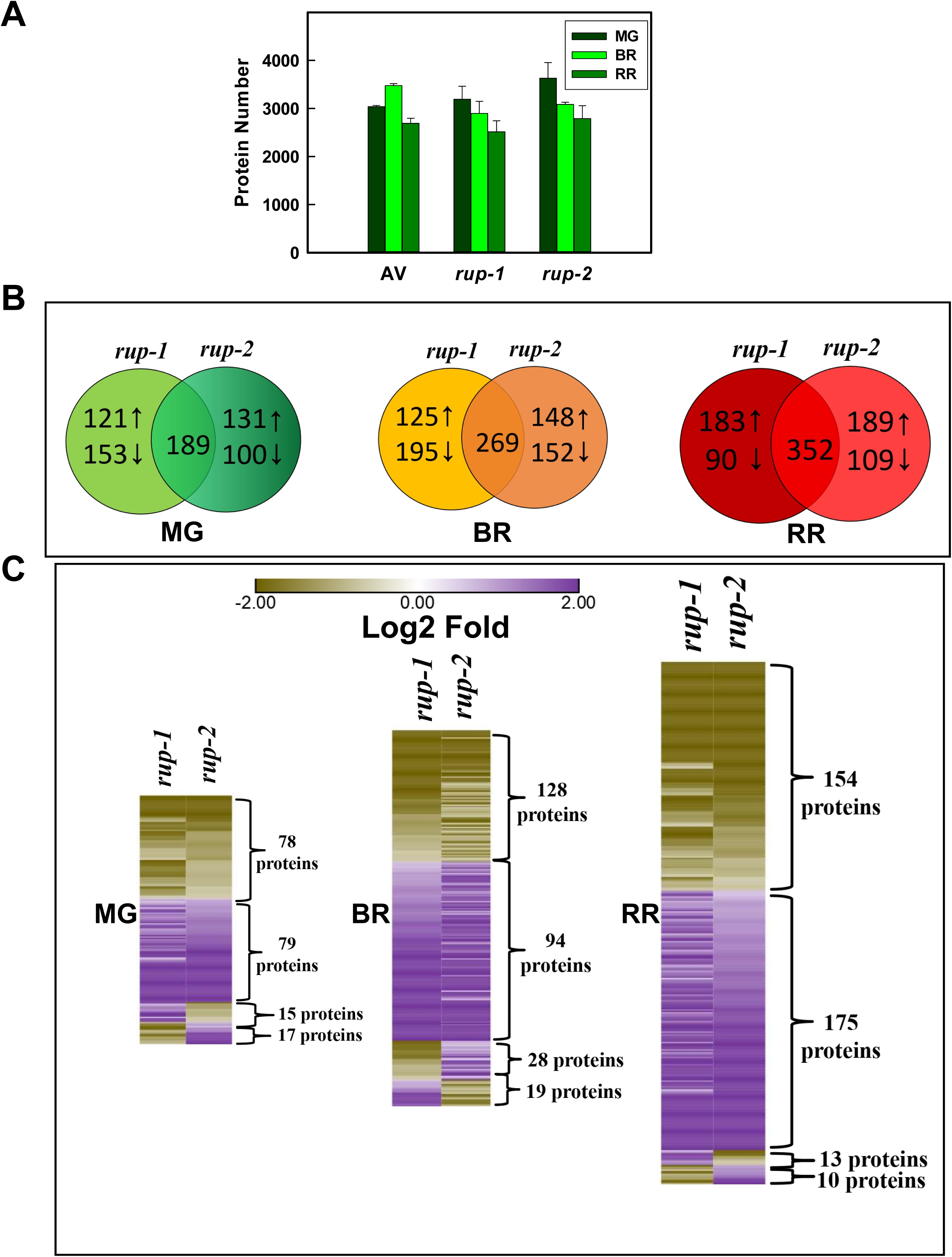
Differentially expressed proteins in *rup-*variants at different stages of fruit ripening relative to AV. **A.** Total number of detected proteins in *rup-1*, *rup-2,* and AV. **B.** Upregulated (↑) and down-regulated (↓) proteins in *rup-1* and *rup-2* compared to AV. The numbers in the intersection of circles indicate proteins common to *rup-1* and *rup-2*. The regulation pattern of these shared proteins is shown in the **C** panel**. C**. Regulation of overlapping proteins between *rup-1* and *rup-2.* Note: Some overlapping proteins exhibit opposite regulation. For details, see **Dataset S4.** Ripening stages: **MG-** mature green, **BR**-Breaker, **RR**-Red ripe.

**Figure S18:**
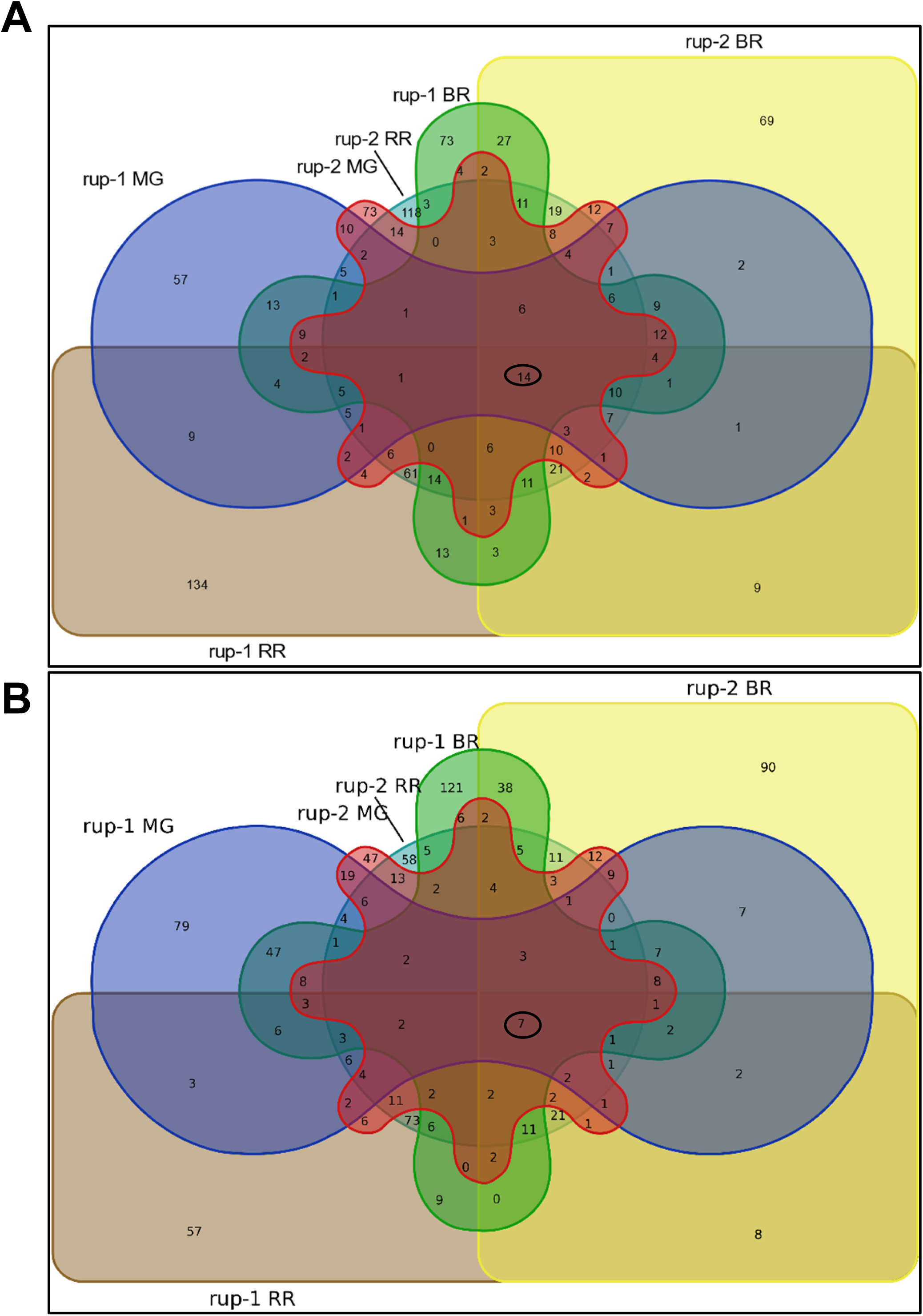
Six-set Venn diagram representation of differentially expressed proteins in *rup-1* and *rup-2* during fruit ripening. **A.** Upregulated proteins. **B.** Downregulated proteins. The numbers within shapes represent unique and overlapping proteins in *rup-*variants compared to Arka Vikas. Proteins commonly regulated across all six datasets are circled with a black line. **Abbreviations: MG**-mature green, **BR**-Breaker, **RR**-Red ripe. For details on specific unique and overlapping proteins in each group, see **Dataset S5.**

**Figure S19.**
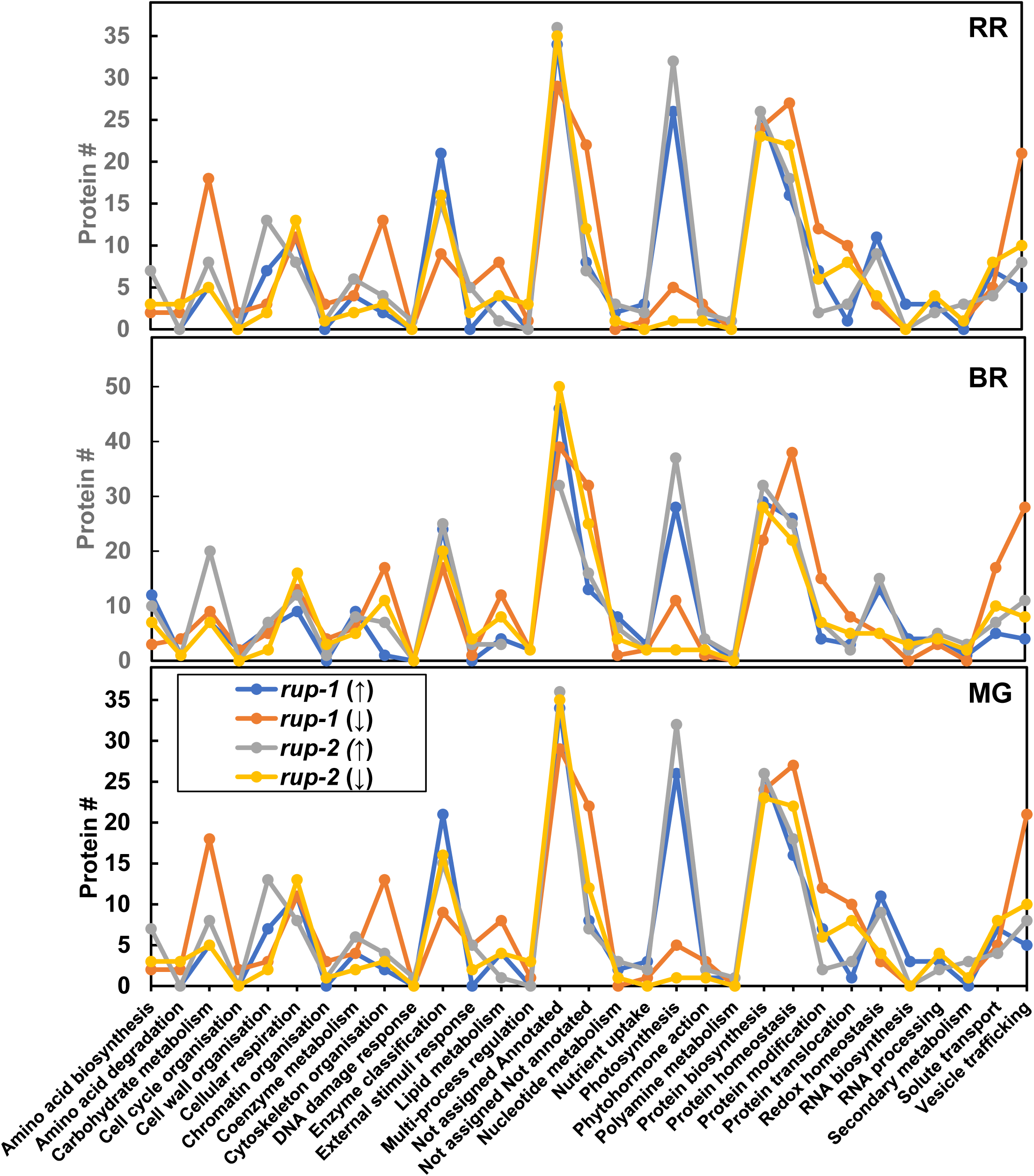
Functional classification of differentially regulated proteins in *rup-1* and *rup-2* during fruit ripening. Proteins were functionally categorized using Go-MapMan. In the figure legend, the upregulated proteins are indicated by upward arrows (↑), while downregulated proteins are marked with downward arrows (↓). For detailed protein data, see **Dataset S4**. **Abbreviations: MG**, Mature green; **BR**, Breaker; **RR**, Red ripe.

**Figure S20.**
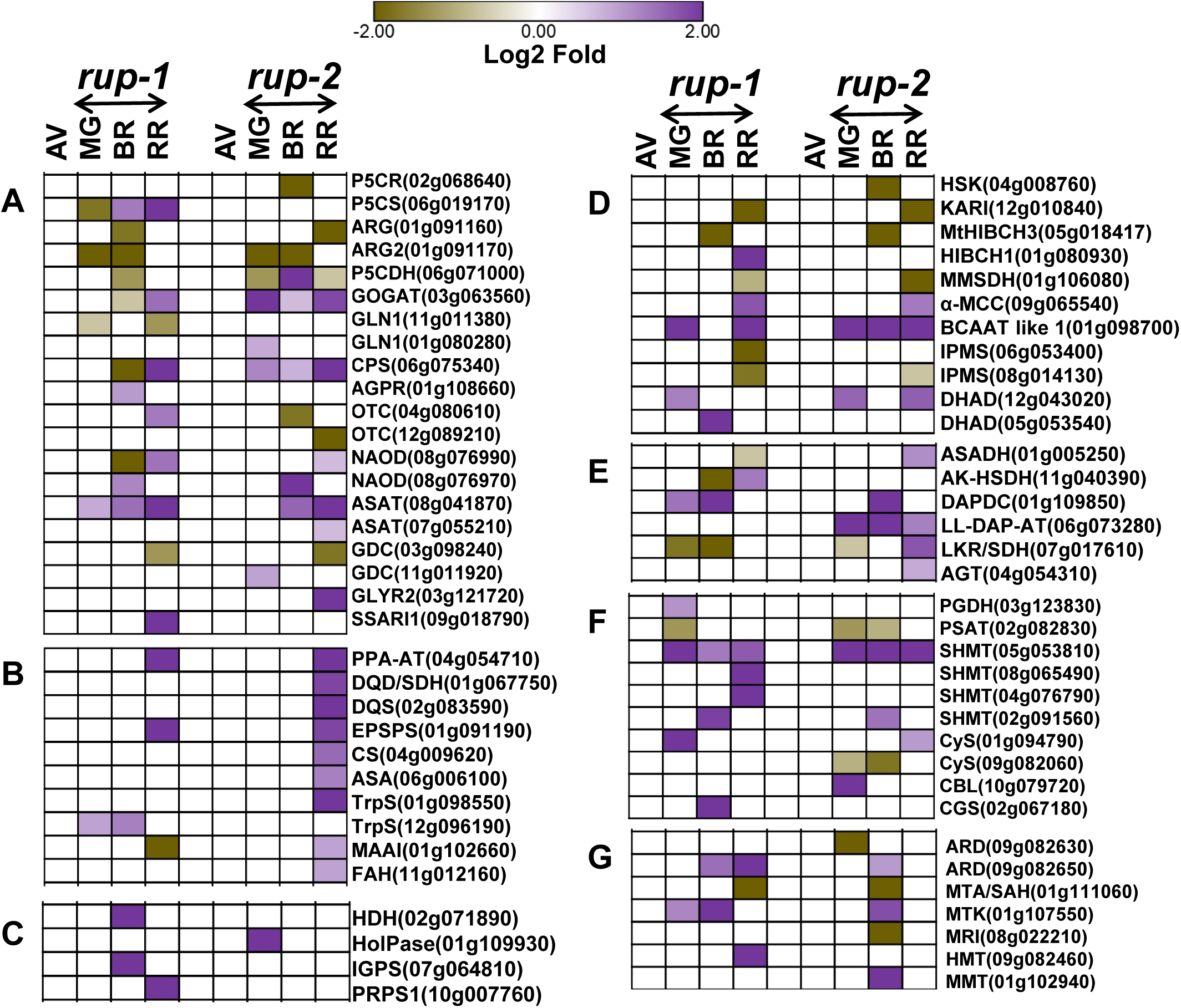
Differential regulation of amino acid metabolism-related proteins in *rup-1* and *rup-2* during fruit ripening relative to AV. Proteins related to (**A**) Glutamate, Glutamine, Proline, Arginine, (**B**) aromatic amino acids, (**C**) Histidine, (**D**) Leucine, Isoleucine, Valine, (**E**) Lysine. Alanine (**F)** Serine, Glycine, Cysteine, and (**G**) Methionine metabolism. Heat maps display significantly regulated proteins (Log2 fold change > ± 0.584, p < 0.05, n = 3) at different ripening stages. For better visualization of data, a lane between *rup-1* and *rup-2* has been left as blank. The SOLYC prefix before each protein ID is removed for convenience. For detailed proteome data, refer to **Dataset S6**. **Abbreviations:** AGPR-N-acetyl-gamma-glutamyl-phosphate reductase; AGT-Alanine-glyoxylate aminotransferase; AK-SHDH-Aspartokinase-homoserine dehydrogenase; ARD-Acidoreductone dioxygenase; ARG-Arginase; ASA-Anthranilate synthase; ASADH-Aspartate-semialdehyde dehydrogenase; ASAT-Aspartate aminotransferase; BCAAT-Branched-chain-amino-acid aminotransferase; CBL-Cystathionine β-lyase; CGS-Cystathionine γ-synthase; CPS-Carbamoyl-phosphate synthase; CS-Chorismate synthase; CYS-Cysteine synthase; DAPDC-Diaminopimelate decarboxylase; DHAD-Dihydroxy-acid dehydratase; DQD/ SDH-3-Dehydroquinate dehydratase and shikimate dehydrogenase; DQS-3-Dehydroquinate synthase; EPSPS-5-Enolpyruvyl-shikimate 3-phosphate synthase; FAH-Fumarylacetoacetase; GDC-Glutamate decarboxylase; GLYR-Glyoxylate reductase; GOGAT-Glutamate synthase; GLN1-Glutamine synthetase; HDH-Histidinol dehydrogenase; HIBCH1-3-Hydroxyisobutyryl-CoA hydrolase.1; HMT-Homocysteine S-methyltransferase; HolPase-Histidinol-phosphate phosphatase; HSK-Homoserine kinase; IGPS-Imidazole glycerol phosphate synthase; IPMS-Isopropylmalate synthase; KARI-Keto l-acid reductoisomerase; LKR/SDH-Bifunctional Lysine ketoglutarate reductase and Saccharopine dehydrogenase; LL-DAP-AT-LL-Diaminopimelate aminotransferase; MAAI-Maleylacetoacetate isomerase.; MMSDH-Methylmalonate-semialdehyde dehydrogenase; MTA/SAH- 5-methylthioadenosine/S-adenosylhomocysteine nucleosidase.1; MMT- Methionine S-methyltransferase; MtHIBCH3- 3-Hydroxyisobutyryl-CoA hydrolase-like protein 3 mitochondrial; MTK, Methylthioribose kinase; MRI- Methylthioribose-1-phosphate isomerase; NAOD- Acetylornithine deacetylase; OTC- Ornithine carbamoyltransferase; P5CDH- δ-1-pyrroline-5-carboxylate dehydrogenase; P5CR- Pyrroline-5-carboxylate reductase; P5CS- δ-1-Pyrroline-5-carboxylate synthase; PGDH- D-3-Phosphoglycerate dehydrogenase; PPA-AT- Prephenate aminotransferase; PRPS1- Ribose-phosphate pyrophosphokinase; PSAT- Phosphoserine aminotransferase; SHMT- Serine hydroxymethyltransferase; SSARI1- Succinic semialdehyde reductase isofom1; TrpS- Tryptophan synthase α chain; α- MCAA- α-Subunit of methylcrotonoyl-CoA carboxylase complex. **MG-** Mature green, **BR**- Breaker, **RR**- Red ripe.

**Figure S21.**
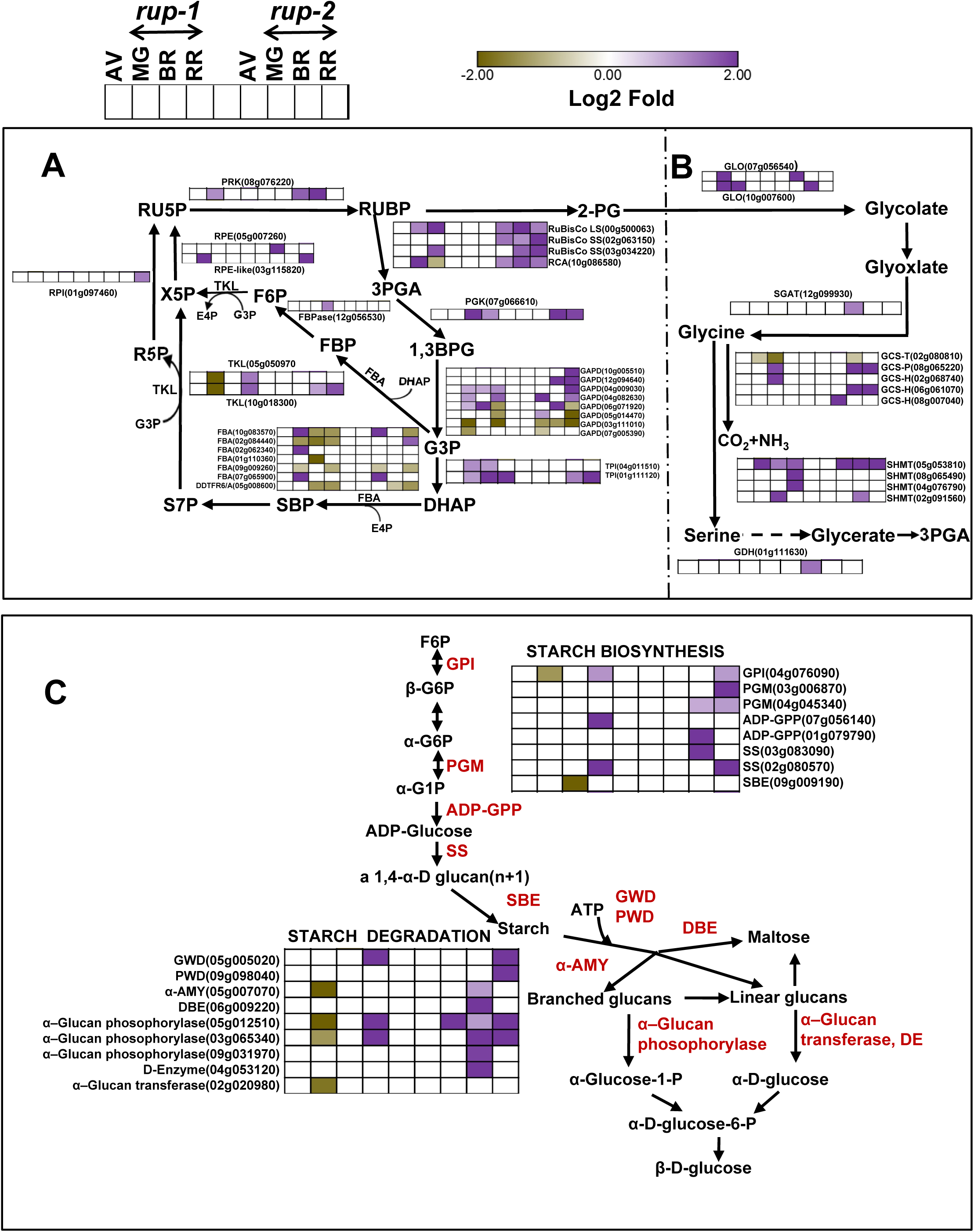
Differential regulation of carbohydrate metabolism-related proteins in *rup-1* and *rup-2* during fruit ripening relative to AV**. A**. Calvin cycle**. B**. Photorespiration. **C.** Starch metabolism. Heat maps display significantly regulated proteins (Log2 fold change > ± 0.584, p < 0.05, n = 3) at different ripening stages. For better visualization of data, a lane between *rup-1* and *rup-2* has been left as blank. The SOLYC prefix before each protein ID is removed for convenience. The dotted line in the top panel separates the A and B pathways. The cartoon on the top left corner depicts ripening stages used for the rup-variants relative to AV. For detailed proteome data, refer to **Dataset S6**. **Abbreviations:** 1,3BPG- 1,3-bisphosphoglyceric acid; 2-PG- 2-phosphoglycolate; 3PGA- 3-Phosphoglycerate; α-AMY- α-amylase; ADP-GPP- ADP-glucose pyrophosphorylase; DBE- Starch debranching enzyme; DDTFR6/A- Ripening regulated protein; DE- Disproportionating enzyme; DHAP- Dihydroxyacetone phosphate; E4P- Erythrose 4-phosphate; F6P- Fructose 6- phosphate; FBA- Fructose-bisphosphate aldolase; FBPASE- Fructose-16- bisphosphatase; FBP- Fructose 1,6-bisphosphate; G3P- Glyceraldehyde 3- phosphate; G6P- Glucose 6-phosphate; GAPD- Glyceraldehyde-3-phosphate dehydrogenase; GCS-H, GCS-T, GCS-P- Glycine cleavage system H, T, & P protein; GDH- Glycerate dehydrogenase; GPI- Glucose-6-phosphate isomerase; GWD- α-Glucan water dikinase; PGK- Phosphoglycerate kinase; PGM- Phosphoglucomutase protein; PGM-Phosphoglucomutase; PRK- Phosphoribulokinase; PWD- Phosphoglucan water dikinase; R5P- Ribose-5- phosphate; RCA- Ribulose bisphosphate carboxylase/oxygenase activase chloroplastic; RPE- Ribulose-phosphate 3-epimerase; RPI- Ribose 5-phosphate isomerase; RU5P- Ribulose 5-phosphate; RUBISCO- Ribulose bisphosphate carboxylase; RUBP- Ribulose bisphosphate; S7P- Sedoheptulose 7-phosphate; SBE- Starch Branching Enzyme; SBP- Sedoheptulose 1,7-bisphosphate; SGAT- Serine glyoxylate aminotransferase; SHMT- Serine hydroxymethyltransferase; SS- Starch synthase; TKL- Transketolase; TPI- Triosephosphate isomerase; X5P- Xylulose 5-phosphate. **MG**- Mature green, **BR**- Breaker, **RR**- Red ripe.

**Figure S22.**
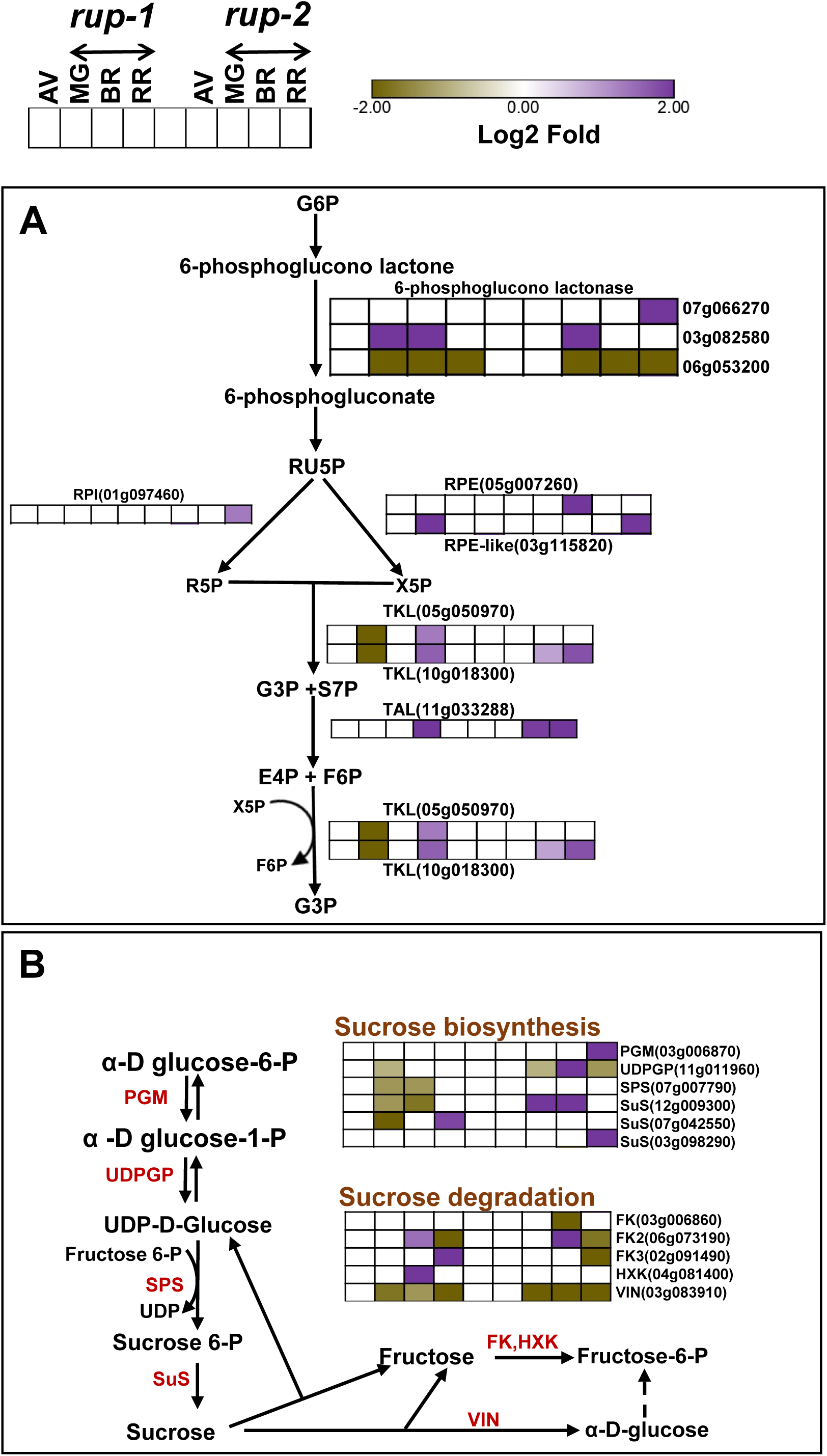
Differential regulation of carbohydrate metabolism-related proteins in *rup-1* and *rup-2* during fruit ripening relative to AV. **A**. Pentose phosphate pathway. **B.** Sucrose metabolism. Heat maps display significantly regulated proteins (Log2 fold change > ± 0.584, p < 0.05, n = 3) at different ripening stages. For better visualization of data, a lane between *rup-1* and *rup-2* has been left as blank. The SOLYC prefix before each protein ID is removed for convenience. The cartoon in the top left corner depicts ripening stages used for the *rup-*variants relative to AV. For detailed proteome data, refer to **Dataset S6. Abbreviations:** E4P- Erythrose 4-phosphate; F6P- Fructose 6-phosphate; FK- Fructose kinase; G3P- Glyceraldehyde 3-phosphate; G6P-Glucose 6- phosphate; HXK- Plastidic hexokinase; PGM- Phosphoglucomutase protein; R5P- Ribose-5-phosphate; RPE- Ribulose-phosphate 3-epimerase; RPI-Ribose 5-phosphate isomerase; RU5P- Ribulose 5-phosphate; S7P- Sedoheptulose 7-phosphate; SPS- Sucrose phosphate synthase; SuS- Sucrose synthase; TAL-Transaldolase; TKL- Transketolase; UDPGP- UDP glucose pyrophosphorylase; VIN- Vacuolar invertase/ Acid β-fructofuranosidase; X5P- Xylulose 5-phosphate. **MG-**Mature green, **BR-** Breaker, **RR-** Red ripe.

**Figure S23.**
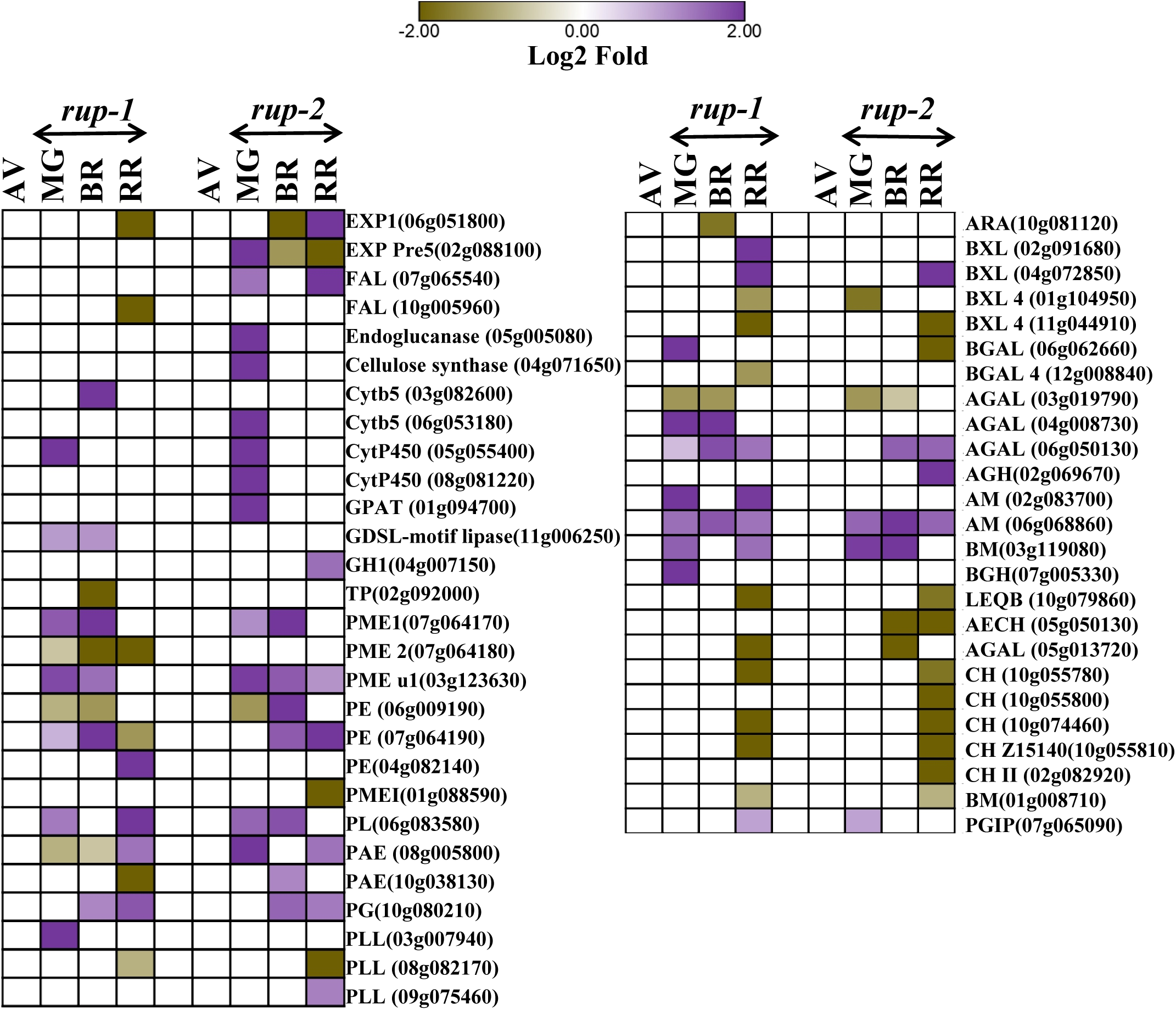
Differential regulation of the cell wall components and cell wall modification-related proteins in *rup-1* and *rup-2* during fruit ripening relative to AV. Heat maps display significantly regulated proteins (Log2 fold change > ± 0.584, p < 0.05, n = 3) at different ripening stages. For better visualization of data, a lane between *rup-1* and *rup-2* has been left as blank. The SOLYC prefix before each protein ID is removed for convenience. For detailed proteome data, refer to **Dataset S6. Abbreviations:** AECH-Acidic endochitinase; AGAL- α-galactosidase; AGAL- α- galactosidase; AGH- α-glucosidase; AM- α-mannosidase; ARA- α-L-arabinofuranosidase; BGAL- β-galactosidase; BGH- Glucan endo-13-β-glucosidase; BM- β-mannosidase enzyme; BXL- β-D-xylosidase; CH- Chitinase; Cytb5-Cytochrome B5; CytP450-Cytochrome P450; EXP- Expansin; FAL- Fasciclin-like arabinogalactan protein.; GH1- Glycoside hydrolase family 1; GPAT- Glycerol-3-phosphate acyltransferase; LEQB- L. esculentum TomQ’b β-glucanase; PAE- Pectin acetyl esterase; PE- Pectin esterase; PG- Polygalactouronase; PL- Pectate lyase; PLL- Pectate lyase like superfamily protein; PME- Pectin methylesterase; PMEI- Pectinesterase inhibitor TP-Transmembrane protein. **MG-** Mature green, **BR-** Breaker, **RR-** Red ripe.

**Figure S24.**
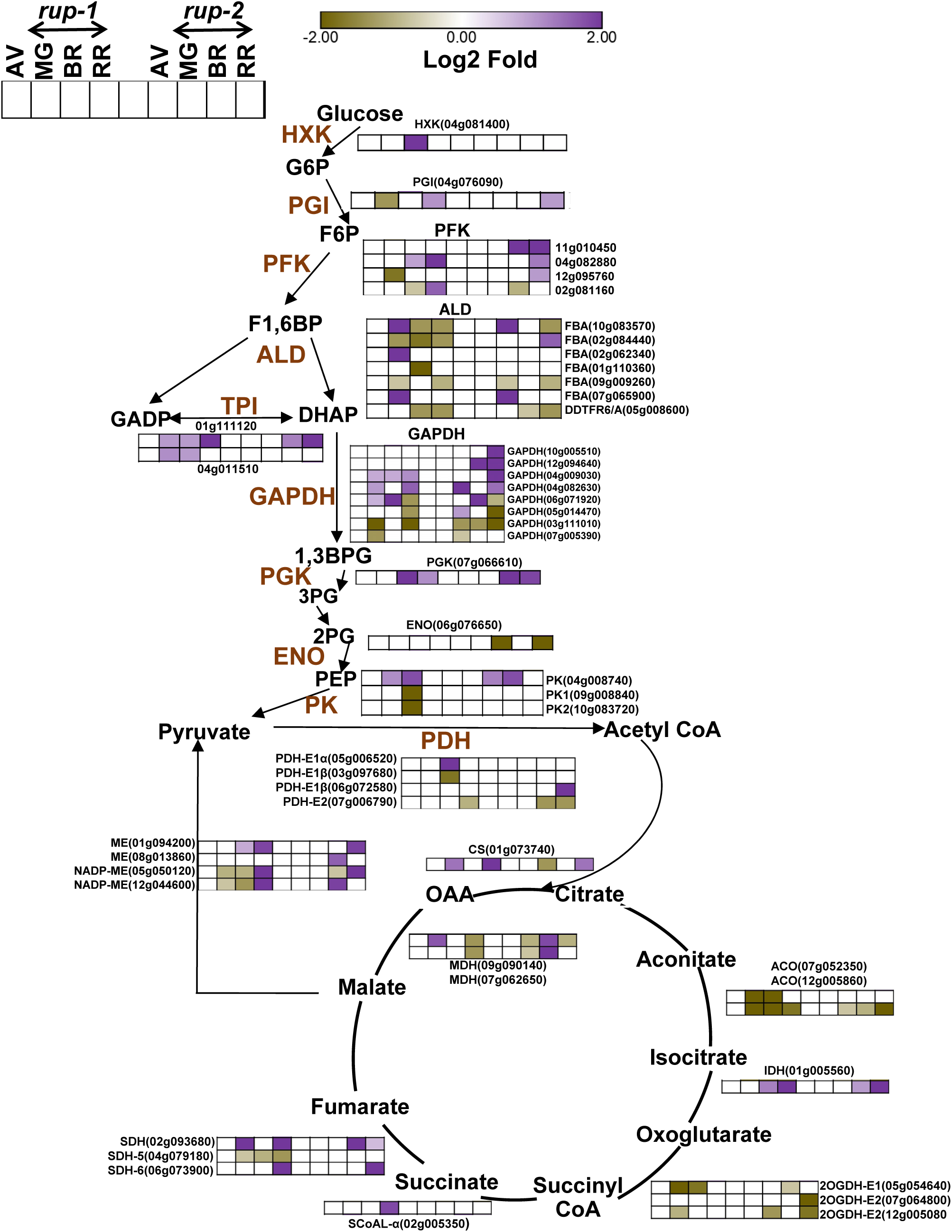
Differential regulation of the glycolysis and citric acid cycle metabolism-related proteins in *rup-1* and *rup-2* during fruit ripening relative to AV. Heat maps display significantly regulated proteins (Log2 fold change > ±0.584, p <0.05, n=3) at different ripening stages. For better visualization of data, a lane between *rup-1* and *rup-2* has been left as blank. The SOLYC prefix before each protein ID is removed for convenience. The cartoon in the top left corner depicts ripening stages used for the *rup*-variants relative to AV. For detailed proteome data, refer to **Dataset S6. Abbreviations:** 1,3BPG- d-1,3-Bisphosphoglycerate; 2OGDH-E1- Component E1 of 2-oxoglutarate dehydrogenase complex; 2OGDH-E2- Component E2 of 2-oxoglutarate dehydrogenase complex; 2PG- 2-Phosphoglyceric acid; 3PG-3-Phosphoglyceric acid; ACO- Aconitase; ALD- Aldolase; CS- Citrate synthase; DHAP- Dihydroxyacetone phosphate; ENO- Enolase; FBA- Fructose-bisphosphate aldolase; F1,6BP- Fructose 1,6-bisphosphate; F6P- Fructose 6-phosphate; GADP- Glyceraldehyde 3-phosphate; GAPDH- Glyceraldehyde-3- phosphate dehydrogenase; GPI- Glucose-6-phosphate isomerase; G6P-Glucose 6-phosphate; HXK- Plastidic hexokinase; IDH- Isocitrate dehydrogenase; MDH - Malate dehydrogenase; ME- Malic enzyme; NADP-ME- Cytosolic NADP-dependent malic enzyme; OAA- Oxaloacetic acid; PDH-e1α, PDH-e1β- Pyruvate dehydrogenase complex component E1 heterodimer subunit α, β; PDH-E2- Mitochondrial pyruvate dehydrogenase complex component E2; PEP- Phosphoenolpyruvic acid; PFK- ATP-dependent 6-phosphofructokinase; PGI- phosphoglucose isomerase; PGK- Phosphoglycerate kinase; PK- Pyruvate kinase; SCoAL- α- Subunit alpha of succinyl-coA ligase; SDH2- Succinate dehydrogenase 2; SDH-5- Component SDH5 of succinate dehydrogenase complex; SDH6- Component SDH6 of succinate dehydrogenase complex; TPI- Triosephosphate isomerase. **MG**- Mature green, **BR**- Breaker, **RR**- Red ripe.

**Figure S25.**
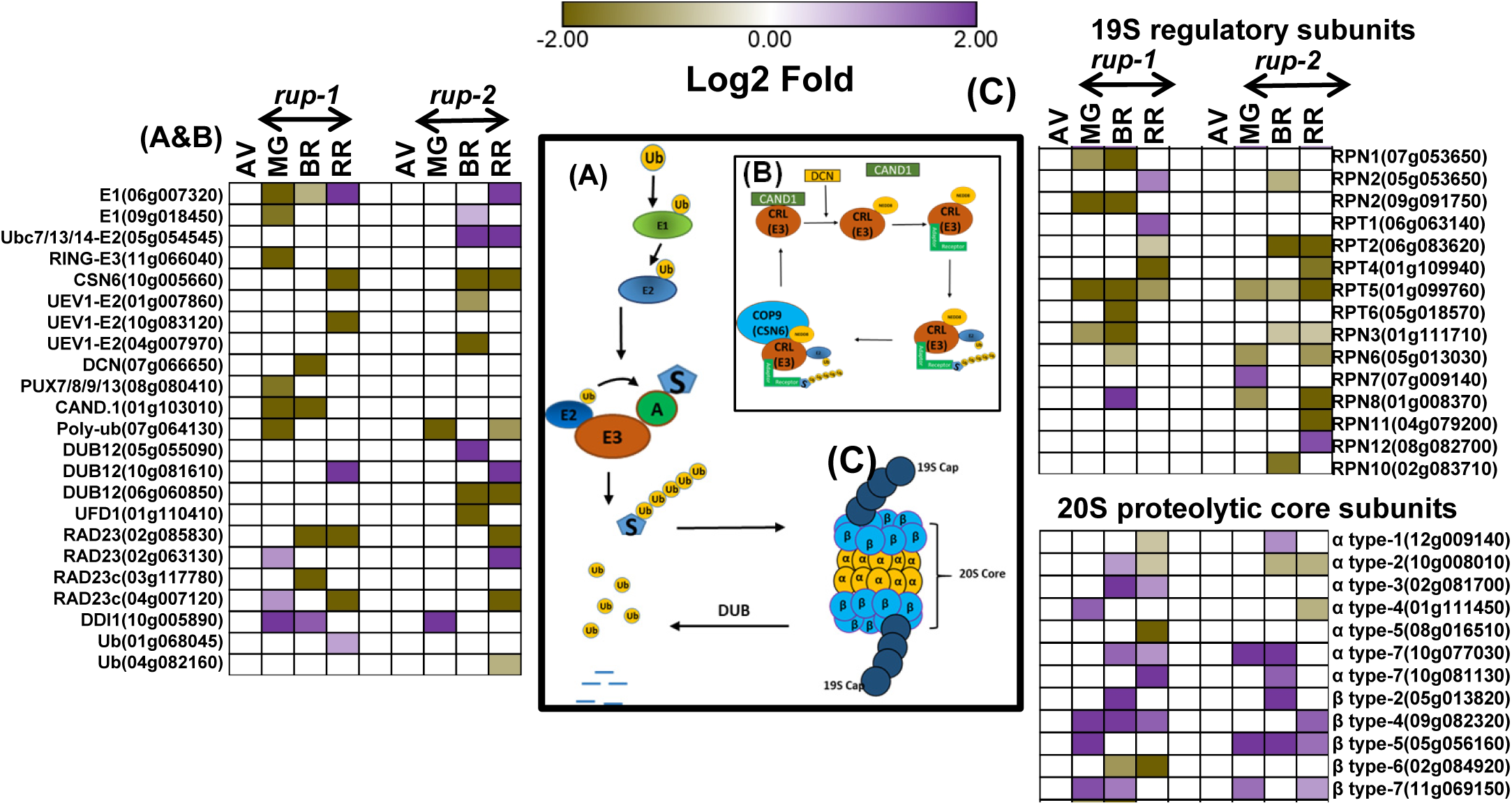
Differential regulation of ubiquitin-proteasome system-related proteins in *rup-1* and *rup-2* during fruit ripening relative to AV. **A.** Ubiquitin-26s proteasome-mediated protein degradation. **B.** Cullin ring E3 ligase (CRL) regulation by NEDD8, DCN, and COP9 signalosome. **C.** 26S Proteosome-related components. Heat maps display significantly regulated proteins (Log2 fold change > ± 0.584, P < 0.05, n = 3) at different ripening stages. The center panel shows the composition of (**A**), (**B**), and (**C**) and the steps involved in protein degradation. For better visualization of data, a lane between *rup-1* and *rup-2* has been left as blank. The SOLYC prefix before each protein ID is removed for convenience.. For detailed proteome data, refer to **Dataset S6. Abbreviation:** A-Adaptor; CAND.1-Cullin-associated NEDD8-dissociated protein.1 (CAND.1-adaptor protein exchange factor); CRL-Cullin/RING ubiquitin ligases; CSN6-Component CSN6 of COP9 signalosome complex; DCN-Defective in cullin neddylation protein; DDI1-DDI1-like//DNA-damage-inducible protein ddi1; DUB-deubiquitinase; E1-Ubiquitin-activating E1 protein; NEDD8-Neural cell expressed developmentally down-regulated protein 8;; Poly-ubi-Polyubiquitin; PTRE1-Proteasome inhibitor-related protein; PUX7/8/9/13-Plant UBX domain-containing protein 8 (PUX7/8/9/13)-adaptor protein); RAD23-Shuttle protein; RAD23c-Proteasome adaptor protein; RING-E3-E3 ubiquitin ligase RING-type E3 ubiquitin transferase; RPN-26S proteasome non-ATPase regulatory subunit (RPN1, RPN2, RPN3, RPN6, RPN7, RPN8, RPN10, RPN11, RPN12); RPT-26S protease regulatory subunit 7 (RPT1, RPT2, RPT4, RPT5, RPT6); S-Substrate; Ub-Ubiquitin; Ubc7/13/14-E2-Ubiquitin-conjugating component Ubc7/13/14 (E)2 of HRD1 E-3 Ubiquitin ligase complex (HRD1 E3-Ubc7/13/14); UEV1-E2-Component UEV1 of Ubc13-Uev1 conjugating E2 complex; UFD1-Component UFD1 of ER-associated protein degradation machinery. **MG-** Mature green, **BR-** Breaker, **RR**-Red ripe.

**Figure S26.**
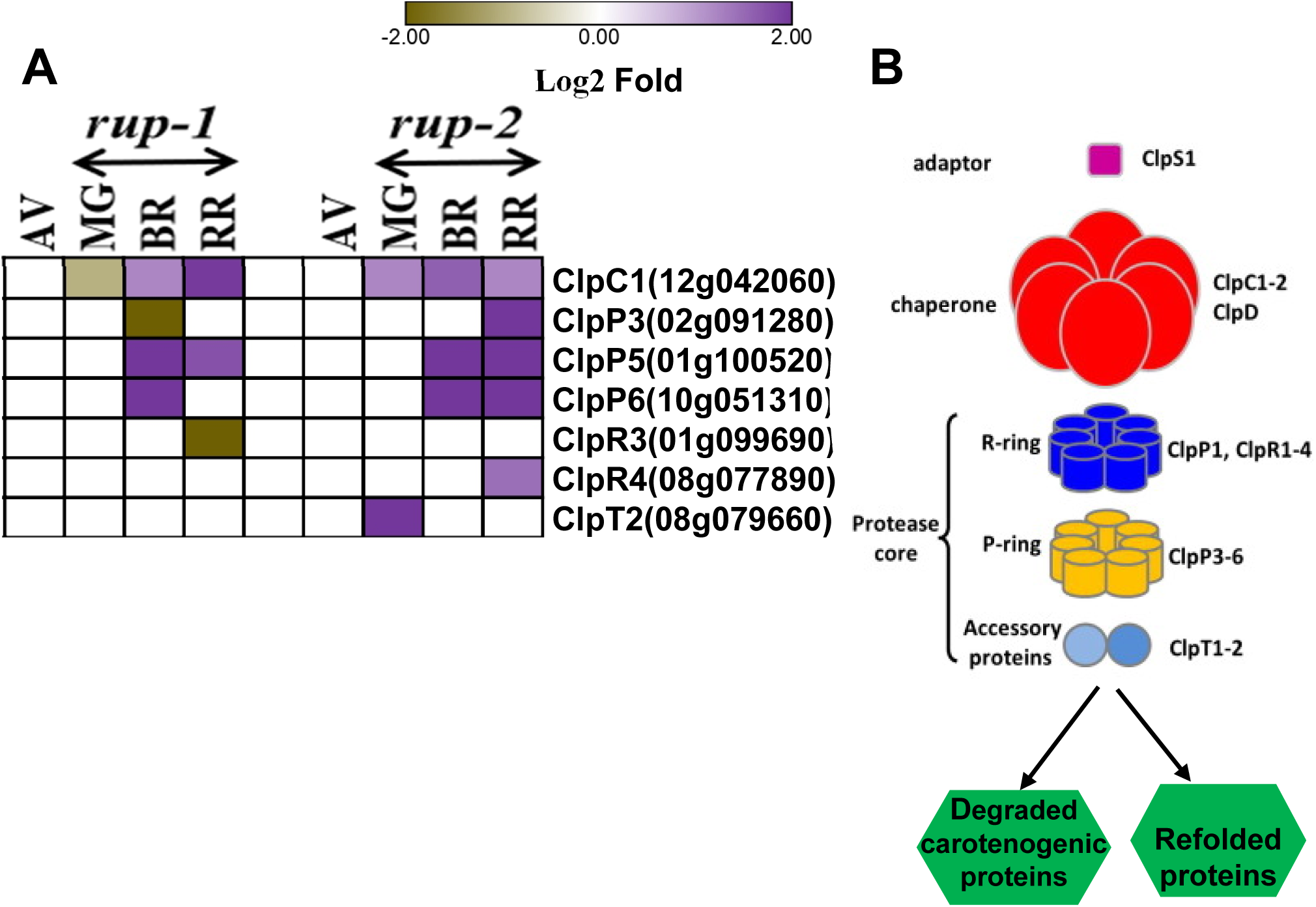
Differential regulation of Clp proteolytic complex related proteins in *rup-1* and *rup-2* relative to AV during fruit ripening. **A.** Differential regulation of Clp proteolytic components. **B.** Clp proteolytic system. Heat maps display significantly regulated proteins (Log2 fold change > ± 0.584, p < 0.05, n = 3) at different fruit ripening stages. For better visualization of data, a lane between *rup-1* and *rup-2* has been left as blank. The SOLYC prefix before each protein ID is removed for convenience. For detailed proteome data, refer to **Dataset S6**. **Abbreviations:** ClpC1-ATP-dependent Clp protease ATP-binding subunit; ClpP3/ClpR3/ClpR4-ATP-dependent Clp protease proteolytic subunit; ClpT2-ATP-dependent Clp protease ATP-binding subunit ClpC; ClpP5-Clp protease 2 proteolytic subunit; ClpP6-Clp-like energy-dependent protease; **MG**-Mature green, **BR**-Breaker, **RR**-Red ripe.

**Figure S27.**
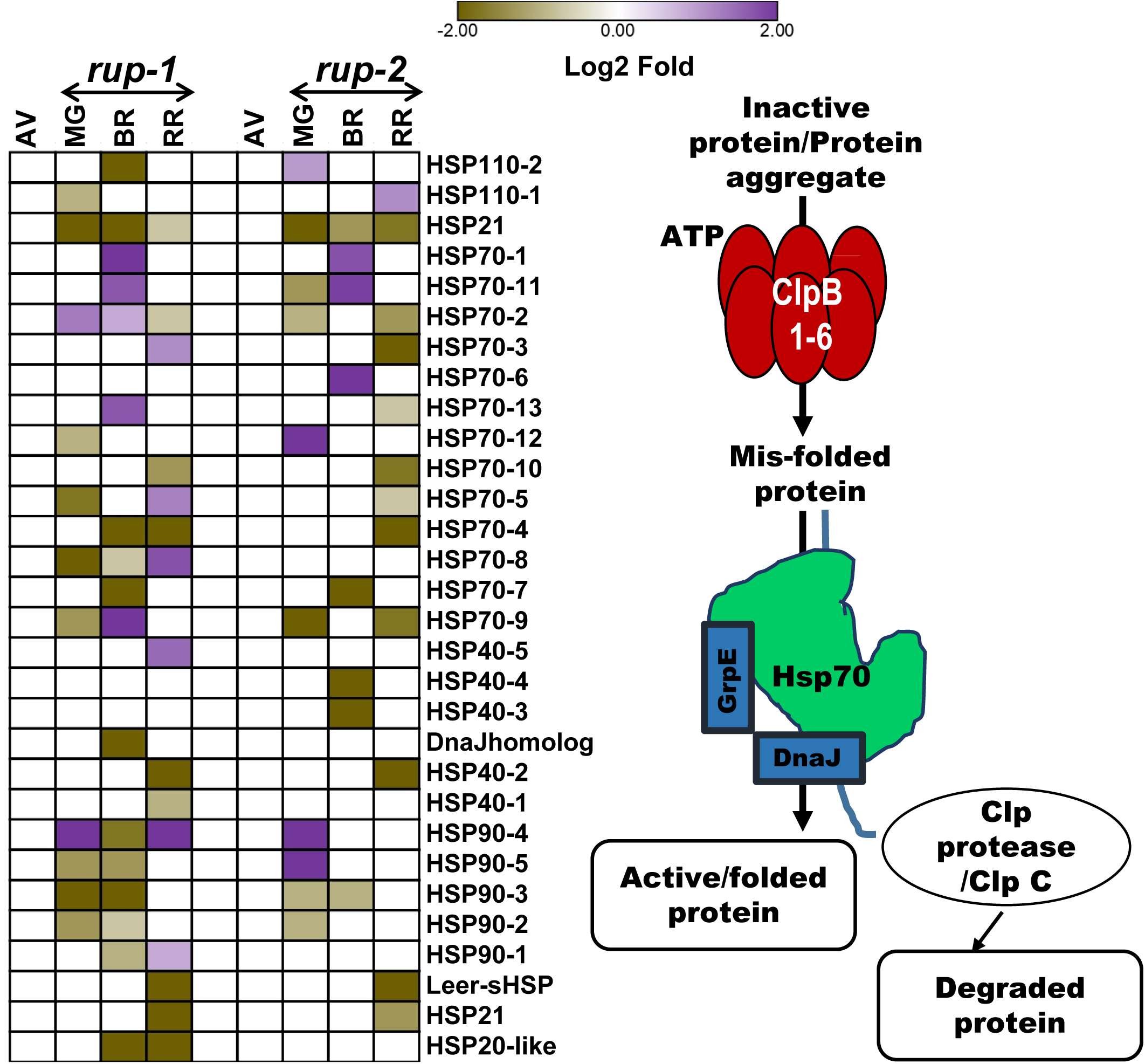
Differential regulation of chaperone system-related heat shock proteins in *rup-*1 and *rup-2* relative to AV. **A.** Relative levels of differentially regulated HSPs. **B**. Cartoon showing the role of HSPs in protein protection. The HSP are numbered in ascending order of their respective Solyc ID from chromosomes 1-12. Heat maps display significantly regulated proteins (Log2 fold change > ± 0.584, p < 0.05, n = 3) at different fruit ripening stages. For better visualization of data, a lane between *rup-1* and *rup-2* has been left as blank. For detailed proteome data, refer to **Dataset S6**. **Abbreviations:** HSP-Heat shock protein; NEF-Nucleotide exchange factor; DnaJ homolog-Solyc08g021920; HSP20-like-Solyc09g075010; HSP21-Solyc03g082420; HSP40-1/DnaJ-Solyc01g105340; HSP40-2/DnaJ-Solyc02g077670; HSP40-3/DnaJ - Solyc04g009770; HSP40-4/DnaJ - Solyc05g055160; HSP40-5/DnaJ-Solyc11g006460; HSP70-1-Solyc01g103450; HSP70-2-Solyc01g106210; HSP70-3-Solyc01g106260; HSP70-4-Solyc03g117630; HSP70-5-Solyc04g011440; HSP70-6-Solyc06g052050; HSP70-7/NEF-Solyc07g043560; HSP70-8-Solyc09g010630; HSP70-9/NEF-Solyc09g011030; HSP70-10-Solyc09g075950; HSP70-11-Solyc11g020040; HSP70-12-Solyc11g066060; HSP70-13-Solyc11g066100; HSP90-1-Solyc04g081570; HSP90-2-Solyc06g036290; HSP90-3-Solyc07g047790; HSP90-4-Solyc07g065840; HSP90-5-Solyc12g015880; HSP100-1-Solyc03g115230; HSP110-1/ClpB1-Solyc02g088610; HSP110-2-Solyc12g043120; Leer-sHSP-Solyc11g020330. **MG**-Mature green; **BR**-Breaker; **RR**-Red ripe.

**Figure S28.**
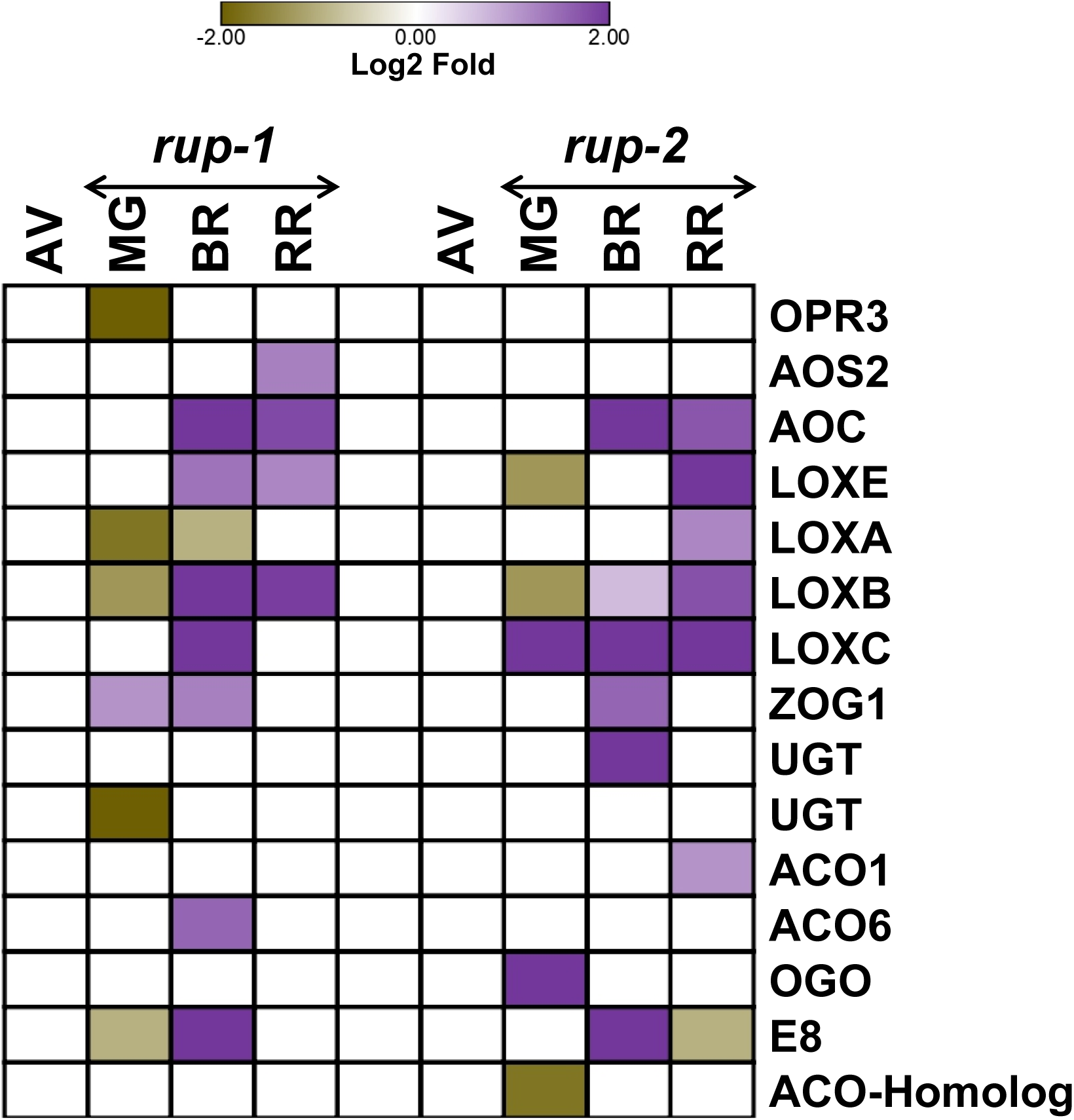
Differential regulation of the hormone metabolism-related proteins in *rup-1* and *rup-2* relative to AV. Heat maps display significantly regulated proteins (Log2 fold change > ± 0.584, p < 0.05, n = 3) at different fruit ripening stages. For better visualization of data, a lane between *rup-1* and *rup-2* has been left as blank. For detailed proteome data, refer to **Dataset S6 Abbreviations:** ACO-1-aminocyclopropane-1-carboxylate oxidase, ACO1 (Solyc07g049530), ACO6 (Solyc02g036350), ACO Homolog (Solyc04g007980); AOC-Allene oxide cyclase (Solyc02g085730); AOS2-Allene oxide synthase 2 (Solyc11g069800); E8 (Solyc09g089580); LOX-Lipoxygenase, LOXA (Solyc08g014000), LOXB (Solyc01g099190), LOXC (Solyc01g006540), LOXE (Solyc01g099160); OGO- 2OG-Fe(II) oxygenase (Solyc02g093080); OPR3- 12-oxophytodienoate reductase3 (Solyc07g007870); UGT- UDP-glycosyltransferase (Solyc10g085870), (Solyc07g043490; ZOG- Zeatin O-glucosyltransferase (Solyc04g016220). **MG**- Mature green, **BR**- Breaker, **RR-** Red ripe.

**Dataset S1.** Carotenoids determination in *rup*-variants and during backcrossing with AV.

**Dataset S2.** List of metabolites identified at MG, BR, and RR stages of Arka Vikas, *rup-1*, and *rup-2*.

**Dataset S3.** List of metabolites identified at RR stages of Arka Vikas, *rup-1, rup-2,* and backcrossed fruits.

**Dataset S4**. The differentially regulated proteins in Arka Vikas, *rup-1*, and *rup-2* at different stages of fruit ripening.

**Dataset S5.** List of shared and unique proteins shown in six-set Venn diagram.

**Dataset S6.** Differentially expressed proteins of *rup-1* and *rup-2* depicted in the Figures.

**Dataset S7.** List of tomato accessions used to screen *RUP* gene polymorphism.

---

**Table S1.** Tomato accessions with SNPs in the *RUP* gene. A total of 391 tomato accessions were examined for the presence of SNPs in the *RUP* gene (**Dataset S7**). The EcoTILLING protocols used were as described in **Mohan et al. (2016).** The impact of SNPs on protein function was assessed using SIFT4G software **(Vasser et al., 2016)** and DDMut (**Zhou et al., 2023**).

| S.No | Accession number | Sequence validation | Amino acid change | SIFT prediction (SIFT Score) | DDMut prediction ($\Delta\Delta G^{\text{Stability wt} \rightarrow \text{mt}}$) | Effect/Nature of SNP |
| --- | --- | --- | --- | --- | --- | --- |
| 1 | Arka Vikas (AV) | Reference | - | - | - | - |
| 2 | EC398697<br>*rup-1* | C489CC | Frameshift | - | - | Protein truncation R164* |
|  |  | A513C | - | - | - | - |
|  |  | G531A | - | - | - | - |
|  |  | G567A | - | - | - | - |
|  |  | G731C | - | - | - | - |
|  |  | T787A | - | - | - | - |
| 3 | EC529085<br>*rup-2* | T456TT | Frameshift | - | - | Protein truncation R164* |
|  |  | G512GG | Frameshift | - | - | - |
|  |  | T688TT | Frameshift | - | - | - |
|  |  | C1023G | - | - | - | - |
|  |  | A1024T | - | - | - | - |
| 4 | EC2790<br>*rup-3* | G462C | S154= | - | -0.18 kcal/mol (Stabilizing) | Synonymous |
|  |  | G533A | S178N | Tolerated (0.96) |  | Nonsynonymous |
|  |  | C564A | T188= | - |  | Synonymous |
|  |  | G569A | G190E | Tolerated (1.0) |  | Nonsynonymous |
|  |  | C594T | V198= | - |  | Synonymous |
|  |  | T705C | H235= | - |  | Synonymous |
|  |  | G733C | V245L | Tolerated (0.89) |  | Nonsynonymous |
| 5 | EC398716<br>*rup-4* | T328A | L110I | Tolerated (0.65) | -0.42 kcal/mol (Destabilizing) | Nonsynonymous |
|  |  | G570A | G190= | - |  | Synonymous |

**Note:** SIFT score <0.05 is considered deleterious for the protein function.
**Mohan V, Gupta S, Thomas S, Mickey H, Charakana C, Chauhan VS, Sharma K, Kumar R, Tyagi K, Sarma S, Gupta SK, Kilambi HV, Nongmaithem S, Kumari A, Gupta P, Sreelakshmi Y, Sharma R** (2016) Tomato fruits show wide phenomic diversity but fruit developmental genes show low genomic diversity. *PLoS ONE* **11(4)**: e0152907.
**Vaser R, Adusumalli S, Leng SN, Sikic M, Ng PC.** (2016) SIFT missense predictions for genomes. *Nature Protocols*. **11**: 1–9.
**Zhou Y, Pan Q, Pires DE, Rodrigues CH, Ascher DB** (2023) DDMut: predicting effects of mutations on protein stability using deep learning. *Nucl. Acids Res.* **51**: W122–W128.

**Table S2.**
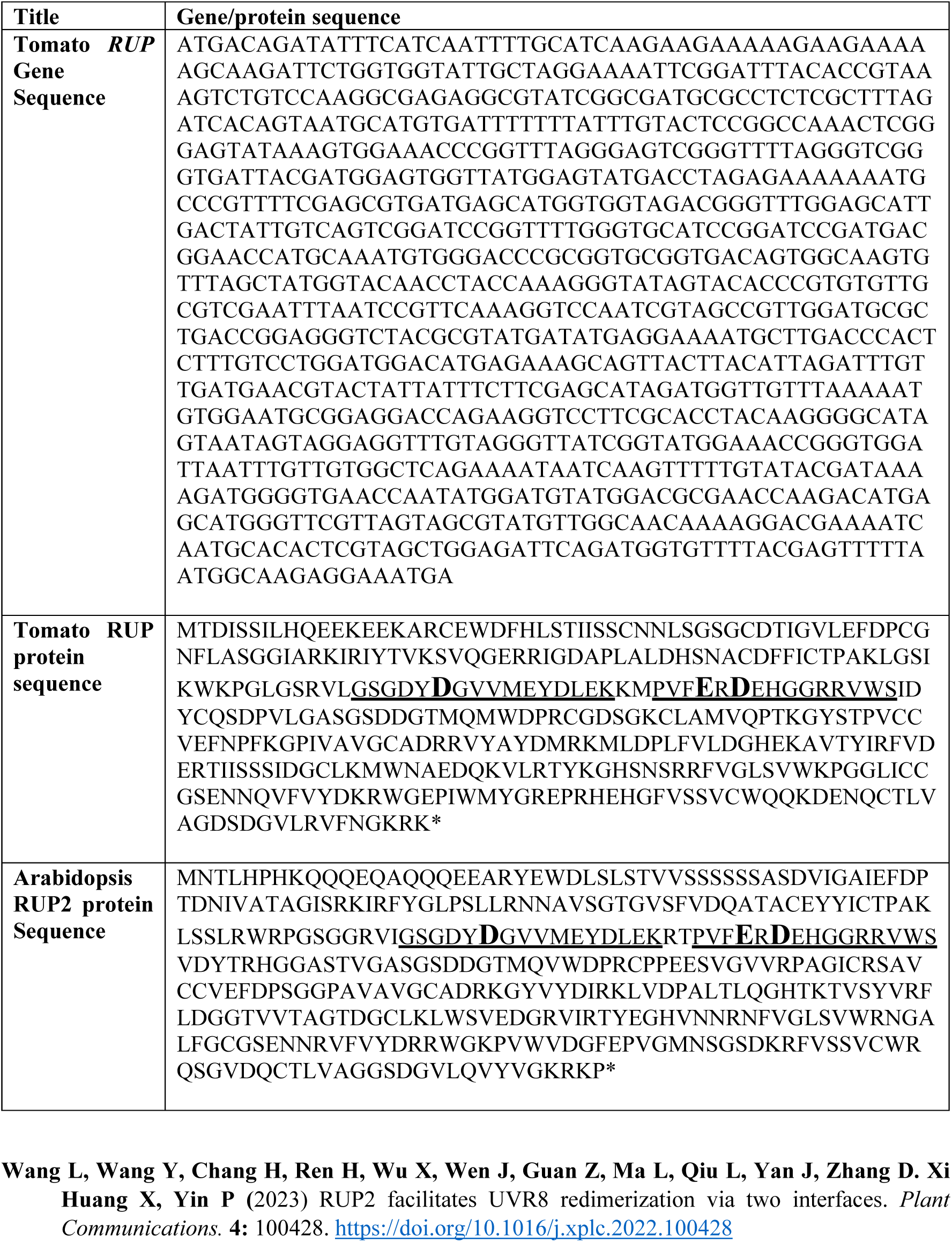
Tomato *RUP* gene and protein sequence. The residue D122, E138, and D140 of Arabidopsis RUP2 that plays a role in the interaction of RUP with UVR8 are highlighted in bold and bigger font (**Wang et al., 2023**). The sequences flanking above residues are underlined, to show the homology between tomato RUP and Arabidopsis RUP2 proteins.

**Table S3.**
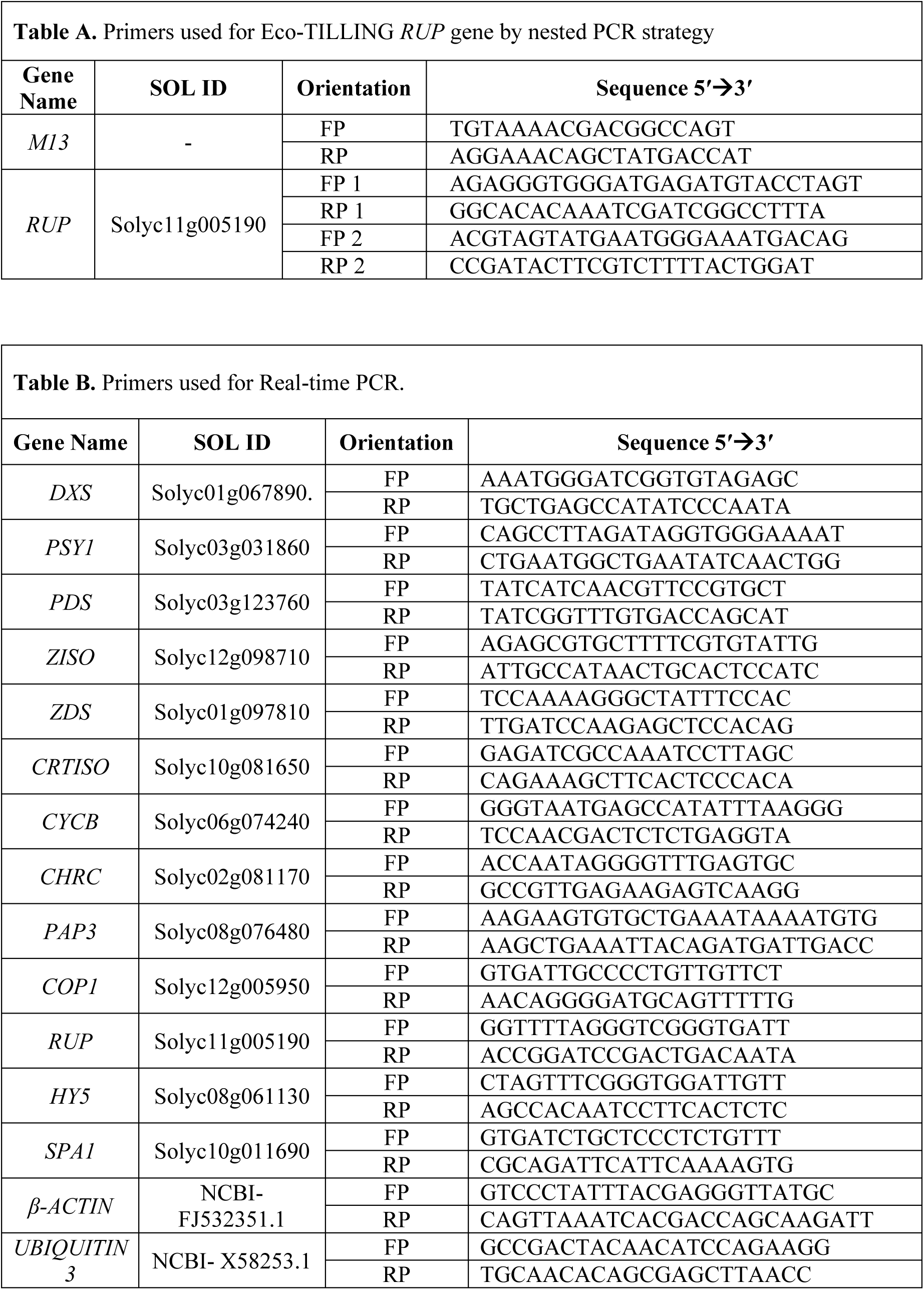
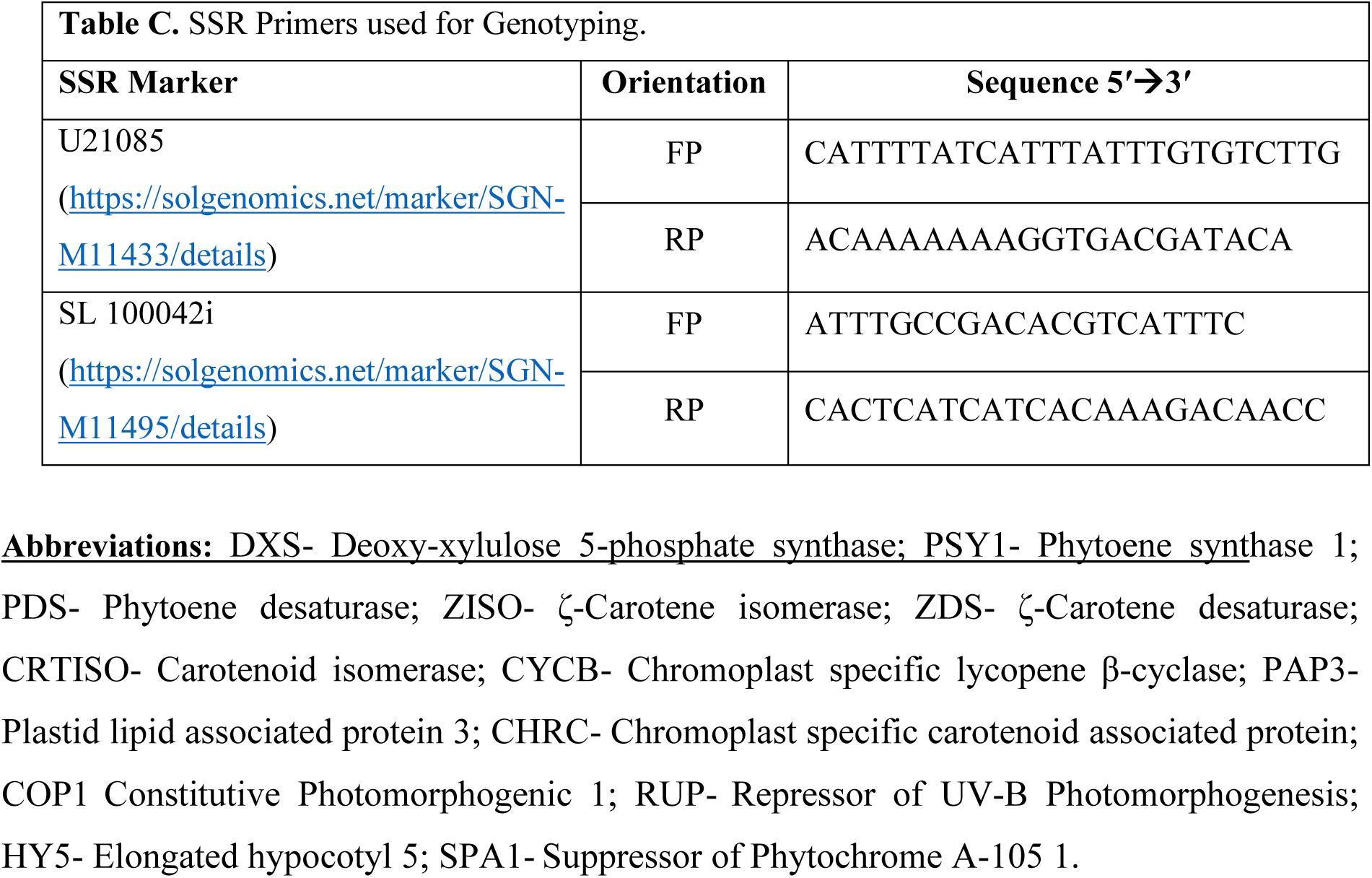
Primers used for EcoTILLING *RUP* gene and RT-PCR. **Table A.** Primers used for EcoTILLING. **Table B**. Primers used for RT-PCR. **Table C.** SSR Primers used for Genotyping.

## Notes

### Competing Interest Statement

The authors have declared no competing interest.

## References

Bianchetti RE, Lira SB, Monteiro SS, et al. (2018) Fruit-localized phytochromes regulate plastid biogenesis, starch synthesis, and carotenoid metabolism in tomato. J Exp Bot. 69: 3573–86.

Bodanapu R, Gupta SK, Basha PO, et al. (2016) Nitric oxide overproduction in tomato *shr* mutant shifts metabolic profiles and suppresses fruit growth and ripening. Front. Plant Sci. 7: 1714.

D’Andrea L, Rodriguez-Concepcion M. (2019) Manipulation of plastidial protein quality control components as a new strategy to improve carotenoid contents in tomato fruit. Front Plant Sci. 10: 1071.

de la Garza RID, Gregory III JF, Hanson AD. (2007) Folate biofortification of tomato fruit. Proc Nat Acad Sci. 104: 4218–22.

Deng H, Pirrello J, Chen Y, et al. (2018) A novel tomato F-box protein, SlEBF3, is involved in tuning ethylene signaling during plant development and climacteric fruit ripening. Plant J. 95: 648–658.

Dong H, Liu X, Zhang C, et al. (2021) Expression of Tomato UVR8 in Arabidopsis reveals conserved photoreceptor function. Plant Sci. 303: 110766.

Eckardt NA, Avin-Wittenberg T, Bassham DC, et al. (2024) The lowdown on breakdown: Open questions in plant proteolysis. Plant Cell. 36: 2931–75.

Fang F, Lin L, Zhang Q, et al. (2022) Mechanisms of UV-B light-induced photoreceptor UVR8 nuclear localization dynamics. New Phytol. 236: 1824–37.

Fantini E, Falcone G, Frusciante S, et al. (2013) Dissection of tomato lycopene biosynthesis through virus-induced gene silencing. Plant physiology. 163:986–98.

Fantini E, Sulli M, Zhang L, et al. (2019) Pivotal roles of cryptochromes 1a and 2 in tomato development and physiology. Plant Physiology. 179: 732–48.

Fernandez-Pozo N, Zheng Y, et al. (2017) The tomato expression atlas. Bioinformatics. 33: 2397–8.

Fraser PD, Romer S, Shipton CA, et al. (2002) Evaluation of transgenic tomato plants expressing an additional phytoene synthase in a fruit-specific manner. Proc Nat Acad Sci. 99: 1092–7.

Fray RG, Wallace A, Fraser PD, et al. (1995) Constitutive expression of a fruit phytoene synthase gene in transgenic tomatoes causes dwarfism by redirecting metabolites from the gibberellin pathway. Plant Journal. 8: 693–701.

Gruber H, Heijde M, Heller W, et al. (2010) Negative feedback regulation of UV-B–induced photomorphogenesis and stress acclimation in Arabidopsis. Proc Nat Acad Sci. 107: 20132–7.

Gupta P, Hirschberg J. (2022) The genetic components of a natural color palette: a comprehensive list of carotenoid pathway mutations in plants. Front. Plant Sci. 12: 806184.

Gupta P, Sreelakshmi Y, Sharma R (2015) A rapid and sensitive method for determination of carotenoids in plant tissues by high-performance liquid chromatography. Plant Methods 11:5

Gupta SK, Sharma S, Santisree P, et al., (2014) Complex and shifting interactions of phytochromes regulate fruit development in tomato. Plant Cell Environ. 37: 1688–1702.

Heijde M, Ulm R. (2013) Reversion of the Arabidopsis UV-B photoreceptor UVR8 to the homodimeric ground state. Proc Nat Acad Sci. 110: 1113–8.

Hetherington S, Smillie R, Davies W. (1998). Photosynthetic activities of vegetative and fruiting tissues of tomato. J Exp Bot 49: 1173–1181.

Hospital, F. (2003) Marker-assisted breeding. In: H.J. Newbury, editor. Plant Molecular Breeding. Oxford and Boca Raton: Blackwell Publishing and CRC Press; p. 30–59

Huang X, Yang P, Ouyang X, et al. (2014). Photoactivated UVR8-COP1 module determines photomorphogenic UV-B signaling output in Arabidopsis. PLoS Genetics. 10: e1004218.

Isaacson T, Ronen G, Zamir D, et al. (2002). Cloning of tangerine from tomato reveals a carotenoid isomerase essential for the production of β-carotene and xanthophylls in plants. Plant Cell. 14:333–42.

Jen JJ (1974). Spectral quality of light and the ripening characteristics of tomato fruit. Hortscience, 9: 548–549.

Kharshiing E, Sreelakshmi Y, Sharma R (2019) The Light Awakens! Sensing Light and Darkness. *In “*Plant Sensory Biology*”* Edited by SK Sopory, Springer Nature Singapore, pp 21–57

Kilambi HV, Dindu A, Sharma K, et al. (2021) The new kid on the block: A dominant-negative mutation of phototropin1 enhances carotenoid content in tomato fruits. Plant J. 106: 844-

Kilambi HV, Kumar R, Sharma R et al. (2013) Chromoplast-specific carotenoid-associated protein appears to be important for enhanced accumulation of carotenoids in hp1 tomato fruits. Plant Physiol. 161: 2085–2101.

Kilambi HV, Manda K, Sanivarapu H, et al. (2016) Shotgun Proteomics of Tomato Fruits: Evaluation, Optimization and Validation of Sample Preparation Methods and Mass Spectrometric Parameters. Front. Plant Sci. 7: 00969

Kim YE, Hipp MS, Bracher A, et al., (2013) Molecular chaperone functions in protein folding and proteostasis. Ann Rev Biochem. 82:323–55.

Li H, Li Y, Deng H, et al. (2018) Tomato UV-B receptor SlUVR8 mediates plant acclimation to UV-B radiation and enhances fruit chloroplast development via regulating SlGLK2. Scientific Reports. 17: 6097.

Liu B, Zuo Z, Liu H, et al. (2011). Arabidopsis cryptochrome 1 interacts with SPA1 to suppress COP1 activity in response to blue light. Genes Dev. 25:1029–34.

Liu CC, Ahammed GJ, Wang GT, et al. (2018) Tomato CRY1a plays a critical role in the regulation of phytohormone homeostasis, plant development, and carotenoid metabolism in fruits. Plant Cell Environ. 41:354–66.

Liu W, Jenkins GI. (2025) Recent advances in UV-B signalling: interaction of proteins with the UVR8 photoreceptor. J Exp Bot. 76: 873–881.

Liu X, Zhang Q, Yang G, et al. (2020). Pivotal roles of Tomato photoreceptor SlUVR8 in seedling development and UV-B stress tolerance. Biochem. Biophys. Res. Commun. 522: 177–83

Liu Y, Roof S, Ye Z, et al. (2004) Manipulation of light signal transduction as a means of modifying fruit nutritional quality in tomato. Proc Nat Acad Sci. 101: 9897–902.

McQuinn RP, Wong B, Giovannoni JJ. (2018) AtPDS overexpression in tomato: exposing unique patterns of carotenoid self-regulation and an alternative strategy for the enhancement of fruit carotenoid content. Plant Biotech J. 16: 482–94.

Mohan V, Gupta S, Thomas S, et al. (2016) Tomato fruits show wide phenomic diversity but fruit developmental genes show low genomic diversity. PLoS One 11: e0152907.

Neta-Sharir I, Isaacson T, Lurie S, et al., (2005) Dual role for tomato heat shock protein 21: protecting photosystem II from oxidative stress and promoting color changes during fruit maturation. Plant Cell. 17:1829–38.

Oravecz A, Baumann A, Máté Z, et al. (2006). CONSTITUTIVELY PHOTOMORPHOGENIC1 is required for the UV-B response in Arabidopsis. Plant Cell 18: 1975–1990.

Paik I, Huq E. (2019) Plant photoreceptors: Multi-functional sensory proteins and their signaling networks. Semi Cell Devl Bio. 92: 114–121.

Podolec R, Demarsy E, Ulm R. (2021) Perception and signaling of ultraviolet-B radiation in plants. Ann Rev Plant Biol. 72: 793–822.

Ponnu J, Hoecker U. (2021) Illuminating the COP1/SPA ubiquitin ligase: fresh insights into its structure and functions during plant photomorphogenesis. Front Plant Sci, 12: 662793.

Ren H, Han J, Yang P, et al. (2019) Two E3 ligases antagonistically regulate the UV-B response in Arabidopsis. Proc Nat Acad Sci. 116: 4722–31.

Sarma S, Pandey AK, Sharma K, et al., (2018) *MutS-Homolog2* silencing generates tetraploid meiocytes in tomato (*Solanum lycopersicum*). Plant Direct. 2: 1–15.

Sheerin DJ, Menon C, Oven-Krockhaus S, et al. (2015) Light-activated phytochrome A and B interact with members of the SPA family to promote photomorphogenesis in Arabidopsis by reorganizing the COP1/SPA complex. Plant Cell 27:189–201

Shinozaki Y, Nicolas P, Fernandez-Pozo N, et al., (2018) High-resolution spatiotemporal transcriptome mapping of tomato fruit development and ripening. Nature Communications 9: 364. | DOI: 10.1038/s41467-017-02782-9

Sim NL, Kumar P, Hu J, et al., (2012) SIFT web server: predicting effects of amino acid substitutions on proteins. Nucl Acid Res. 40(W1): W452–7.

Singh D, Mitra O, Mahapatra K, et al., (2024) REPRESSOR OF UV-B PHOTOMORPHOGENESIS proteins target ABSCISIC ACID INSENSITIVE 5 for degradation to promote early plant development. Plant Physiol. 196:2490–503.

Sreelakshmi Y, Gupta S, Bodanapu R, et al., (2010) NEATTILL: A simplified procedure for nucleic acid extraction from arrayed tissue for TILLING and other high-throughput reverse genetic applications. Plant Methods 6: 3.

Tanaka A, Fujita K, Kikuchi K. (1974) Nutrio-physiological studies on the tomato plant. III. Photosynthetic rate on individual leaves in relation to dry matter production of plants. Soil Sci Plant Nutr. 20: 173–183

Tissot N, Ulm R. (2020) Cryptochrome-mediated blue-light signalling modulates UVR8 photoreceptor activity and contributes to UV-B tolerance in Arabidopsis. Nature Commu. 11: 1323.

Tyagi K, Sunkum A, Gupta P, et al., (2023) Reduced γ-glutamyl hydrolase activity likely contributes to high folate levels in Periyakulam-1 tomato. *Horticulture Research*, Volume 10: uhac235.

Tyagi K, Sunkum A, Rai M, et al. (2022) Seeing the unseen: A trifoliate (MYB117) mutant allele fortifies folate and carotenoids in tomato fruits. Plant J. 112: 38–54.

Tyagi K, Upadhyaya P, Sarma S, et al. (2015) High performance liquid chromatography coupled to mass spectrometry for profiling and quantitative analysis of folate monoglutamates in tomato. Food Chemistry. 179: 76–84.

Upadhyaya P, Tyagi K, Sarma S, et al. (2016) Natural Variation in Folate Levels among Tomato (*Solanum lycopersicum*) Accessions. Food Chem. 217: 610–619.

Vizcaíno JA, Deutsch EW, Wang R, et al. (2014) ProteomeXchange provides globally coordinated proteomics data submission and dissemination. Nature Biotech. 32: 223–6.

Wang A, Chen D, Ma Q, et al. (2019) The tomato HIGH PIGMENT1/DAMAGED DNA BINDING PROTEIN 1 gene contributes to regulation of fruit ripening. Hort Res. 6: 15.

Wang HY, Ma LG, et al. (2001). Direct interaction of Arabidopsis cryptochromes with COP1 in light control development. Science. 294:154–8.

Wang L, Wang Y, Chang H, et al. (2023) RUP2 facilitates UVR8 redimerization via two interfaces. Plant Comm. 4: 100428.

Wang S, Liu J, Feng Y, et al. (2008). Altered plastid levels and potential for improved fruit nutrient content by downregulation of the tomato DDB1-interacting protein CUL4. The Plant J. 55: 89–103.

Wang W, Wang P, Li X, et al. (2021) The transcription factor SlHY5 regulates the ripening of tomato fruit at both the transcriptional and translational levels. Hort Res. 8: 83

Welsch R, Zhou X, Yuan H, et al. (2018). Clp protease and OR directly control the proteostasis of phytoene synthase, the crucial enzyme for carotenoid biosynthesis in Arabidopsis. Molecular Plant. 2018 11:149–62.

Yan C, Yang T, Wang B, et al. (2023) Genome-Wide Identification of the WD40 Gene Family in Tomato (*Solanum lycopersicum* L.). Genes.14: 1273.

Yang Y, Wu Y, Pirrello J, et al. (2010) Silencing Sl-EBF1 and Sl-EBF2 expression causes constitutive ethylene response phenotype, accelerated plant senescence, and fruit ripening in tomato. J Exp Bot. 61: 697–708.

Yin R, Arongaus AB, Binkert M, et al. (2015). Two distinct domains of the UVR8 photoreceptor interact with COP1 to initiate UV-B signaling in Arabidopsis. Plant Cell. 27: 202–13.

Zhang C, Zhang Q, Guo H, et al., (2021) Tomato SlRUP is a negative regulator of UV-B photomorphogenesis. Mol Hort. 1: 1–4.

Zhou Y, Pan Q, Pires DE, et al. (2023). DDMut: predicting effects of mutations on protein stability using deep learning. Nucleic Acids Research. 51(W1): W122–8.

Zhu F, Wen W, Cheng Y, et al. (2022) The metabolic changes that effect fruit quality during tomato fruit ripening. Mol Hort. 20:2.

